# Hydrogen sulfide blocks HIV rebound by maintaining mitochondrial bioenergetics and redox homeostasis

**DOI:** 10.1101/2021.04.21.440760

**Authors:** Virender Kumar Pal, Ragini Agrawal, Srabanti Rakshit, Pooja Shekar, Diwakar Tumkur Narasimha Murthy, Annapurna Vyakarnam, Amit Singh

## Abstract

A fundamental challenge in HIV eradication is to understand how the virus establishes latency, maintains stable cellular reservoirs, and promotes rebound upon interruption of antiretroviral treatment (ART). Here, we discovered an unexpected role of the ubiquitous gasotransmitter hydrogen sulfide (H_2_S) in HIV latency and reactivation. We show that reactivation of HIV-1 is associated with down-regulation of the key H_2_S producing enzyme cystathionine-*γ*-lyase (CTH) and reduction in endogenous H_2_S. Genetic silencing of CTH disrupts redox homeostasis, impairs mitochondrial function, and remodels the transcriptome of latent cells to trigger HIV reactivation. Chemical complementation of CTH activity using a slow-releasing H_2_S donor, GYY4137, suppressed HIV reactivation and diminished virus replication. Mechanistically, GYY4137 blocked HIV reactivation by inducing the Keap1-Nrf2 pathway, inhibiting NF-*κ*B, and recruiting the epigenetic silencer, YY1, to the HIV promoter. In latently infected CD4^+^ T cells from ART-suppressed human subjects, GYY4137 in combination with ART prevented viral rebound and improved mitochondrial bioenergetics. Moreover, prolonged exposure to GYY4137 exhibited no adverse influence on proviral content or CD4^+^ T cell subsets, indicating that diminished viral rebound is due to a loss of transcription rather than a selective loss of infected cells. In summary, this work provides mechanistic insight into H_2_S-mediated suppression of viral rebound and suggests the inclusion of an H_2_S donor in the current ART regimen to achieve a functional HIV-1 cure.

## Introduction

Human Immunodeficiency Virus (HIV), the causative agent of Acquired Immuno-Deficiency Syndrome (AIDS), is responsible for 0.6 million deaths and 1.7 million new infections in 2019 (https://www.who.int/gho/hiv/epidemic_status/deaths_text/en/). Despite advances in antiretroviral therapy (ART), the persistence of latent but replication-competent HIV in cellular reservoirs is a major barrier to cure (1). Understanding host factors and signaling pathways underlying HIV latency and rebound upon cessation of ART is of the highest importance in the search for an HIV cure (2). Host-generated gaseous signaling molecules such as nitric oxide (NO), carbon monoxide (CO), and hydrogen sulfide (H_2_S) modulate antimicrobial activity, inflammatory response, and metabolism of immune cells to influence bacterial and viral infections (3–8). Surprisingly, while circumstantial evidence links NO and CO with HIV-1 infection (9–11), the role of H_2_S remained completely unexplored.

H_2_S is primarily synthesized in mammals, including humans, via the reverse transsulfuration pathway using methionine and cysteine metabolizing enzymes cystathionine-gamma-lyase (CSE/CTH), cystathionine-beta-synthase (CBS), and cysteine-aminotransferase (CAT)-3-mercaptopyruvate sulfurtransferase (MPST), respectively (4). Because H_2_S is lipophilic, it rapidly permeates through biological membranes without membrane channels (lipid bilayer permeability, *P_M_ ³*0.5 ± 0.4 cm/s), dissociates into HS^−^ and S^2−^ in aqueous solution, and maintains an HS^−^: H_2_S ratio of 3:1 at physiological pH (4). Interestingly, expression and enzymatic activity of CTH was diminished in human CD8^+^ T cells and hepatic tissues derived from AIDS patients, respectively (12, 13). Additionally, the levels of H_2_S precursors L-cysteine and methionine were severely limited in AIDS patients (14). Also, HIV infection reduces the major cellular antioxidant glutathione (GSH) (15), whose production is dependent on L-cysteine and H_2_S (16, 17). Consequently, supplementation with N-acetyl cysteine (NAC), which is an exogenous source of L-cysteine, has been proposed for AIDS treatment (18). Since functional CBS and CTH enzymes are known to promote T cell activation and proliferation (19), endogenous levels of H_2_S can potentiate T cell-based immunity against HIV. Therefore, we think that the endogenous levels and biochemical activity of H_2_S, which critically depends upon the cysteine and transsulfuration pathway (20), are likely to be important determinants of HIV infection.

Calibrated production of H_2_S improves the functioning of various immune-cells by protecting from deleterious oxidants, stimulating mitochondrial oxidative phosphorylation (OXPHOS), and counteracting systemic inflammation (4, 21, 22). In contrast, supraphysiological concentrations of H_2_S exert cytotoxicity by inhibiting OXPHOS, promoting inflammation, and inducing redox imbalance (4, 21, 22). These H_2_S-specific physiological changes are also crucial for HIV infection. First, chronic inflammation contributes to HIV-1 pathogenesis by affecting innate and adaptive immune responses (23), impairing recovery of CD4^+^ T cells post-ART (24), increasing the risk of comorbidities (25, 26), and worsening organ function (27). Second, HIV-1 preferentially infects CD4^+^ T cells with high OXPHOS (28), relies on glycolysis for continuous replication (29), and reactivates in response to altered mitochondrial bioenergetics (30). Third, a higher capacity to resist oxidative stress promotes virus latency, whereas increased oxidative stress levels induce reactivation (31–33). Fourth, H_2_S directly (via S-linked sulfhydration) or indirectly modulates the activity of transcription factors NF-*κ*B and AP-1 (34, 35) that are well known to trigger HIV-1 reactivation from latency (36–38). Lastly, the thioredoxin/thioredoxin reductase (Trx/TrxR) system crucial for maintaining HIV-1 reservoirs (39) has been shown to regulate H_2_S- mediated S-persulfidation (40). Altogether, many physiological processes vital for HIV pathogenesis overlap with the underlying mechanism of H_2_S mediated-signaling.

Based on the role of H_2_S in regulating redox balance, mitochondrial bioenergetics, and inflammation, we hypothesize that intracellular levels of H_2_S modulate the HIV-1 latency and reactivation program. To test this hypothesis, we utilized cell line based models of HIV-1 latency and CD4^+^ T cells of HIV-infected patients on suppressive ART, which harbors the latent but replication-competent virus. Biochemical and genetic approaches were exploited to investigate the link between H_2_S and HIV-1 latency. Finally, we used NanoString gene expression technology, oxidant-specific fluorescent dyes, and real-time extracellular flux analysis to examine the role of H_2_S in mediating HIV-1 persistence by regulating gene-expression, redox signaling, and mitochondrial bioenergetics.

## Results

### Diminished biogenesis of endogenous H_2_S during HIV-1 reactivation

To investigate the link between HIV-1 latency and H_2_S, we measured changes in the expression of genes encoding H_2_S-generating enzymes CBS, CTH, and MPST in monocytic (U1) and lymphocytic (J1.1) models of HIV-1 latency (*Figure 1A*). The U1 and J1.1 cell lines are well-studied models of post-integration latency and were derived from chronically infected clones of a promonocytic (U937) and a T-lymphoid (Jurkat) cell line, respectively (41, 42). Both U1 and J1.1 show very low basal expression of the HIV-1 genome, which can be induced by several latency reversal agents (LRAs) such as PMA, TNF-*α*, and prostratin (*Figure 1B*) (41, 42). First, we confirmed virus reactivation by measuring HIV-1 *gag* transcript in U1 with a low concentration of PMA (5 ng/ml). Treatment of PMA induces detectable expression of *gag* at 12 h, which continued to increase for the entire 36 h duration of the experiment (*Figure 1C*). Next, we assessed the mRNA and protein levels of CBS, CTH, and MPST in U1 during virus latency (untreated) and reactivation (PMA-treated). The mRNA and protein expression levels of CTH showed a significant reduction at 24 and 36 h post-PMA treatment as compared to the untreated control, whereas the expression of CBS was not detected and MPST remained unaffected (*Figure 1D-E*). As an uninfected control, we measured the expression of CBS, CTH, and MPST in U937 cells. In contrast to U1, PMA treatment stimulated the mRNA and protein levels of CTH in U937 (*Figure 1F-G*), while CBS was barely detectable. Similar to U1 monocytic cells, PMA treatment reactivated HIV-1 and reduced the expression of CTH in J1.1 lymphocytic cells but not in the uninfected Jurkat cells in a time-dependent manner (*Figure 1*–*figure supplement 1A-B*). The expression of CBS was reduced in both J1.1 and Jurkat upon PMA treatment, indicating that only CTH specifically down-regulates in response to HIV-1 reactivation (*Figure 1–figure supplement 1A-B*). Finally, we directly measured endogenous H_2_S levels using a fluorescent probe 7-azido-4-methylcoumarin (AzMc) that quantitatively detects H_2_S in living cells (43). Consistent with the expression data, reactivation of HIV-1 by PMA reduced H_2_S generation in U1, whereas H_2_S levels were significantly increased by PMA in the uninfected U937 (*Figure 1H-I*). Taken together, these data indicate that HIV-1 reactivation is associated with diminished biogenesis of endogenous H_2_S.

**Figure 1.**
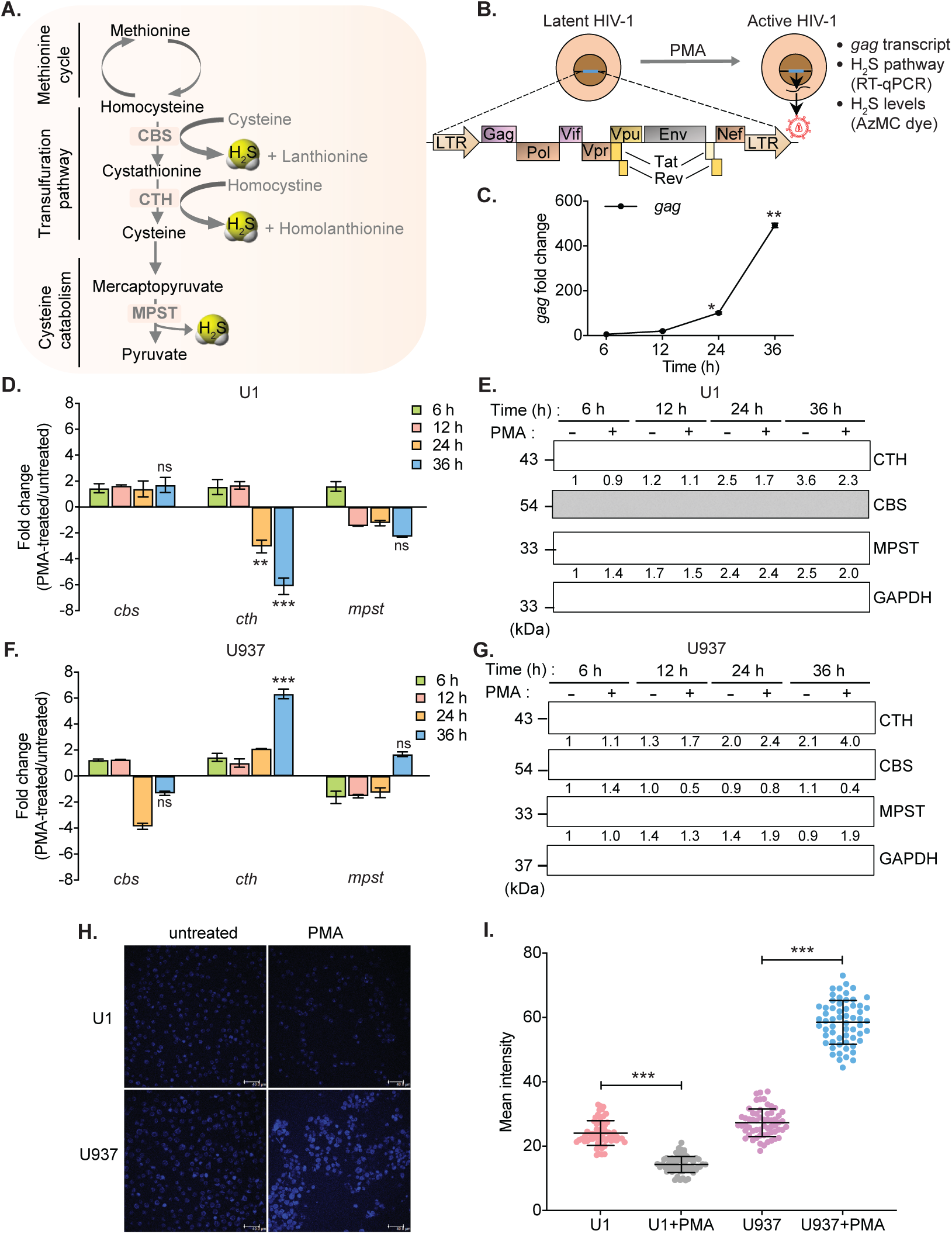
HIV reactivation diminishes expression of H_2_S metabolic enzymes and endogenous H_2_S levels. **(A)** Schematic showing H_2_S producing enzymes in mammalian cells. **(B)** Experimental strategy for measuring HIV-1 reactivation and H_2_S production in U1. **(C)** U1 cells were stimulated with 5 ng/ml PMA and the expression of *gag* transcript was measured at indicated time points. **(D and E)** Time-dependent changes in the expression of CTH, CBS and MPST at mRNA and protein level during HIV-1 latency (-PMA) and reactivation (+PMA [5 ng/ml]) in U1 cells. **(F and G)** Time-dependent changes in the expression of CTH, CBS and MPST at mRNA and protein level in U937 cells with or without PMA treatment. Results were quantified by densitometric analysis for CTH, CBS and MPST band intensities and normalized to GAPDH, using ImageJ software. **(H and I)** U1 and U937 cells treated with 5 ng/ml PMA for 24 h or left untreated, stained with AzMC for 30 min at 37°C, and images were acquired using Leica TCS SP5 confocal microscope (H). Scale bar represents 40 μm. Average fluorescence intensity was quantified by ImageJ software (I). Results are expressed as mean ± standard deviation and are representative of data from three independent experiments. *, P<0.05; **, P<0.01; ***, P<0.001, by two-way ANOVA with Tukey’s multiple comparison test. Figure 1 includes the following figure supplement: **Figure supplement 1.** HIV-1 reactivation decreases levels of H_2_S metabolizing enzymes in a T cell line model of latency.

### CTH-mediated reactivation of HIV-1 from latency

Our results suggest that H_2_S biogenesis via CTH is associated with HIV-1 latency. Therefore, we next asked whether CTH-derived H_2_S regulates reactivation of HIV-1 from latency. To test this idea, we depleted endogenous CTH levels in U1 using RNA interference (RNAi). The short hairpin RNA specific for CTH (U1-shCTH) silenced the expression of CTH mRNA and protein by 90 % as compared to non-targeting shRNA (U1-shNT) (*Figure 2A-B*). Moreover, using AzMC probe, we confirmed reduction in endogenous H_2_S levels in U1-shCTH as compared to U1-shNT (*Figure 2–figure supplement 1A-B)*. Next, we investigated the effect of H_2_S depletion via CTH suppression on HIV-1 latency by measuring viral transcription (*gag* transcript), translation (HIV-1 p24 capsid protein), and release (HIV-1 p24 abundance in the cell supernatant). We found that the low basal expression of HIV-1 *gag* in U1-shCTH was stimulated by 16-fold upon depletion of CTH (*p =* 0.0018) (*Figure 2C*). Furthermore, while PMA induced expression of *gag* transcript by 73-fold in U1-shNT, a further enhancement to 200-fold was observed in U1-shCTH (*Figure 2C*). Consistent with this, levels of p24 capsid protein inside cells or released in the supernatant were significantly elevated in U1-shCTH as compared to U1-shNT with (*p* = 0.028) or without PMA-treatment (*p* = 0.015) (*Figure 2D-E*). We also depleted CTH levels in J1.1 cells using RNAi and monitored HIV-1 reactivation by assessing intracellular p24 levels under basal conditions. The depletion of CTH triggered HIV-1 reactivation from latency as evident from a 7-fold increase in p24 levels as compared to shNT (*Figure 2–figure supplement 1C*). A minor increase in p24 levels (3.6-fold) was also apparent upon the depletion of CBS in J1.1 (*Figure 2–figure supplement 1C*). These data indicate that CTH and likely CTH-derived H_2_S supports latency and impedes the reactivation of HIV-1.

**Figure 2.**
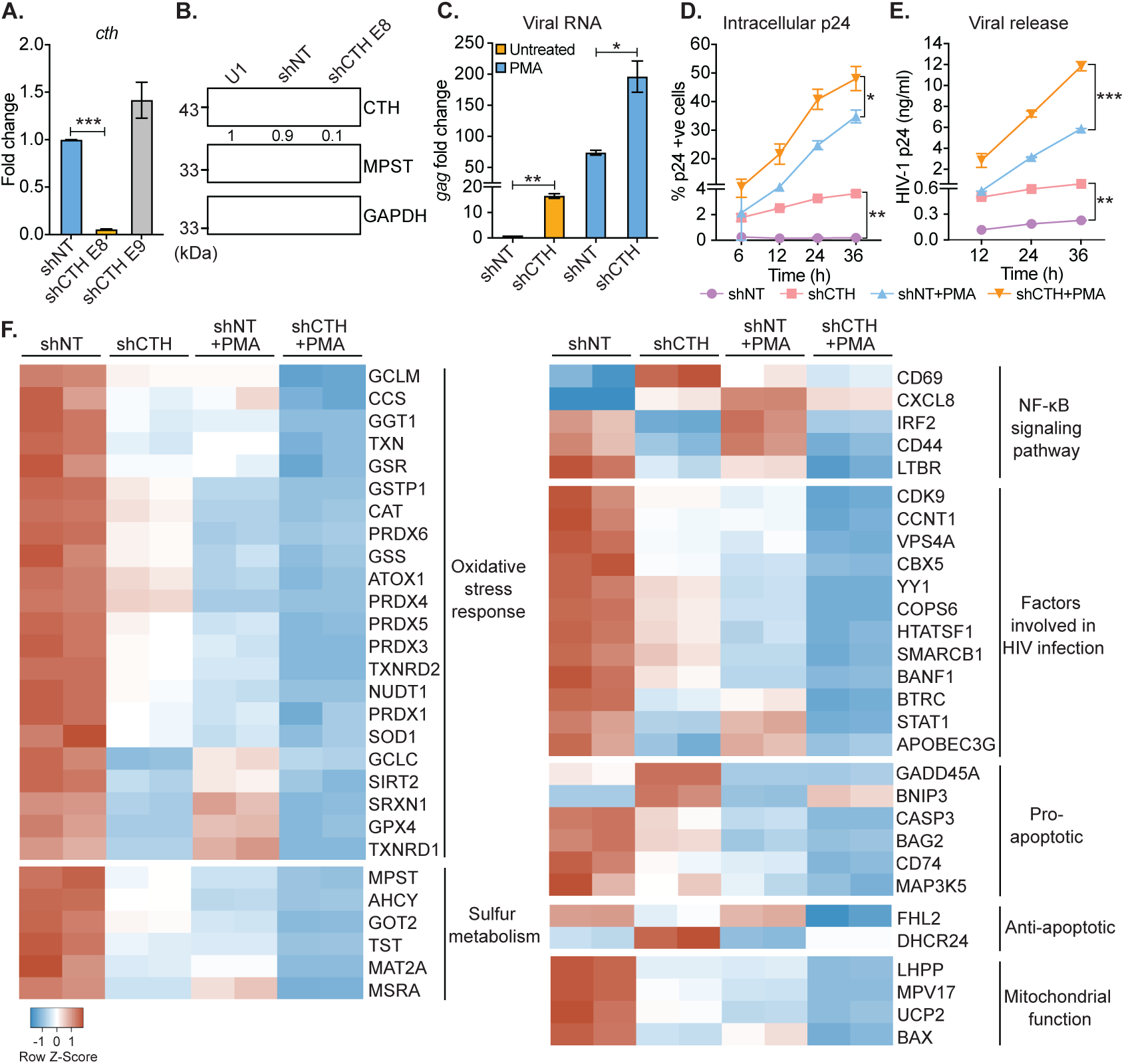
Genetic silencing of CTH reactivates HIV-1 and alters gene expression associated with redox stress, apoptosis, and mitochondrial function. **(A)** Total RNA was isolated from shCTH E8, shCTH E9, and non-targeting shRNA (shNT) lentiviral vectors transduced U1 cells and change in CTH mRNA was examined by RT-qPCR. **(B)** Cell lysates of U1, shNT, and shCTH E8 were assessed for CTH abundance using immuno-blotting. The CTH band intensities were quantified by densitometric analysis and normalized to GAPDH. **(C)** shCTH and shNT were treated with 5 ng/ml PMA or left untreated for 24 h and HIV-1 reactivation was determined by *gag* RT-qPCR. **(D and E)** shCTH and shNT were treated with 5 ng/ml PMA or left untreated. At the indicated time points, HIV-1 reactivation was measured by flow-cytometry using fluorescently tagged (PE-labeled) antibody specific to intracellular p24 (Gag) antigen and p24 ELISA in the supernatant. Results are expressed as mean ± standard deviation and are representative of data from two independent experiments. *, P<0.05; **, P<0.01; ***, P<0.001, by two-way ANOVA with Tukey’s multiple comparison test. **(F)** Total RNA isolated from untreated or PMA (5 ng/ml; 24 h)-treated U1-shNT and U1-shCTH were examined by NanoString Technology to assess the expression of genes associated with HIV infection and oxidative stress response. Heatmap showing functional categories of significantly differential expressed genes (DEGs). Gene expression data obtained were normalized to internal control β_2_ microglobulin (B2M), and fold changes were calculated using the nSolver 4.0 software. Genes with fold changes values of >1.5 and P <0.05 were considered as significantly altered. Figure 2 includes the following figure supplement: **Figure supplement 1.** Genetic silencing of CTH reduces endogenous H_2_S levels and reactivates HIV-1 from latency. **Source data file 1.** This file contains RAW values associated with the NanoString analysis of host genes affected upon depletion of CTH in U1.

### Altered expression of genes involved in HIV-1 reactivation upon CTH depletion

The above results indicate that CTH is a target controlling HIV-1 reactivation from latency, prompting us to investigate the mechanism. Several pathways that induce HIV-1 reactivation from latency are also influenced by H_2_S. These include redox signaling (44), NF-*κ*B pathway (45), inflammatory response (46), and mitochondrial bioenergetics (47). Hence, we conducted a focused expression profiling of 185 human genes intrinsically linked to HIV-1 reactivation using the NanoString nCounter technology (see *supplementary Table S2* for the gene list) to measure absolute amounts of multiple transcripts without reverse transcription. Because depletion of CTH promoted reactivation of the HIV-1 in U1, we compared the expression profile of U1-shCTH with U1-shNT to further understand the link between H_2_S biogenesis and HIV-1 latency. Examination of 84 genes related to oxidative stress response revealed differential regulation of 22 genes in U1-shCTH compared with U1-shNT (*Figure 2F* and *supplementary Table S3*). Interestingly, more than 90 % of genes showing altered expression were down-regulated in U1-shCTH. Of these, genes encoding key cellular antioxidant enzymes and buffers such as catalase (CAT), superoxide dismutase (SOD1), peroxiredoxin family, thioredoxin family (TXN, TXNRD1 and TXNRD2), sulfiredoxin (SRXN1), glutathione metabolism (GCLM, GSS, and GSR), and sulfur metabolism (MPST, AHCY, and MAT2A) were significantly less expressed in U1-shCTH as compared to U1-shNT (*Figure 2F*). Since oxidative stress elicits HIV-1 reactivation (31), these findings indicate that HIV-1 reactivation through CTH depletion could be a consequence of an altered redox balance. Further, pathways involved in promoting HIV-1 reactivation such as NF-*κ*B signaling (36) and apoptosis (48) were also induced in U1-shCTH as compared to U1-shNT (*Figure 2F*). Notably, multiple HIV-1 restriction factors important for viral latency including type I interferon signaling (IRF2, STAT-1) (49, 50), APOBEC3G (51), CDK9-CCNT1 (52, 53), SLP-1 (54), and chromatin remodelers (SMARCB1, BANF1, YY1) (55, 56) were down-regulated in U1-shCTH. HIV-1 proteins are known to target mitochondria to induce mitochondrial depolarization, elevate mitochondrial ROS, and apoptosis for replication (57). Several genes involved in sustaining mitochondrial function and membrane potential (*e.g.,* BAX, UCP2, LHPP, MPV17) were repressed in U1-shCTH (*Figure 2F*), highlighting a potential association between unrestricted virus replication, mitochondrial dysfunction, and CTH depletion.

Since PMA is known to reactivate HIV-1 (58), we examined the gene expression in U1-shNT upon reactivation of HIV-1 by PMA. We found that 80 % of genes affected by the depletion of CTH were similarly perturbed in response to PMA. Signature of transcripts associated with oxidative stress response, sulfur metabolism, anti-viral factors, and mitochondrial function was comparable in U1-shCTH and PMA-treated U1-shNT (*Figure 2F*). Based on these similarities, we hypothesized that combining PMA with CTH depletion would have an additive effect on the expression of genes linked to HIV-1 reactivation. Indeed, exposure of U1-shCTH to PMA induced gene expression changes which surpassed those produced by PMA treatment or CTH-depletion alone (*Figure 2F*). Notably, significant expression changes in U1-shCTH upon PMA treatment are consistent with our data showing maximal HIV-1 activation under these conditions (see *Figure 2C*). Overall, these results indicate that CTH contributes to HIV-1 latency by modulating multiple pathways coordinating cellular homeostasis (*e.g.,* redox balance, mitochondrial function) and the anti-viral response.

### CTH is required to maintain redox homeostasis and mitochondrial function

Physiological levels of H_2_S support redox balance and mitochondrial function by maintaining GSH balance (16), protecting against ROS (16), and acting as a substrate for the electron transport chain (21, 22). On this basis, we reasoned that the depletion of CTH could contribute to redox imbalance and mitochondrial dysfunction to promote HIV-1 reactivation. We first measured total glutathione content (GSH+GSSG) and GSSG concentration in U1-shNT and U1-shCTH using chemical-enzymatic analysis of whole-cell extract (31). Whole-cell glutathione content was not significantly different in U1-shNT and U1-shCTH (*p* = 0.2801) (*Figure 3A*). However, the GSSG concentration was elevated in U1-shCTH resulting in a concomitant decrease in the GSH/GSSG ratio of U1-shCTH as compared to U1-shNT (*p* = 0.0032) (*Figure 3A*). The increased GSSG pool and reduced GSH/GSSG poise confirm that cells are experiencing oxidative stress upon CTH depletion. We measured total ROS using a fluorescent probe, 5,6-chloromethyl-2*′*,7*′*-dicholrodihydrofluorescein diacetate (CM-H_2_DCFDA), which non-specifically responds to any type of ROS within the cells. Both U1-shNT and U1-shCTH showed comparable levels of cytoplasmic ROS (*Figure 3– figure supplement 1A*), which remained unchanged after stimulation with PMA. In addition to cytoplasmic ROS, we also measured mitochondrial ROS (mitoROS) using the red fluorescent dye MitoSOX, which selectively stains mitoROS. Lowering the levels of CTH severely increased mitoROS in U1, which was further accentuated after PMA stimulation (*Figure 3B*). We then studied the effect of CTH depletion on mitochondrial functions using a Seahorse XF Extracellular Flux Analyzer (Agilent) as described previously by us (*Figure 3C*) (30, 47). Both basal and ATP-coupled respiration was significantly decreased in U1-shCTH as compared to U1-shNT (*Figure 3D-E*), consistent with the role of endogenous H_2_S in reducing cytochrome C oxidase for respiration (47). The maximal respiratory capacity, attained by the dissipation of the mitochondrial proton gradient with the uncoupler FCCP, was markedly diminished in U1-shCTH (*Figure 3E*). The maximal respiration also facilitated the estimation of the spare respiratory capacity, which was nearly exhausted in U1-shCTH. Additional hallmarks of HIV-1 reactivation and dysfunctional mitochondria such as coupling efficiency and non-mitochondrial oxygen consumption rate (nmOCR) (30) were also adversely affected upon depletion of CTH (*Figure 3E*). These results indicate that CTH depletion decelerates respiration, diminishes the capacity of macrophages to maximally respire, and promotes nmOCR. All of these parameters are important features of mitochondrial health and are likely to be crucial for maintaining HIV-1 latency.

**Figure 3.**
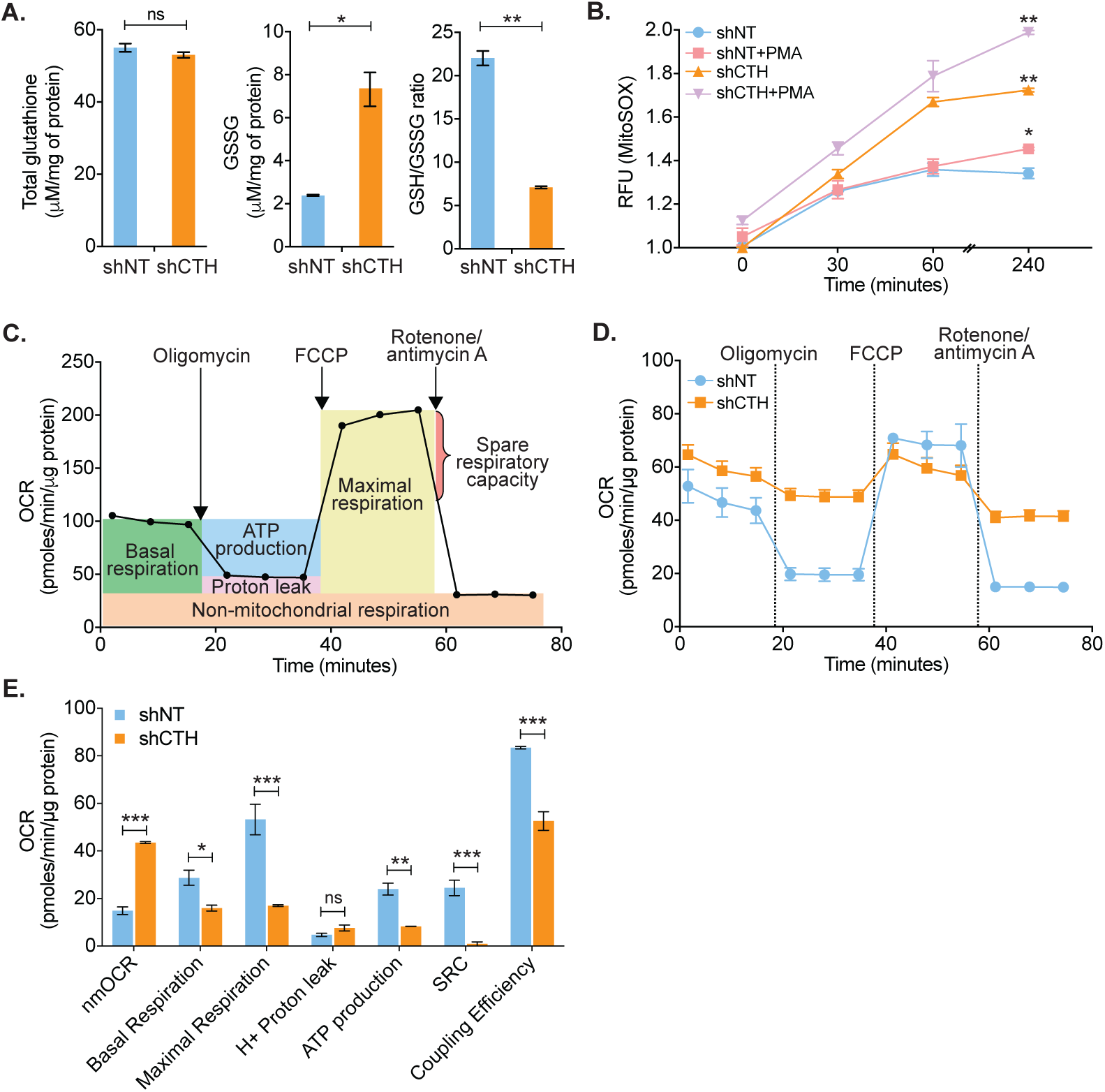
CTH maintains redox homeostasis and mitochondrial bioenergetics to promote HIV-1 latency. **(A)** Total and oxidized cellular glutathione (GSSG) content was assessed in U1-shCTH and U1-shNT cell lysates using glutathione assay kit. **(B)** U1-shNT and U1-shCTH were stained with MitoSOX*™* Red dye for 30 mins at 37°C and analyzed by flow cytometry (Ex-510 nm, Em-580 nm). **(C)** Schematic representation of Agilent Seahorse XF Cell Mito Stress test profile to assess key parameters related to mitochondrial respiration. **(D)** U1-shNT and U1-shCTH (5×10^4^) were seeded in triplicate wells of XF microplate and incubated for 1 h at 37°C in a non-CO_2_ incubator. Oxygen consumption was measured without adding any drug (basal respiration), followed by measurement of OCR change upon sequential addition of 1 μM oligomycin (ATP synthase inhibitor) and 0.25 μM carbonyl cyanide 4-(trifluoromethoxy) phenylhydrazone (FCCP), which uncouples mitochondrial respiration and maximizes OCR. Lastly, rotenone (0.5 μM) and antimycin A (0.5 μM) were injected to completely inhibit respiration by blocking complex I and complex III, respectively. **(E)** Various respiratory parameters derived from OCR measurement were determined by Wave desktop software. nmOCR; non-mitochondrial oxygen consumption rate and SRC; spare respiratory capacity. Error bar represent standard deviations from mean. Results are representative of data from three independent experiments. *, P<0.05; **, P<0.01; ***,P<0.001; ns, nonsignificant, by two-way ANOVA with Bonferroni’s multiple comparison test. Figure 3 includes the following figure supplement: **Figure supplement 1.** Effect of CTH knockdown on cytosolic ROS generation.

### A small-molecule H_2_S donor diminished HIV-1 reactivation and viral replication

One of the aims of the HIV-1 functional cure is to identify small molecules that can be combined with ART to delay or halt virus replication, reactivation, and replenishment of the latent reservoir (59, 60). Having shown that diminished levels of endogenous H_2_S is associated with HIV-1 reactivation, we next examined if elevating H_2_S levels using a small-molecule H_2_S donor-GYY4137 sustains HIV-1 latency. The GYY4137 is a widely used H_2_S donor that releases a low amount of H_2_S over a prolonged period to mimic physiological production (61). Because H_2_S discharge is unusually sluggish by GYY4137, the final concentration of H_2_S released is likely to be significantly lower than the initial concentration of GYY4137. We confirmed this by measuring H_2_S release in U1 cells treated with low (0.3 mM) or high (5 mM) concentrations of GYY4137 using methylene blue colorimetric assay. As a control, we treated U1 cells with 0.3 mM NaHS, which rapidly releases a high amount of H_2_S. As expected, NaHS treatment rapidly released 90 % of H_2_S within 5 min of treatment (*Figure 4A*). In contrast, GYY4137 uniformly released a low amount of H_2_S for the entire 120 h duration of the experiment (*Figure 4A*).

**Figure 4.**
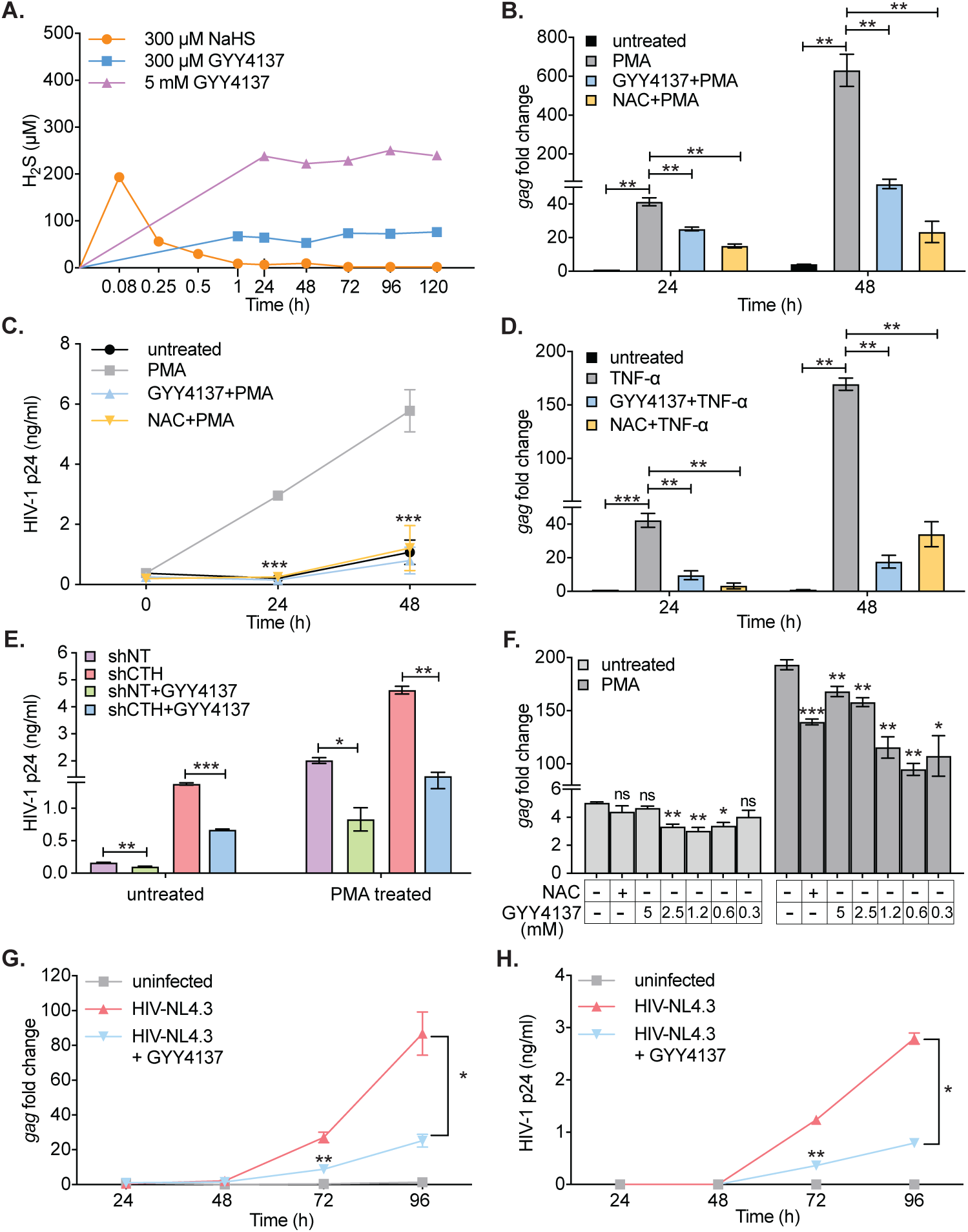
H_2_S donor (GYY4137) suppresses HIV-1 reactivation and replication. **(A)** U1 cells were treated with NaHS or GYY4137 and media supernatant was harvested to assess H_2_S production by methylene blue assay over time. **(B and C)** U1 cells were pre-treated with 5 mM GYY4137 or 5 mM NAC for 24 h and then stimulated with 5 ng/ml PMA for 24 h and 48 h. Total RNA was isolated and HIV-1 reactivation assessed by *gag* RT-qPCR (B). Culture supernatant was harvested to monitor HIV-1 release by p24 ELISA (C). **(D)** U1 cells were pretreated with 5 mM GYY417, 5 mM NAC for 24 h or left untreated and then stimulated with 100 ng/ml TNF-ɑ for 24 h and 48 h. HIV-1 reactivation was assessed by *gag* RT-qPCR. **(E)** U1-shCTH and U1-shNT were pretreated with 5 mM GYY4137 for 24 h and stimulated with 5 ng/ml PMA for 24 h. Culture supernatant was harvested to determine HIV-1 reactivation by HIV-1 p24 ELISA. **(F)** J1.1 cells were pretreated with indicated concentrations of GYY4137 or 5 mM NAC for 24 h and then stimulated with 5 ng/ml PMA for 12 h. Cells were harvested to isolate total RNA and HIV-1 reactivation was assessed by *gag* RT-qPCR. **(G and H)** Jurkat cells were infected with HIV-NL4.3 at 0.2 MOI for 4 h, cells were washed, seeded in fresh media, and cultured in presence or absence of 300 μM GYY4137. HIV-1 replication was monitored by measuring intracellular *gag* transcript levels and p24 protein levels in culture supernatant by ELISA. Results are expressed as mean ± standard deviation and data are representative of three independent experiments. *, P<0.05; **, P<0.01; ***, P<0.001, by two-way ANOVA with Tukey’s multiple comparison test. Figure 4 includes the following figure supplement: **Figure supplement 1.** Effect of GYY4137 on HIV-1 reactivation and cellular viability.

We systematically tested the effect of GYY4137 on HIV-1 reactivation using multiple models of HIV-1 latency and replication. As a control, we used N-acetyl cysteine (NAC) that is known to block HIV-1 reactivation (62). Pretreatment of U1 with a non-toxic dose of GYY4137 (5 mM) (*Figure 4–figure supplement 1A*) diminished the expression of *gag* transcript by 2-fold at 24 h and 10-fold at 48 h post-PMA treatment (*Figure 4B*). The effect of GYY4137 on p24 levels in the supernatant was even more striking as it completely abolished the time-dependent increase in p24 concentration post-PMA treatment (*Figure 4C*). As expected, pretreatment with NAC similarly prevented PMA-triggered reactivation of HIV-1 in U1 (*Figure 4C*). Pretreatment of U1 with spent GYY4137, which comprises the decomposed backbone, showed no effect on PMA induced *gag* transcript (*Figure 4–figure supplement 1B*). Because TNF-*α* is a physiologically relevant cytokine that reactivates HIV-1 from latency (41), we tested the effect of GYY4137 on TNF-*α*-mediated virus reactivation. Treatment of U1 with TNF-*α* stimulated the expression of *gag* transcript by 42- and 169-fold at 24 and 48 h, post-treatment, respectively (*Figure 4D*). The addition of GYY4137 or NAC nearly abolished the reactivation of HIV-1 in response to TNF-*α* treatment (*Figure 4D*). Earlier, we showed that depletion of CTH stimulated HIV-1 reactivation in U1 (see *Figure 2C*). Therefore, we tested if GYY4137 could complement this genetic deficiency and subvert HIV-1 reactivation. The U1-shCTH cells were pre-treated with 5 mM GYY4137 and p24 levels in the supernatant were measured. The elevated levels of p24 in U1-shCTH were reduced by 2-fold under basal conditions and 3.2-fold upon PMA stimulation in response to GYY4137 (*Figure 4E*). Both RNAi and chemical complementation data provide evidence that H_2_S is one of the factors regulating HIV-1 latency and reactivation in U1.

Similar to U1, we next examined whether GYY4137 subverts HIV-1 reactivation in the J1.1 T-cell line model. Treatment of J1.1 with PMA for 12 h resulted in a 38-fold increase in *gag* transcript as compared to untreated J1.1 (*Figure 4F*), indicating efficient reactivation of HIV-1. Importantly, pretreatment with various non-toxic concentrations of GYY4137 (*Figure 4–figure supplement 1C*) reduced the stimulation of *gag* transcription by 2-fold upon subsequent exposure to PMA (*Figure 4F*). The inhibitory effect of GYY4137 on HIV-1 reactivation was relatively greater compared to NAC in J1.1 (*Figure 4F*). We used another well-established T cell-based model of HIV-1 latency (J-Lat) to examine the influence of H_2_S. In J-Lat, the integrated HIV-1 genome encodes green-fluorescent protein (GFP), which allows precise quantification of HIV-1 reactivation from latency in response to LRAs such as PMA, TNF-*α*, and prostratin (63). Consistent with this, treatment of J-Lat with 5 μM prostratin for 24 h induced significant HIV-1 reactivation, which was translated as 100 % increase in GFP^+^ cells (*Figure 4–figure supplement 1D*). Pretreatment with GYY4137 significantly reduced HIV-1 reactivation in a dose-dependent manner (*Figure 4–figure supplement 1D*).

We also examined if an endogenous increase in H_2_S by GYY4137 halts the replication of HIV-1. To this end, we infected CD4^+^ T cell (Jurkat) with CXCR4-using HIV-1 (pNL4.3) and monitored HIV-1 replication by measuring the expression of *gag* transcript and the levels of p24 in the supernatant. A progressive increase in *gag* transcription and p24 release was detected for the entire 96 h duration of the experiment, indicating efficient viral replication (*Figure 4G-H*). Pretreatment with a non-toxic concentration of GYY4137 (0.3 mM) (*Figure 4–figure supplement 1E*) resulted in a 3 to 3.4-fold inhibition of HIV-1 replication at 72 and 96 h post-infection (*Figure 4G-H*). Overall, these data establish that elevated levels of endogenous H_2_S efficiently suppress HIV-1 reactivation and replication.

### GYY4137 reduced the expression of host genes involved in HIV-1 reactivation

To dissect the mechanism of GYY4137-mediated inhibition on HIV-1 reactivation, we examined the expression of 185 genes associated with HIV-1 reactivation using the NanoString nCounter technology as described above (*Figure 5A* and *supplementary Table S4*). Expression was analyzed for viral latency (unstimulated U1), reactivation (PMA-stimulated treated U1; PMA), and H_2_S-mediated suppression (U1-GYY+PMA). Consistent with the role of PMA-mediated oxidative stress in preceding HIV-1 reactivation (31), expression of genes encoding ROS, and RNS generating enzymes (e.g., NADPH oxidase [NCF1, CYBB] and Nitric oxide synthase [NOS2]) were up-regulated (*Figure 5A*). Also, the expression of major antioxidant enzymes (*e.g.,* GPXs, PRDXs, and CAT) and redox buffers (GSH and TRX pathways) remained repressed in U1-PMA, indicative of elevated oxidative stress. Additionally, pro-inflammatory signatures (*e.g.,* TGFB1 and SERPINA1) and trans-activators (*e.g.,* FOS) were induced, whereas host factors involved in HIV-1 restriction (*e.g.,* IRF1 and YY1) were repressed in U1-PMA as compared to U1. In agreement with the reduction in endogenous H_2_S levels during HIV-1 reactivation, expression of CTH involved in H_2_S anabolism was down-regulated and H_2_S catabolism (SQRDL) was up-regulated by PMA.

**Figure 5.**
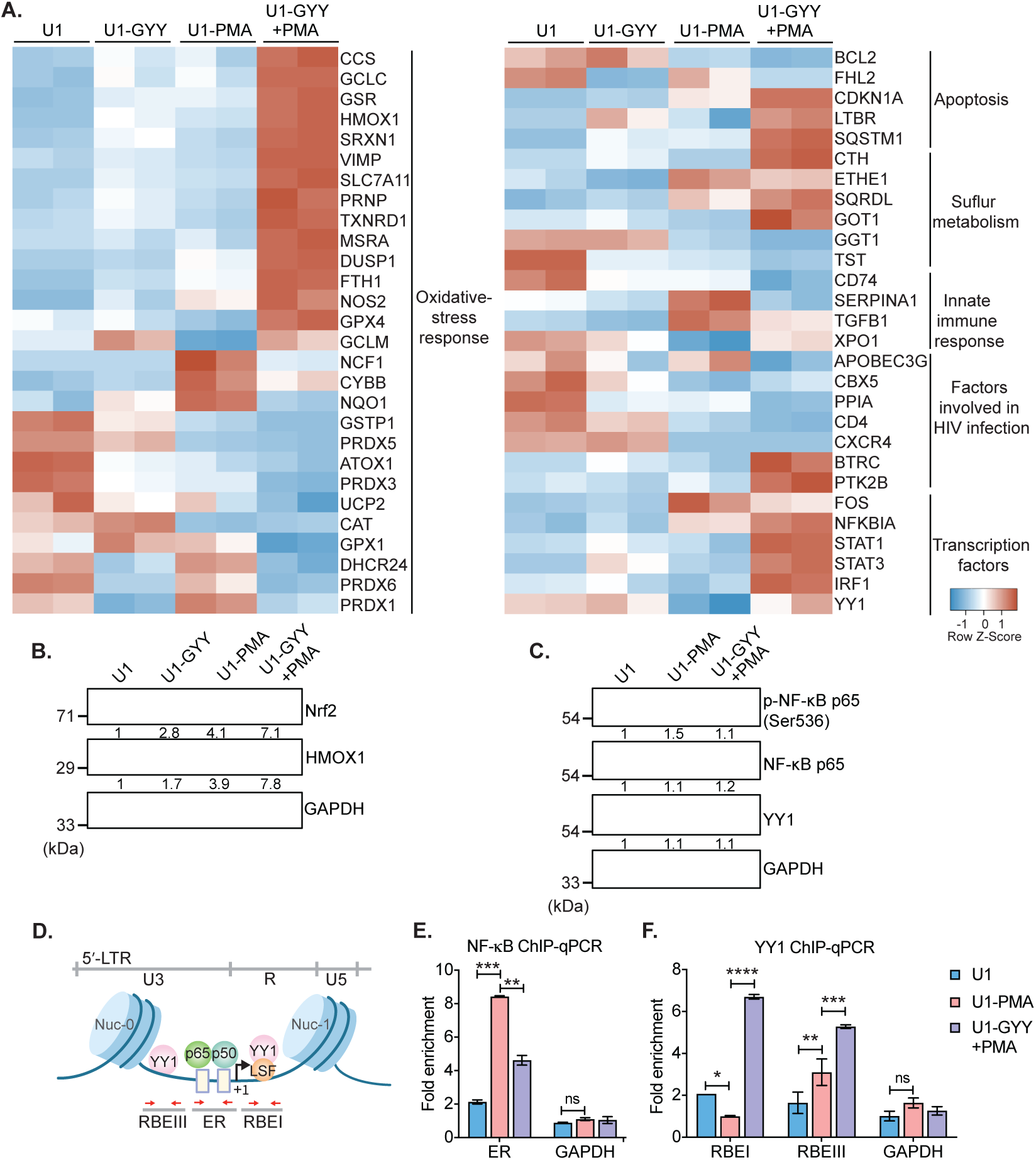
GYY4137 modulates Nrf2, NF-κB, and YY1 pathways. **(A)** U1 cells pre-treated with 5 mM GYY4137 for 24 h or left untreated and then stimulated with 5 ng/ml PMA for 24 h or left unstimulated. Total RNA was isolated and expression of genes associated with HIV infection and oxidative stress response was assessed by nCounter NanoString technology. Heatmap showing functional categories of significant DEGs in all four conditions: untreated (U1), GYY4137 alone (U1-GYY) or PMA-alone (U1-PMA) and GYY4137 +PMA (U1-GYY+PMA). Gene expression data obtained were normalized to internal control β_2_ microglobulin (B2M), and fold changes were calculated using the nSolver 4.0 software. Genes with fold changes values of >1.5 and P <0.05 were considered as significantly altered. **(B)** Total cell lysates were used to analyze the expression levels of Nrf2 and HMOX1 by immuno-blotting. Results were quantified by densitometric analysis of Nrf2 and HMOX1 band intensities and normalized to GAPDH. **(C)** U1 cells were pretreated with 5 mM GYY4137 for 6 h and then stimulated with 30 ng/ml PMA for 4 h or left unstimulated. Cells were harvested to prepare total cell lysate. Levels of phosphorylated NF-κB p65 (Ser536), NF-κB p65, and YY1 were determined by immuno-blotting. Results were quantified by densitometric analysis for each blot and were normalized to GAPDH. **(D)** Schematic depiction of the binding sites for NF-κB (p65-p50 heterodimer) and YY1 on HIV-1 5′-LTR. Highlighted arrow in red indicates the regions targeted for genomic qPCR; ER site for NF-κB, and RBEI and RBEIII sites for YY1 enrichments, respectively. **(E and F)** U1 cells were pretreated with 5 mM GYY4137 for 6 h, stimulated with PMA (30 ng/ml) for 4 h, fixed with formaldehyde, and lysed. Lysates were subjected to immunoprecipitation for p65 and YY1 and protein-DNA complexes were purified using protein-G magnetic beads. The enrichment of NF-κB p65 and YY1 on HIV-1 LTR was assessed by qPCR for designated regions using purified DNA as a template. Results are expressed as mean ± standard deviation and data are representative of three independent experiments *, P<0.05; **, P<0.01; ***, P<0.001; ****, P<0.0001; ns, nonsignificant, by two-way ANOVA with Tukey’s multiple comparison test. Figure 5 includes the following figure supplement: **Source data file 1.** This file contains RAW values associated with the NanoString analysis of host genes affected upon PMA induced HIV reactivation in presence or absence of GYY4137 treatment.

We noticed that the treatment with GYY4137 reversed the effect of PMA on the expression of genes associated with oxidative stress, inflammation, anti-viral response, apoptosis, and trans-activators (*Figure 5A*). For example, GYY4137 elicited a robust induction of genes regulated by nuclear factor erythroid 2-related factor 2 (Nrf2) in U1-GYY+PMA compared to U1 or U1-PMA (*Figure 5A*). Nrf2 acts as a master regulator of redox metabolism (64) by binding to the antioxidant response element (ARE) and initiating transcription of major antioxidant genes (*e.g.,* GSH pathway, TXNRD1, HMOX1, CTH, GPX4, and SRXN1). The Nrf2 activity has been shown to pause HIV-1 infection by inhibiting the insertion of reverse-transcribed viral cDNA into the host chromosome (65). Furthermore, sustained activation of Nrf2-dependent antioxidant response is essential for the establishment of viral latency (33). A few genes encoding H_2_O_2_ detoxifying enzymes (*e.g.,* CAT, GPX1, and PRDXs) were repressed in U1-GYY+PMA, indicating that the expression of these enzymes was likely counterbalanced by the elevated expression of other antioxidant systems by H_2_S. In line with the antagonistic effect of GYY4137 on HIV-1 transcription, the expression of HIV-1 trans-activator (FOS) was down-regulated, and an inhibitor of NF-*κ*B signaling (NFKBIA) was induced in U1-GYY+PMA as compared to U1 or U1-PMA. Lastly, GYY4137 stimulated the expression of several anti-HIV (YY1, STAT1, STAT3, and IRF1) and pro-survival factors (CDKN1A and LTBR) that were repressed by PMA (*Figure 5A*). Altogether, H_2_S supplementation induces a major realignment of redox metabolism and immune pathways associated with HIV-1 reactivation.

### GYY4137-mediated modulation of the Keap1-Nrf2 axis, NF-κB signaling, and activity of epigenetic factor YY1

Our expression data indicate activation of the Nrf2 pathway and modulation of transcription factors such as NF-*κ*B and YY1 upon treatment with GYY4137. We tested if the mechanism of H_2_S-mediated subversion of HIV-1 reactivation involves these pathways. H_2_S has recently been shown to prevent cellular senescence by activation of Nrf2 via S-persulfidation of its negative regulator Keap-1 (66). Under unstimulated conditions, Nrf2 binds to Keap1, and the latter promotes Nrf2 degradation via the proteasomal machinery (66). Nrf2 disassociates from the S-persulfidated form of Keap1, accumulates in the cytoplasm, and translocates to nuclei where it induces transcription of antioxidant genes upon oxidative stress (66). We first examined if GYY4137 treatment accumulates Nrf2 in the cytoplasm. As expected, Nrf2 was not detected in U1 owing to its association with Keap1 under unstimulated conditions. However, a noticeable accumulation of Nrf2 was observed in U1-GYY+PMA compared to U1 or U1-PMA (*Figure 5B*). As an additional verification, we quantified the levels of an Nrf2-dependent protein HMOX-1. Similar to Nrf2, the levels of HMOX-1 were also induced in U1-GYY+PMA (*Figure 5B*). These findings are consistent with our NanoString data showing activation of Nrf2-specific oxidative stress responsive genes in GYY4137 treated U1.

Next, we determined if GYY4137 treatment targets NF-*κ*B, which is a major regulator of HIV-1 reactivation (36). The cellular level of phosphorylated serine-536 in p65, a major subunit of NF-*κ*B, is commonly measured to assess NF-*κ*B activation (67). Since PMA reactivates HIV-1, we observed an increase in p65 ser-536 phosphorylation in U1-PMA (*Figure 5C*). In contrast, pretreatment with GYY4137 significantly decreased PMA-induced p65 ser-536 phosphorylation (*Figure 5C*). These findings suggest that GYY4137 is likely to affect the DNA binding and transcriptional activity of NF-*κ*B. Consistent with this idea, a two-step chromatin immunoprecipitation and genomic qPCR (ChIP-qPCR) assay showed that GYY4137-treatment significantly reduced the occupancy of p65 at its binding site (ER, enhancer region) on the HIV-1 LTR (*Figure 5D-E*).

GYY4137 induces the expression of another transcription factor YY1, which binds to HIV-1 LTR at RBEI and RBEIII and recruits histone deacetylase (HDACs) to facilitate repressive chromatin modifications (56). Overexpression of YY1 is known to promote HIV-1 latency (68). We tested if GYY4137 promotes the binding of YY1 at RBEI and RBEIII sites on HIV-1 LTR by ChIP-qPCR. The occupancy of YY1 at RBEI and RBEIII was significantly enriched in the case of U1-GYY+PMA as compared to U1-PMA or U1 (*Figure 5F*). Taken together, these results indicate that increasing endogenous H_2_S levels by using a slow-releasing donor effectively modulate the Keap1-Nrf2 axis and activity of transcription factors to maintain redox balance and control HIV-1 reactivation.

### GYY4137 blocks HIV-1 rebound from latent CD4^+^ T cells isolated from infected individuals on suppressive ART

Having shown that H_2_S suppresses HIV-1 reactivation in multiple cell line models of latency, we next studied the ability of GYY4137 to limit virus transcription in primary CD4**^+^** T cells derived from virally suppressed patients. We used a previously established methodology of maintaining CD4**^+^** T cells from infected individuals on ART for a few weeks without any loss of phenotypic characteristics associated with HIV reservoirs (60). We isolated CD4**^+^** T cells from peripheral blood mononuclear cells (PBMCs) of five HIV-infected subjects on suppressive ART. The CD4**^+^** T cells were expanded in the presence of interleukin-2 (IL-2), phytohemagglutinin (PHA), and feeder cells (*Figure 6A*). The expanded cells were cultured for 4 weeks in a medium containing IL-2 with ART (100 nM efavirenz, 180 nM zidovudine, and 200 nM raltegravir) either in the presence or absence of GYY4137 (100 μM) (*Figure 6A*). In this model, the virus reactivates by day 14 followed by progressive suppression at day 21 and latency establishment by day 35. The virus can be reactivated from this latent phase using well-known LRAs (60). We quantified the temporal expression of viral RNA using RT-qPCR periodically at 7 days interval. The levels of viral RNA increased initially (day 7 to 14), followed by low to undetectable levels by day 28 (*Figure 6B*). By day 28, virus RNA was uniformly untraceable in cells derived from GYY4137 treated and untreated groups, indicating the establishment of latency (*Figure 6B*).

**Figure 6.**
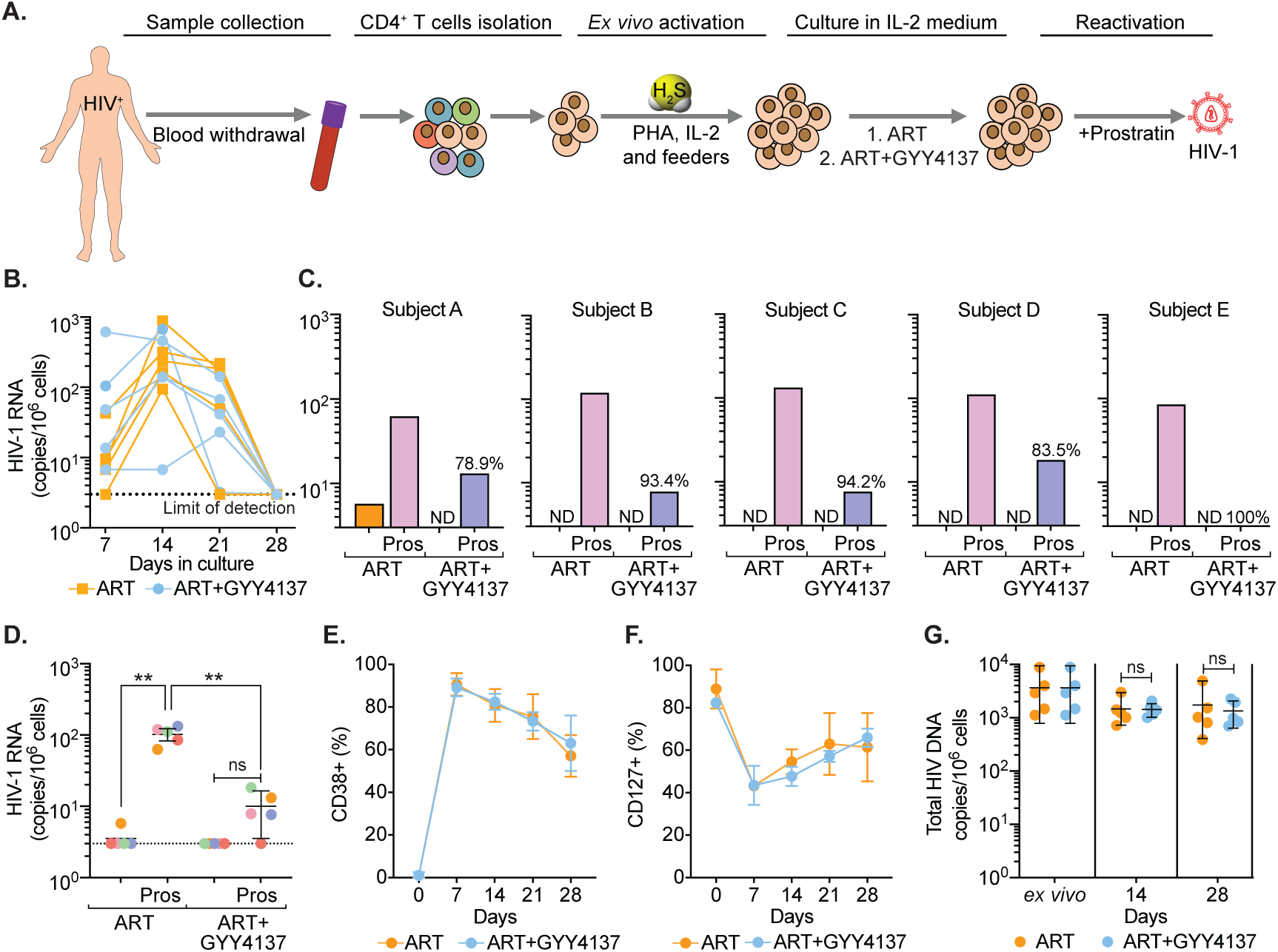
GYY4137 subverts HIV-1 reactivation in latent CD4^+^ T cells derived from HIV-1 infected patients. **(A)** Schematic representation of PBMCs extraction from blood samples of ART-treated HIV-1 infected subjects. CD4^+^ T cells were sorted and activated with PHA (1 μg/ml), IL-2 (100 U/ml), feeder PBMCs (gamma-irradiated) from healthy donor. CD4^+^ T cells were activated in presence of ART or ART in combination with 100 μM GYY4137. Post-activation cells were culture with ART alone or ART plus GYY4137 treatment in IL-2 containing medium. **(B)** Total RNA was isolated from five patients CD4^+^ T cells expanded in ART or ART plus GYY4137. HIV-1 RNA levels were measured every 7 days by ultrasensitive semi-nested RT-qPCR with detection limit of three viral RNA copies per million cells. **(C)** On day 28, cells were washed of any treatment and both ART or ART+GYY4137 treatment groups were stimulated with 1 μM prostratin for 24 h. HIV-1 RNA copies were assessed by RT-qPCR. Reduction in viral stimulation in GYY4137 treated samples are represented as percentage values. ND - non-determined. **(D)** Aggregate plot for 5 patients from data shown in C. **(E-F)** Primary CD4^+^ T cells expanded and cultured in presence of ART or ART+GYY4137 were analyzed overtime for the expression of activation (CD38) and quiescence markers (CD127) by flow cytometry. **(G)** Total HIV-1 DNA content was determined up to 28 days in ART or ART+GYY4137 treated groups. Results are expressed as mean ± standard deviation. **, P<0.01; ns, nonsignificant, by two-way ANOVA with Tukey’s multiple comparison test. Figure 6 includes the following figure supplement: **Figure supplement 1.** Phenotypic features of CD4^+^ T cells were preserved upon prolong treatment with GYY4137.

We next assessed the ability of GYY4137 to efficiently block viral reactivation. On day 28, we stimulated the CD4^+^ T cells with the protein kinase C (PKC) activator, prostratin, in the absence of any treatment. The activation of viral transcription was measured 24 h later by RT-qPCR. The removal of ART uniformly resulted in a viral reactivation by prostratin in CD4**^+^** T cells of HIV-patients (*Figure 6C*). In contrast, upon ART+GYY4137 removal followed by prostratin stimulation, viral reactivation was attenuated by 90 % (N = 5, mean) for all 5 patient samples, and individual inhibition ranged from 78.9 % to 100 % (*Figure 6C*). Overall, pretreatment with ART+GYY4137 significantly reduced prostratin-mediated HIV-reactivation when compared to ART alone (*p* < 0.01) (*Figure 6D*).

We also examined the immune-phenotype of CD4^+^ T cells treated with ART or ART+GYY4137 by monitoring the expression of activation (CD38) and quiescence (CD127) marker. As expected, CD38 expression increased, and CD127 expression decreased at day 7 and 14 during the activation phase, followed by a gradual reversal of the pattern during the resting phase at day 21 and 28 (*Figure 6E-F* and *Figure 6– figure supplement 1A*). Interestingly, GYY4137 treatment did not alter the temporal changes in the expression of activation and quiescence markers on CD4^+^ T cells (*Figure 6E-F*). Since memory CD4^+^ T cells are preferentially targeted by HIV (69, 70), we further analyzed if the status of naive (T_N_), central memory (T_CM_), transitional memory (T_TM_), and effector memory (T_EM_) is affected by the GYY4137 in our *ex vivo* expansion model. We observed that the CD4^+^ T cells that responded to *ex vivo* activation were mainly composed of T_TM_ and T_EM_ in our patient samples (*Figure 6–figure supplement 1B-E*). This is consistent with the study reporting the presence of translation-competent genomes mainly in the T_TM_ and T_EM_ of ART-suppressed individuals (71). Interestingly, the fraction of T_TM_ shows a progressive decline, whereas T_EM_ displays gradual increase during transition from activation (7-14 days) to quiescence phase (21-28 days) (*Figure 6–figure supplement 1D-E*). The addition of GYY4137 did not affect dynamic changes in the frequency of T_TM_ and T_EM_ subpopulations (*Figure 6–figure supplement 1D-E*).

Finally, we tested if the reduction in viral RNA upon GYY4137 treatment is due to the loss of cells with ability to reactivate virus or selection of cells subset that is non-responsive to reactivating stimuli. We did not find significant differences in total HIV DNA content between the CD4^+^ T cells immediately isolated from the patient’s PBMCs (*ex vivo*) and the expanded cells treated with ART+GYY4137 or ART alone for the entire duration of the experiment (*Figure 6G*). The viability of cells remained comparable between ART+GYY4137 and ART treatment groups overtime (*Figure 6– figure supplement 1F*). Altogether, using a range of cellular and immunological assays, we confirmed that the characteristics of an individual’s viral reservoir remain preserved, and the suppression of viral RNA upon GYY4137 treatment is the result of H_2_S-mediated inhibition of HIV-1 transcription rather than a reduction in proviral content or an altered CD4^+^ T cell subsets. In sum, prolonged exposure to GYY4137 results in potent inhibition of viral reactivation, suggesting a new H_2_S-based mechanism to neutralize bursts of virus reactivation under suppressive ART *in vivo*.

### GYY4137 prevents mitochondrial dysfunction in CD4^+^ T cells of HIV-patients during viral rebound

Because virus reactivation upon depletion of endogenous H_2_S resulted in mitochondrial dysfunction and redox imbalance in U1, we tested if the elevation of H_2_S levels by GYY4137 improves mitochondrial health of primary CD4**^+^** T cells during virus reactivation *ex vivo*. As described earlier, CD4^+^ T cells harboring latent virus upon prolonged (28 days) treatment of ART and ART+GYY4137 were stimulated by prostratin or left unstimulated and subjected to mitochondrial flux analysis. The unstimulated cells from both ART and ART+GYY4147 cultures did not show any difference in OCR (*Figure 7A*). In contrast, several features reflecting efficient mitochondrial activity such as basal respiration and ATP-coupled respiration were significantly higher in prostratin stimulated CD4^+^ T cells in case of ART+GYY4137 treatment than ART alone (*Figure 7B-C*). Consistent with this, mitoROS generation upon stimulation with prostratin or other LRAs such as PMA/ionomycin was reduced in ART+GYY4137 treated CD4^+^ T cells than ART alone (*p* = 0.005 and *p* = 0.014), and was nearly comparable to unstimulated cells (*Figure 7D-E*). The reduction in mitoROS could be a consequence of GYY4137-mediated increase in the expression of Nrf2-dependent antioxidant systems. We directly tested this by RT-qPCR analysis of a selected set of Nrf2-dependent genes on CD4^+^ T cells treated with GYY4137 for 28 days. Consistent with our findings in U1, treatment with ART+GYY4137 uniformly induced the expression of antioxidant genes in the latently infected CD4^+^ T cells compared to cell treated with ART alone (*Figure 7F*).

**Figure 7.**
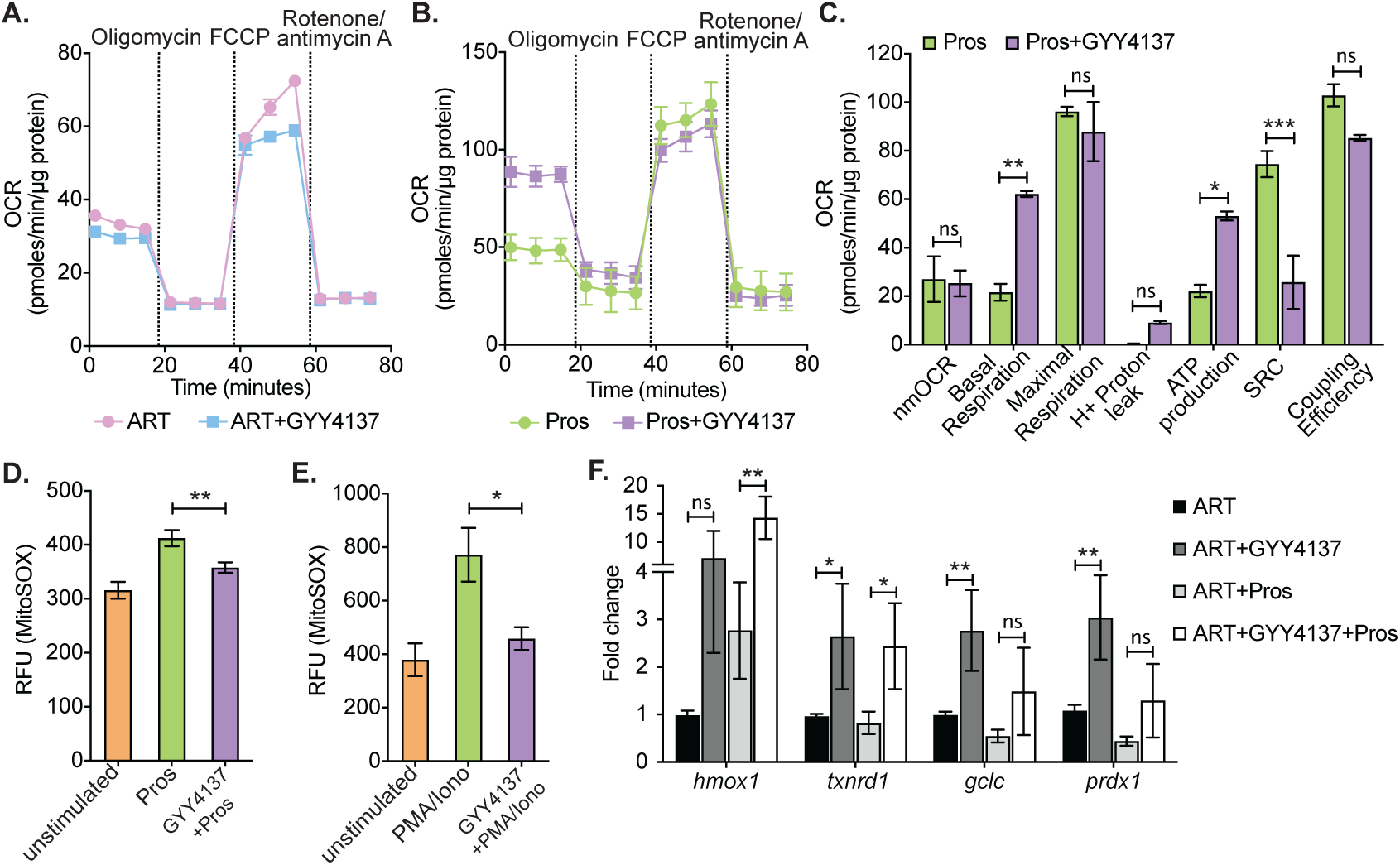
Effect of H_2_S on mitochondrial respiration and ROS generation in latent CD4^+^ T cells derived from HIV-1 patients. **(A)** Primary human CD4^+^ T cells from HIV infected subjects were activated and cultured *ex vivo* with ART or ART+GYY4137. On day 28, cells from ART or ART+GYY4137 treatment groups were harvested to assess mitochondrial respiration by using Seahorse XF mito-stress test as described in materials and methods. **(B)** Cells from ART and ART+GYY4137 treated groups were stimulated with 1 μM prostratin 6 h. Post-stimulation mitochondrial respiration profile was determined by Seahorse XF mito-stress test. **(C)** Various mitochondrial respiratory parameters derived from OCR measurement were determined by Wave desktop software. nmOCR; non-mitochondrial oxygen consumption rate and SRC; spare respiratory capacity. **(D)** On day 28, Cells from both ART and ART+GYY4137 treatment groups were stimulated with 1 μM Prostratin for 6 h. Cells were harvested and stained with 5 μM MitoSOX-Red dye for 30 min followed by washing. Samples were analyzed by flow cytometry. Unstimulated-cells cultured under ART alone. **(E)** Both ART and ART+GYY4137 treated cells at day 14 post-activation were stimulated with 1 μg/ml PMA and 100 μg/ml ionomycin (Iono) for 6 h. Cells were harvested post-stimulation and stained with 5 μM MitoSOX-Red dye. Samples were analyzed by flow cytometry to assess mitoROS generation. Unstimulated-cells cultured under ART alone. **(F)** CD4^+^ T cells from ART and ART+GYY4137 treated groups were stimulated with 1 μM prostratin on 28^th^ day for 24 h or left unstimulated. Cells were harvested to isolate total RNA and expression of *hmox1*, *txnrd1*, *gclc* and *prdx1* were determined by RT-qPCR. Data obtained were normalized to internal control β_2_ microglobulin (B2M). Error bar represent standard deviations from mean. Results are representative of data from three patients samples. *, P<0.05; **, P<0.01, by two-way ANOVA with Tukey’s multiple comparison test.

Altogether, these data suggest that H_2_S not only prevents virus reactivation but also improves mitochondrial bioenergetics and maintains redox homeostasis, which could be important for *in vivo* suppression of viral rebound and replenishment of the reservoir.

## Discussion

The major conclusion of our study is that HIV-1 reactivation is coupled to depletion of endogenous H_2_S, which is associated with dysfunctional mitochondrial bioenergetics, in particular suppressed OXPHOS, GSH/GSSG imbalance, and elevated mitoROS. Decreased H_2_S also impaired the expression of genes involved in maintenance of redox balance, mitochondrial function, inflammation, and HIV-1 restriction, which correlates with reactivation of HIV-1. This conclusion is supported in multiple cell line models of HIV-1 latency. Chemical complementation with GYY4137 identified H_2_S as an effector molecule. Finally, we confirmed clinical relevance of our finding in primary CD4^+^ T cells isolated from ART-suppressed patients to recapitulate features of HIV-1 latency and reactivation (60). Overall, our data show that while H_2_S deficiency reactivates HIV-1, the H_2_S donor GYY4137 can potently inhibit residual levels of HIV-1 transcription during suppressive ART and block virus reactivation upon stimulation. Hence, our findings highlight that H_2_S based therapies (72) can be exploited to lock HIV in a state of persistent latency by impairing the ability to reactivate.

How does H_2_S promote HIV-1 latency and suppress reactivation? Several studies revealed that HIV-1 reactivation is associated with loss of mitochondrial functions, oxidative stress, inflammation, and apoptosis (30, 31, 48, 57). Several of these biological dysfunctions are corrected by H_2_S. For example, localized delivery of H_2_S in physiological concentrations improves mitochondrial respiration, mitigates oxidative stress, and exerts anti-inflammatory functions (5, 16, 22, 44–47, 72). Our NanoString findings, XF flux analysis, and redox measurement supports a mechanism whereby H_2_S sufficiency promotes sustenance of mitochondrial health and redox homeostasis to control HIV-1 reactivation. Conversely, genetic depletion of endogenous H_2_S promotes HIV-1 reactivation by decelerating OXPHOS, GSH/GSSH imbalance, and elevated mitoROS. It is widely known that reduced OXPHOS leads to accumulation of NADH, which traps flavin mononucleotide (FMN) in the reduced state on respiratory complex I leading to mitoROS generation by incomplete reduction of O_2_ (73). Also, H_2_S-depletion increased nmOCR, which indicates elevated activities of other ROS producing enzymes such as NADPH oxidase and cyclooxygenases/lipoxygenase (74, 75). All of these metabolic changes are likely contributors of overwhelming ROS and disruption of GSH/GSSG poise in U1-shCTH, resulting in HIV-1 reactivation. In fact, consistent with our findings, oxidative shift in GSH/GSSG poise, deregulated mitochondrial bioenergetics, and ROI have been reported to promote HIV-1 reactivation (30, 31). The addition of GYY4137 stimulated OXPHOS and blocked HIV reactivation, which is in line with the known ability of H_2_S in sustaining respiration by acting as a substrate of cytochrome C oxidase (CcO) (47). Also, H_2_S can prevent mitochondria-mediated apoptosis (76), which is crucial for HIV-1 reactivation and infection (48). Finally, activation of Nrf2-specific antioxidant pathways upon exposure to GYY4137 agrees with the potential of H_2_S in augmenting the cellular antioxidative defense machinery (4, 16, 44). Importantly, cell lines (U1 and J1.1) modeling HIV-1 latency and PBMCs of individuals that naturally suppress HIV-1 reactivation (long-term non-progressors [LTNPs]) display robust capacities to resist oxidative stress and apoptosis (31, 77).

Our transcription profiling showed a positive correlation between GYY4137- induced H_2_S sufficiency and HIV-1 restriction factors (*e.g.,* YY1, APOBEC3, IRF1, Nrf2) and negative correlation with HIV-1 trans-activators (*e.g.,* NFKB, FOS), consistent with an interrelationship between cell metabolism and HIV latency (28, 78). Interestingly, a sustained induction of Nrf2-driven cellular antioxidant response is essential for the successful transition between productive and latent HIV-1 infection (31, 33). Moreover, inhibition of Nrf2 promoted viral transcription and increased ROS generation (33), whereas its activation reduced HIV-1 infection (65). On this basis, we think that H_2_S-mediated inhibition of HIV is partly related to activation of Nrf2- dependent antioxidant genes. In HIV infection, the suppression of viral reactivation and pro-inflammatory mediator expression was paralleled by the inhibition of NF-*κ*B. Interestingly, H_2_S inhibited activation of NF-*κ*B in several models of inflammatory diseases including hemorrhagic shock, lung injury, and paramyxovirus infections (79, 80). Similarly to these findings, we found that exposure of U1 with GYY4137 inhibited the NF-*κ*B p65 subunit and reduced its binding to HIV-1 LTR. Currently, no antiretroviral drugs inhibit basal transcription of provirus and blocks reactivation upon therapy (ART) interruption. It has been suggested that epigenetic silencing will be an important pre-requisite for persistent latency (81, 82). Our expression and ChIP-qPCR data suggest that H_2_S could exert its influence on HIV replication machinery via activating YY1- an epigenetic silencer of HIV-1 LTR (56, 68). Interestingly, another gasotransmitter NO modulates YY1 function via S-nitrosylation of its cysteine residues (83), indicating that YY1 is amenable to thiol-based-modifications and raises the possibility of YY1 S-persulfidation by H_2_S as a mechanism to epigenetically reconfigure HIV-1/promoter LTR.

The inefficiency of current ART in preventing virus rebound after therapy cessation poses a major hurdle to HIV cure. Clinical studies revealed that reservoir sizes have larger influence on viral rebound after ART cessation (84). For example, smaller reservoir size maintained due to sustained immunological control in case of a Mississippi baby (85), the VISCONTI cohort (86), and a French teenager (87) delayed viral rebound after treatment interruption. Using a primary cell system that has been successfully used to recapitulate latency and reactivation seen in patients (60), we confirmed that pretreatment with GYY4137 attenuated viral rebound in primary human CD4^+^ T cells upon stimulation with prostratin. Collectively, our results suggest that H_2_S reduces HIV-1 transcriptional activity, promotes silencing of its promoter, and reduces its potential for reactivation. The virus-suppressing effects of H_2_S were observed alongside its beneficial consequences on cellular physiology (e.g., mitochondrial function and redox balance). Both mitochondrial dysfunction and oxidative stress are major complications associated with long-term exposure to ART and contributes to inflammation and organ damage (88–90). Future studies will investigate if the prolonged treatment with H_2_S donors results in a better outcome due to recruitment of specific repressor and/or epigenetic silencer of HIV-1 LTR without detrimental consequences on cellular physiology. It’s tempting to speculate that over time (H_2_S in combination with ART) can push transcriptional repression beyond a certain threshold where it will be impossible to reactivate HIV, thereby blocking and locking virus in a state of persistent latency. Our results provide a proof-of-concept of a gasotransmitter H_2_S that can be explored for a functional cure of HIV. Blocking HIV rebound by inhibitors of viral factors (*e.g.,* Tat) invariably results in the emergence of resistant mutants (91). In this context, blocking HIV-1 rebound via H_2_S-directed modulation of host pathways could help in overcoming the problem of evolution of escape variants. Finally, therapy non-compliance and frequent blips contributes to continuous replenishment of latent reservoir *in vivo* (92). Combining H_2_S donors with ART regiments could potentially prevent reservoir replenishment during infection.

In conclusion, we identify H_2_S as a central factor in HIV-1 reactivation. Our systematic mechanistic dissection of the role of H_2_S in cellular bioenergetics, redox metabolism, and latency unifies many previous phenomena associated with viral persistence. Lastly, our results provide a rationale for including pharmacological donors of H_2_S in strategies to eradicate HIV *in vivo*.

## Material and methods

### Study design

The primary objective of this study was to understand the role H_2_S gas in modulating HIV latency and reactivation. First, we examined the differential expression of enzymes involved in H_2_S biogenesis and levels of H_2_S upon HIV-1 latency and reactivation. Next, we genetically silenced the expression of the main H_2_S producing gene CTH and showed its importance in maintaining HIV-1 latency. We performed detailed mechanistic studies on understanding the role of CTH in regulating cellular antioxidant response, mitochondrial respiration, ROS generation to maintain HIV-1 latency. Lastly, pharmacological donor of H_2_S (GYY4137) was used to reliably increase endogenous H_2_S levels and to study its consequence on HIV latency and rebound. Finally, primary CD4^+^ T cells derived from ART treated HIV infected patients were used to assess the effect of GYY4137 in modulating HIV-1 rebound by improving mitochondrial bioenergetics and mitigating redox stress.

### Subject samples

Peripheral blood mononuclear cells (PBMCs) were collected from five HIV-1 seropositive subjects on stable suppressive ART (*Table S5*). All subject signed informed consent forms approved by Indian Institute of Science, Bangalore and Bangalore Medical College and Research Institute (BMCRI) review boards (Institute human ethics committee [IHEC] No-3-14012020).

### Mammalian cell lines and culture conditions

The human pro-monocytic cell line U937, CD4^+^ T lymphocytic cell line Jurkat, HEK293T were procured from ATCC, Manassas, VA. The chronically infected ACH-2, J-Lat 6.3, U1, J1.1 and TZM-bl cell lines were obtained through AIDS Research and Reference Reagent Program, NIH, USA. The cell lines were cultured in RPMI1640 (Cell Clone) supplemented with L-glutamine (2 mM), 10% fetal bovine serum (MP biomedicals), penicillin (100 units/mL), streptomycin (100 μg/mL) at 37°C and 5% CO_2_. HEK293T and TZM-bl cells were cultured in DMEM (Cell Clone) supplemented with 10% FBS.

### Chemical reagents

Sodium hydrosulfide (NaHS), morpholin-4-ium 4-methoxphenyl(morpholino) phosphinodithioate dichloromethane complex (GYY4137), Phorbol-12-myristate-13-acetate (PMA), N-acetyl cysteine (NAC), prostratin and L-Cysteine were purchased from Sigma-Aldrich. Recombinant human TNF-*α* was purchased from InvivoGen. The antiretroviral drugs efavirenz, zidovudine, raltegravir, and lamivudine were obtained through the NIH AIDS Reagent Program.

### Latent viral reactivation

Latently infected U1 and J1.1 (2 *×* 10^5^ cells/ml) were stimulated with PMA (5 ng/ml) or TNF-*α* (100 ng/ml) for the time indicated in the figure legends. HIV-1 reactivation was determined by intracellular *gag* RT-qPCR or p24 estimation in supernatant by HIV-1 p24 ELISA (J. Mitra and Co. Pvt. Ltd., India). J-Lat 6.3 cells were stimulated with Prostratin (2.5 μM) for 24 h and HIV-1 reactivation was assessed by estimating GFP+ cells (excitation: 488 nm; emission: 510 nm) using BD FACSVerse flow cytometer (BD Biosciences). The data were analysed using FACSuite software (BD Biosciences).

### Reverse transcription quantitative PCR (RT-qPCR)

Total cellular RNA was isolated by RNeasy mini kit (Qiagen), according to the manufacturer’s protocol. RNA (500 ng) was reverse transcribed to cDNA (iScript*™* cDNA synthesis kit, Bio-Rad), subjected to quantitative real-time PCR (iQ*™* SYBR^®^ Green Supermix, Bio-Rad), and performed using the Bio-Rad C1000*™* real time PCR system. HIV-1 reactivation was assessed using gene-specific primers (*Table S6*). The expression level of each gene is normalized to human β-actin as an internal reference gene.

### Western blot analysis

Total cell lysates of PMA-treated and untreated U1, J1.1, U937, and Jurkat cell lines were prepared using radioimmunoprecipitation (RIPA) lysis buffer (50 mM Tris [pH 8.0], 150 mM NaCl, 1% Triton X-100, 1 % sodium deoxycholate, 0.1% SDS (sodium dodecyl sulfate), 1*×* protease inhibitor cocktail (Sigma-Aldrich), 1*×* phosphatase inhibitor cocktail (Sigma-Aldrich). After incubation on ice for 20 min, the lysates were centrifuged at 12,000 rpm, 4°C for 15 mins. Clarified supernatant was taken and total protein concentration was determined by Bicinchoninic acid assay (Pierce*™*, Thermo Fisher Scientific). Total protein extracts were separated by SDS-PAGE and transferred onto polyvinylidene difluoride membranes. Membranes were probed with an anti-CBS (EPR8579), CTH (ab151769), MPST (ab154514), and anti-HIV-1 p24 (ab9071) from abcam; Nrf2 (CST-12721), Keap1 (CST-4678), NF-*κ*B p65 (CST-6956), phospho-NF-*κ*B p65 (Ser536) (CST-3033), YY1 (CST-63227) and GAPDH (CST-97166) from Cell Signaling Technologies, Inc. Anti-rabbit IgG (CST-7074) and anti-mouse IgG (CST-7076) were used a secondary antibodies. Proteins were detected by ECL and visualized by chemiluminescence (Perkin Elmer, Waltham, MA) using the Bio-Rad Chemidoc Imaging system (Hercules, CA). For membrane reprobing, stripping buffer was used (2% SDS [w/v], 62 mM Tris-Cl buffer (0.5 M, pH 6.7) and 100 mM *β*-mercaptoethanol) for 20 min at 55°C. After extensive washing with PBS containing 0.1 % Tween 20 (Sigma-Aldrich), membrane was blocked and reincubated with desired antibodies.

### H_2_S detection assays

Endogenous H_2_S levels of U1 and U937 cells were detected using H_2_S specific fluorescent probe as described (43). Briefly, cells were treated with PMA (5 ng/ml) for 24 h, washed with 1*×* PBS, and stained with 200 μM 7-Azido-4-Methylcoumarin (AzMC) (Sigma). The stained cells were mounted on a glass slide and visualized using Leica TCS SP5 confocal microscope (excitation: 405 nm; emission: 450 nm). Images obtained were analyzed using LAS AF Lite software (Leica Microsystems) and semi-quantification of 50 cells was performed using ImageJ software.

H_2_S generation was also measured using methylene blue assay. The supernatant of U1 cells treated with NaHS or GYY4137 was incubated with Zinc acetate (1%) and NaOH (3%) (1:1 ratio) to trap H_2_S for 30 min. The reaction was terminated using 10% trichloroacetic acid solution. Following this, reactants were incubated with 20 mM *N,N*-dimethylphenylendiamine (NNDPD) in 7.2 M HCl and 30 mM FeCl_3_ in 1.2 M HCl for 30 mins and absorbance was measured at 670 nm. The concentration of H_2_S was determined by plotting absorbance on a standard curve generated using NaHS (0-400 μM; R^2^=0.9982).

### Stable cell line generation

For generating CBS and CTH knockdown in U1 and J1.1 cells, we used validated pooled gene specific shRNAs from the RNAi Consortium (TRC) library (Sigma Aldrich, USA; shRNA sequences given in *Table S1*). The lentiviral particles were generated in HEK293T cells using the packaging vectors, psPAX2 and pMD2.G. The pLKO.1-puro vector encoding a non-mammalian targeting shRNA (shNT) was used as a control. The U1 cells were transduced with lentiviral particles in opti-MEM containing polybrene (10 μg/ml) for 6 hours. Cells were washed and stable clones were selected in culture medium containing 250 ng/ml of puromycin. Total RNA or cell lysates were prepared to validate knockdown of CBS and CTH.

### Intracellular HIV-1 p24 staining

For intracellular p24 staining, U1-shCTH and U1-shNT cells were stimulated with PMA (5 ng/ml), washed with PBS followed by fixation and permeabilization using a fixation and permeabilization kit (eBiosciences). Permeabilized cells were then incubated with 50 μl of 1:100 dilution of phycoerythrin (PE)-conjugated mouse anti-p24 monoclonal antibody (KC57-RD1; Beckman Coulter, Inc.) for 30 min at room temperature. After incubation the cells were washed twice and the fluorescence of stained samples were acquired using BD FACSVerse flow cytometer (BD Biosciences). The data were analysed using FACSuite software (BD Biosciences).

### NanoString nCounter assay

Total RNA was isolated using an RNeasy mini kit (Qiagen) according to manufacturer’s instructions. RNA concentration and purity were measured using a Nanodrop spectrophotometer (Thermo Fisher Scientific, Waltham, MA), bioanalyzers systems (Agilent Technologies, Inc.), and Qubit Assays (Thermo Fisher Scientific). An nCounter gene expression assay was performed according to the manufacturer’s protocol. The assay utilized a custom-made NanoString codeset designed to measure 185 genes, including 6 house-keeping genes (*Table S2*). This custom-made panel included genes associated with oxidative stress and HIV-1 infection (*Table S2*) All genes were assayed simultaneously in multiplexed reactions and analyzed by fully automated nCounter Prep Station and digital analyzer (NanoString Technologies). The data were normalized to B2M, used as a housekeeping gene due to its minimum % CV across the samples, and analysis was done using nSolver 4.0 software.

### Measurement of Oxygen Consumption Rates

Oxygen consumption rates (OCR) were measured using a Seahorse XFp extracellular flux analyzer (Agilent Technologies) as per manufacturer’s instructions. Briefly, cells (U1 or Primary CD4^+^ T cells) were seeded at a density of 10^4^-10^5^ per well in a Seahorse flux analyzer plate precoated with Cell-Tak (Corning). Cells were incubated for 1 h in a non-CO_2_ incubator at 37°C before loading the plate in the seahorse analyzer. To assess mitochondrial respiration, three OCR measurements were performed without any inhibitor in XF assay media to measure basal respiration, followed by sequential addition of oligomycin (1 μM), an ATP synthase inhibitor (complex V) and three OCR measurements to determine ATP-linked OCR and proton leakage. Next, cyanide-4-(trifluoromethoxy)phenylhydrazone (FCCP; 0.25 μM), was injected to determine the maximal respiration rate and the spare respiratory capacity (SRC). Finally, rotenone (0.5 μM) and antimycin A (0.5 μM), inhibitors of NADH dehydrogenase (complex I) and cytochrome *c* - oxidoreductase (complex III), respectively, were injected to completely shut down the electron transport chain (ETC) to analyze non-mitochondrial oxygen consumption rate (nmOCR). Seahorse data were normalized to total amount of protein (μg) and mitochondrial respiration parameters were analyzed using Wave Desktop 2.6 software (Agilent Technologies).

### Estimation of intracellular glutathione content and ROS

Total cell lysate was prepared from 10^7^ cells using sonication in MES (2-(N-morpholino)ethanesulfonic acid) buffer. Lysates were clarified by centrifugation and total protein concentration was estimated using BCA assay. Total glutathione, reduced glutathione (GSH), and oxidized glutathione (GSSG) were measured using the glutathione assay kit (Cayman Chemical, Ann Arbor, MI, USA) according to the manufacturer’s instructions. To measure ROS, cells were loaded with 10 μM of CM-H_2_DCFDA (Excitation: 492 nm; Emission; 517 nm) or 5 μM of MitoSOX*™*Red (Excitation: 510 nm; Emission; 580 nm) for 30 min at 37°C and exposed to H_2_O_2_ (100 μM) or antimycin A (2 μM) or PMA (5 ng/ml) or left untreated at 37°C and 5% CO_2_. The fluorescence of stained samples was acquired using BD FACSVerse flow cytometer (BD Biosciences). Data were analyzed using FlowJo software (BD Biosciences).

### Virus Production

HIV-1 particle production was carried out using Lipofectamine 2000 transfection reagent (Invitrogen, Life Technologies), according to the manufacturer’s protocol, in HEK293T cells using HIV-1 NL4-3 DNA (NIH AIDS Reagent Program, Division of AIDS, NIAID, NIH). The medium was replaced with fresh medium at 6 h post-transfection, and supernatants were collected after 60 h, centrifuged (10 min, 200 *×*g, room temperature), and filtered through a 0.45 μm-pore-size membrane filter (MDI; Membrane Technologies) to clear cell debris. Virus was concentrated using 5*×* PEG- *it™* (System Biosciences) as per manufacturer’s protocol and virus pellet obtained was aliquoted in opti-MEM and stored at -80°C. Viral titration was done using HIV-1 reporter cell line, TZM-bl (NIH AIDS reagent program) as described earlier (93).

### Assessing the effect of GYY4137 on HIV-1 replication

Briefly, 0.5*×*10^6^ Jurkat cells were infected with HIV-1 NL4.3 at 0.2 M.O.I for 4 h at 37°C. Post-infection cells were washed with 1*×* PBS and cultured in presence or absence of 300 μM GYY4137. Cells and culture supernatants were harvested to isolate total RNA for *gag* RT-qPCR and p24 estimation, respectively. The p24 concentration was determined in the culture supernatant by sandwich HIV-1 p24 ELISA (J. Mitra and Co. Pvt. Ltd., India) according to the manufacturer’s instruction.

### Chromatin immunoprecipitation and quantitative genomic PCR

The chromatin immunoprecipitation (ChIP) assays was performed using SimpleChIP*®* Enzymatic Chromatin IP Kit (Magnetic Beads) (Cell Signaling Technologies, Inc., MA, US) according to the manufacturer’s instructions. Briefly, 10^7^ U1 cells were pre-treated with GYY4137 (5 mM) for 6 h or left untreated and then stimulated with PMA (30 ng/ml) for 4 h. Cells were fixed using 1% formaldehyde, neutralized with glycine, and harvested in ice-cold PBS. Cells were lysed and nuclei were sheared by sonication using Bioruptor*®* Pico (Diagenode Inc.) to obtain DNA fragments of 200 to 500 nucleotides. The clarified lysates were immunoprecipitated using ChIP-grade anti-NF-*κ*B p65 (CST-6956) or anti-YY1 (CST-63227) or normal rabbit IgG (as a negative control) for overnight at 4°C followed by incubation with ChIP-grade protein G magnetic beads at 4°C for 4 h. Chromatin was eluted from protein G beads, reverse-cross linked, and DNA was column purified. Quantitative genomic PCR was done by SYBR green-based real-time PCR using primers spanning the ER, RBE I, and RBE III regions on HIV-1 LTR. The GAPDH gene was used as the reference gene to see non-specific binding. Total input (10 %) was used to normalize equal amount of chromatin taken across the samples. The relative proportion of co-immunoprecipitated DNA fragments were determined with the help of threshold cycle (*C_T_*) values for each qPCR product using the equation (100 *×*2^[CT(input-3.32)- CT(IP)]^). The data obtained were represented as fold enrichment normalized to IgG background for each IP reaction.

### Generation of expanded primary CD4^+^ T cells from aviremic subjects

The PBMCs isolated from blood samples using Histopaque-1077 (Sigma-Aldrich) density gradient centrifugation were used for CD4^+^ T cells purification. The CD4^+^ T cells were purified from 50 × 10^6^ PBMCs using EasySep^™^Human CD4^+^ T cell isolation kit (STEMCELL Technologies). Primary CD4^+^ T cells were cultured at 37°C in a 5 % (v/v) CO_2_ humidified atmosphere in Gibco*®*RPMI 1640 medium (Life Technologies) with GlutaMAX, HEPES, 100 U/ml interleukin-2 (IL-2; Peprotech, London, United Kingdom), 1 μg/mL phytohemagglutinin (PHA) (Thermo Fisher Scientific), gamma-irradiated feeder PBMCs (healthy control) and either ART alone (100 nM efavirenz, 180 nM zidovudine, and 200 nM raltegravir) or ART + 100 μM GYY4137. After 7 days, CD4^+^ T cells were cultured only with ART alone or ART + GYY4137 and 100 U/ml IL-2. For stimulation experiments on day 28, ART and ART + GYY4137 were washed off and 1 × 10^6^ cells were treated with 1 µM prostratin for 24 h.

### HIV-1 RNA and DNA isolation from primary CD4^+^ T cells

CD4^+^ T cells (1 × 10^6^) from ART or ART + GYY4137 treated groups were harvested to isolate total RNA using Qiagen RNAeasy isolation kit and 200 ng of total RNA was reverse transcribed (iScript*^™^*cDNA synthesis kit, Bio-Rad). Reverse transcribed cDNA was diluted 10-fold and amplified using primers against HIV LTRs and seminested – PCR was performed using primers and probe listed in *Table S6*. Serially diluted pNL4.3 plasmid was used to obtain the standard curve. Isolation and RT-qPCR of total HIV DNA was performed as described earlier (60). Briefly, 1 × 10^6^ cells were lysed (10 mM Tris-HCl, 50 nM KCl, 400 mg/ml proteinase K) at 55°C for 16 h followed by inactivation at 95°C for 5 min. Digested product was used as a template to set up first PCR with Taq polymerase (NEB), 1X Taq buffer, dNTPs, HIV and CD3 primers for 12 cycles. The second round amplification was done using seminested PCR strategy wherein 10 fold dilution of first round PCR product was used as a template, HIV and CD3 primers/probes, Taqman™ Fast Advance master mix (Applied Biosystems*™*) using SetupOnePlus*™* Real-time PCR system (Applied Biosystems*™*). DNA isolated from ACH2 cells that contain single copy of HIV per cell was used to obtain standard curve.

### Surface marker analysis of primary CD4^+^ T cells derived from HIV patients

CD4^+^ T cells derived from HIV-1 infected patients were stained for surface markers using monoclonal antibodies: CD4-BUV395 (SK3), CD45RA-APC-H7 (HI100), CD27-BV785, CCR7-Alexa 647 (G043H7), CD38-PE-Cy5 and CD127-PerCP-Cy5.5. Additionally, cells were stained with Live/Dead fixable Aqua dead cell stain or AviD (Invitrogen) to exclude dead cells from the analysis as per manufacturer’s instructions. Stained samples were run on BD FACSAria*™* Fusion flow cytometer (BD Biosciences, San Jose, CA) and data were analyzed with FlowJo version 9.9.6 software (Treestar, Ashland, OR).

### Statistical analysis

All statistical analyses were performed using GraphPad Prism software for Macintosh (version 9.0.0). The data values are indicated as mean ± S.D. For statistical analysis Student *t*-test (in which two groups are compared) and one-way or two-way ANOVA (for analysis involving multiple groups) were used. Analysis of NanoString data was performed using the nSolver platform. Differences in *P* values <0.05 and fold change >1.5 were considered significant.

## Acknowledgements

We are grateful to Prof. C. Grundner at the Seattle Children’s Research Institute for critical reading of the manuscript and valuable input. We acknowledge Dr. D. K. Saini at Department of Molecular Reproduction, Development and Genetics, IISc for providing shRNA constructs used in this study. We are grateful to Ms. S. Kumar and Dr. N. R. Sundaresan at Department of Microbiology and Cell Biology, IISc for helping with ChIP experiments carried out in this study. We gratefully acknowledge the NanoString services provided by TheraCUES Innovations Pvt Ltd, Bangalore

## Authors contributions

VKP and AS participated in the design of the study. VKP, RA, and SR carried out the experiment. VKP, PS, SR, and DTNM contributed in recruiting and isolating PBMCs from ART treated HIV-1 infected subjects. VKP, SR, AV, and AS contributed to reagents and analyzed the data. VKP and AS conceived the study, supervised the project, and drafted the manuscript. All authors read and approved the final manuscript.

## Data Availability

This study includes no data deposited in external repositories.

## Additional data files

**Figure 2-source data file 1.** This file contains RAW values associated with the NanoString analysis of host genes affected upon depletion of CTH in U1.

**Figure 5-source data file 1.** This file contains RAW values associated with the NanoString analysis of host genes affected upon PMA induced HIV reactivation in presence or absence of GYY4137 treatment.

## Supplementary Materials

**Figure 1-figure supplement 1.**
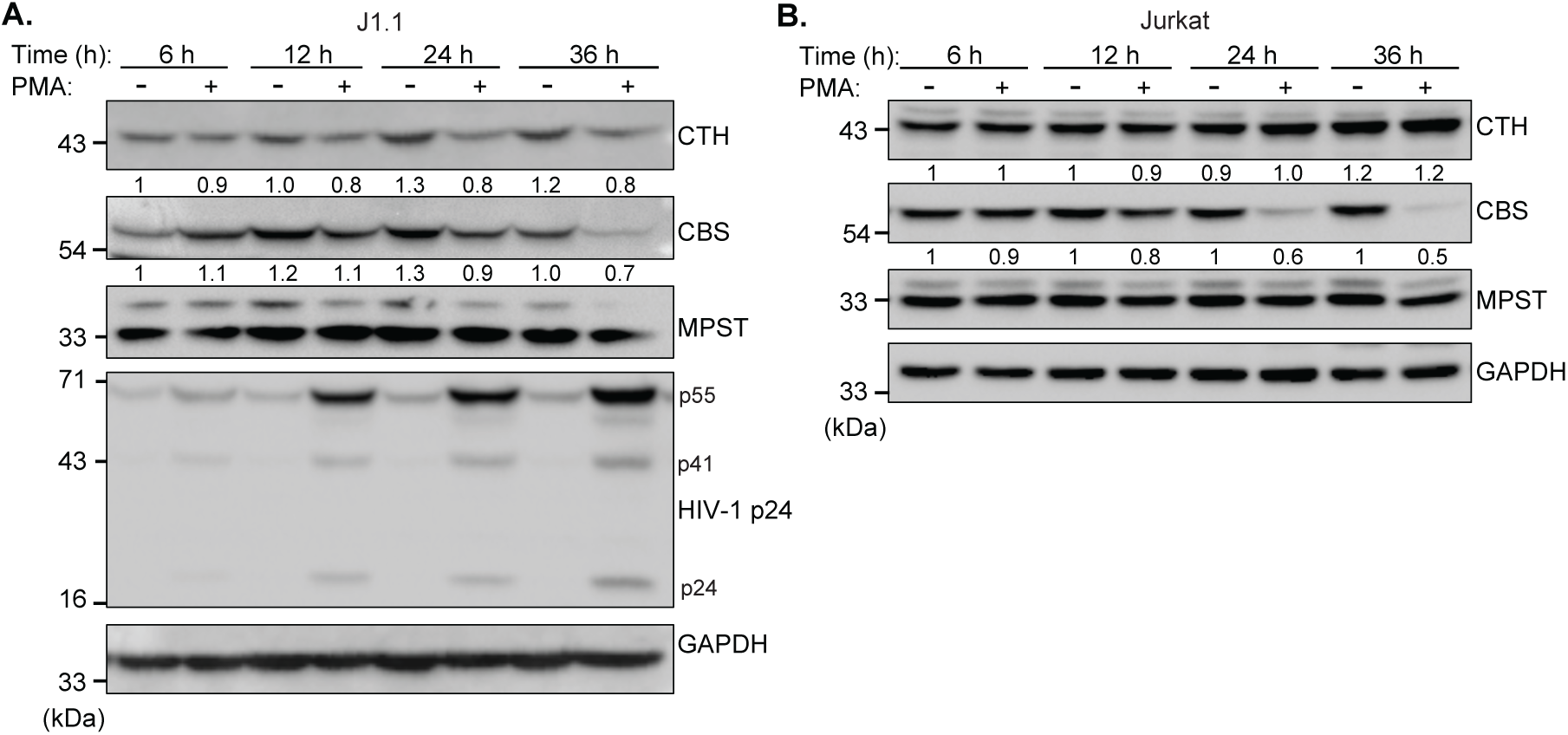
HIV-1 reactivation decreases levels of H_2_S metabolizing enzymes in a T cell line model of latency. **(A-B)** J1.1 and Jurkat cells were stimulated with 5 ng/ml PMA for 6 h, 12 h, 24 h, and 36 h and total cell lysates were prepared to analyze CTH, CBS, and MPST protein levels. HIV-1 reactivation was assessed by immunoblotting for intracellular viral protein, p24. The results are representative of data from three independent experiments. Results were quantified by densitometric analysis for CTH, CBS, and MPST band intensities and normalized to GAPDH.

**Figure 2-figure supplement 1.**
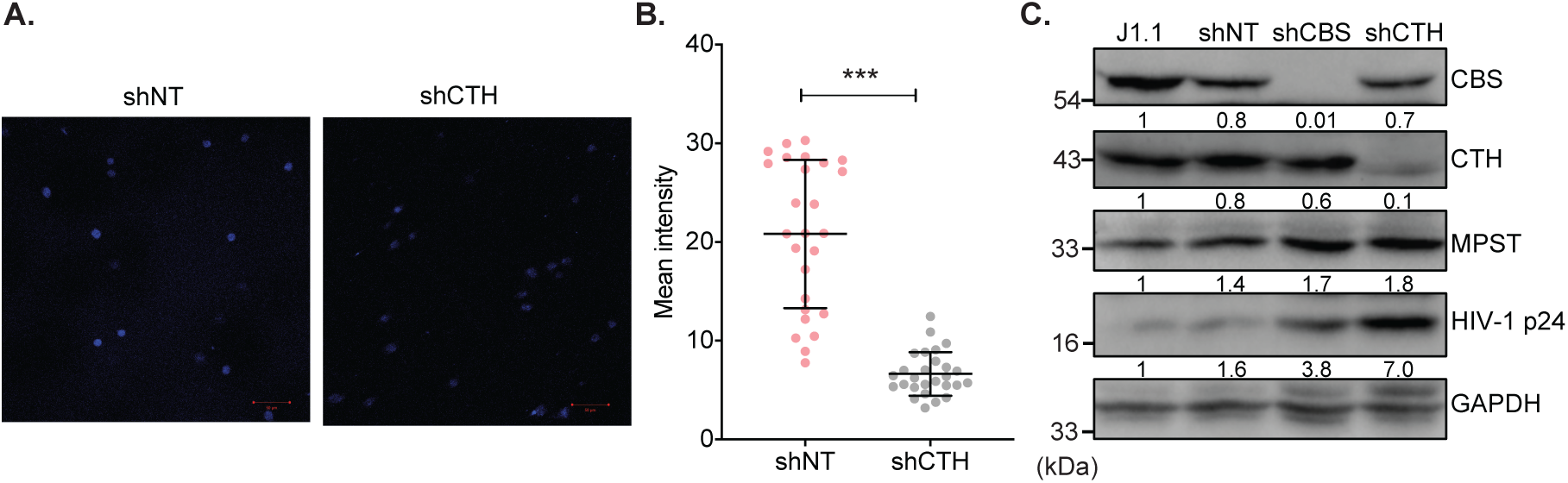
Genetic silencing of CTH reduces endogenous H_2_S levels and reactivates HIV-1 from latency. **(A and B)** U1 cells stably transduced with non-targeting (shNT) and shCTH lentiviral vectors were stained with AzMC for 30 min at 37°C, and images were acquired using ZEISS LSM 880 confocal microscope (A). Scale bar represents 20 μm. Average fluorescence intensity was quantified by ImageJ software (B). Results are expressed as mean ± standard deviation and are representative of data from two independent experiments. ***, P<0.001, by two-tailed student’s t-test. **(C)** Total cell lysates were prepared from un-transduced (J1.1), non-targeting (shNT), shCBS, and shCTH knockdown J1.1 cells. Expression of CBS, CTH, MPST and HIV-1 p24 levels was assessed by immuno-blotting. Results are expressed as mean ± standard deviation and are representative of data from two independent experiments.

**Figure 3-figure supplement 1.**
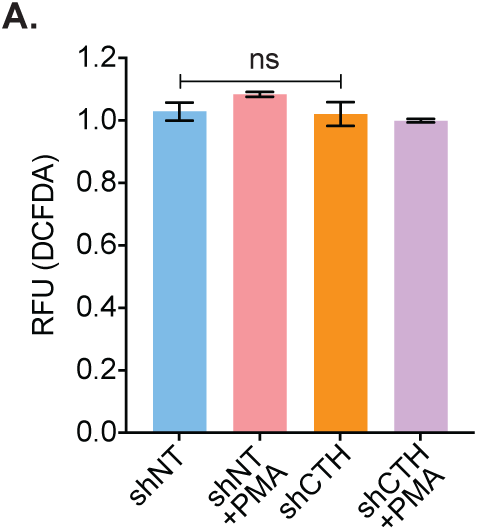
Effect of CTH knockdown on cytosolic ROS generation. **(A)** U1-shNT and U1- shCTH cells were left untreated or treated with 5 ng/ml PMA for 6 h, stained with CM-H_2_DCFDA dye (5 μM) analyzed using flow cytometry. Results are expressed as mean ± standard deviation and data are representative of two independent experiment. ns, nonsignificant, by two-way ANOVA with Tukey’s multiple comparison test.

**Figure 4-figure supplement 1.**
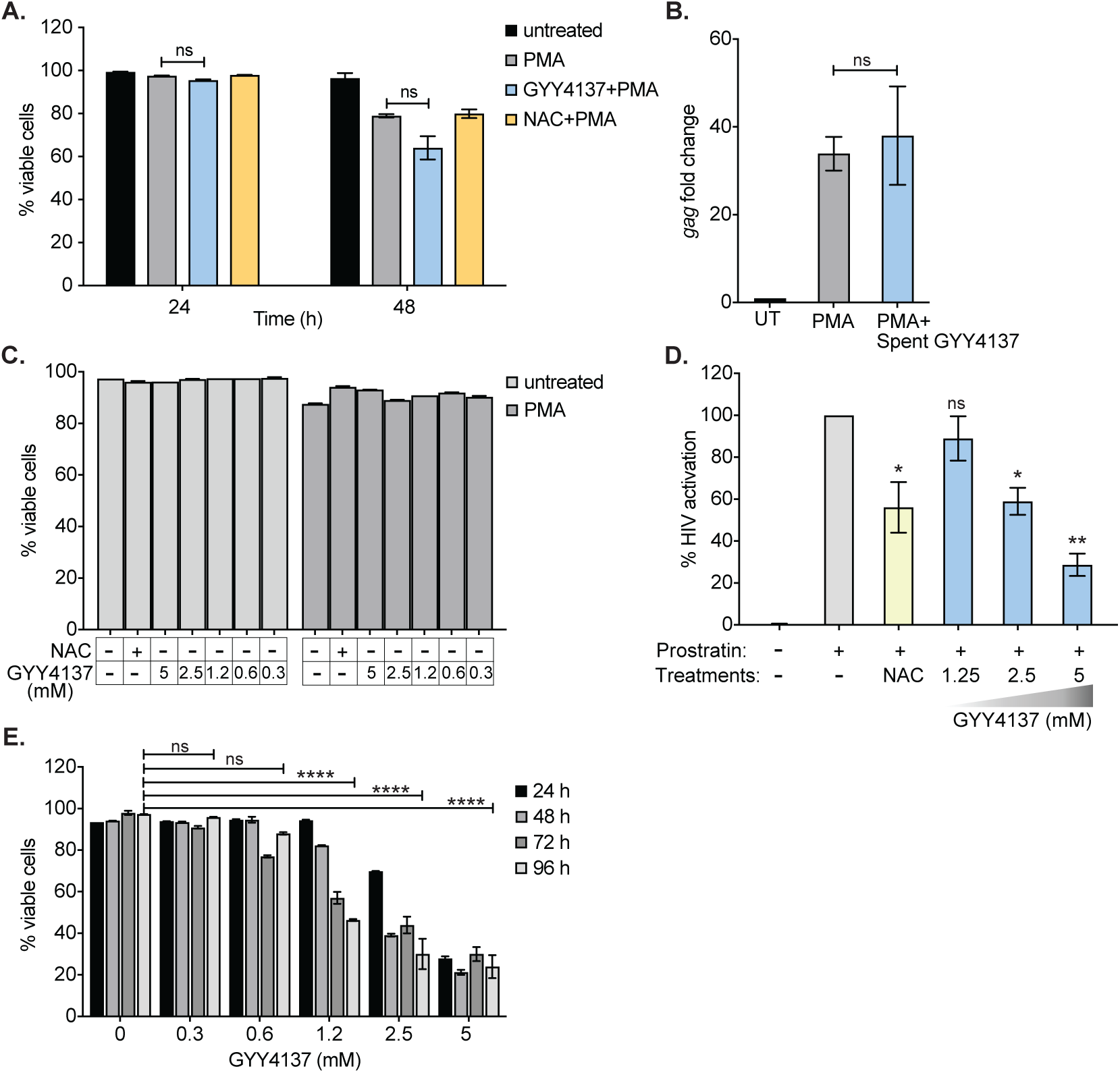
Effect of GYY4137 on HIV-1 reactivation and cellular viability. **(A)** U1 cells were pretreated with 5 mM GYY4137 or 5 mM NAC for 24 and then stimulated with 5 ng/ml PMA for 24 h or 48 h. Cells were stained with 3 μM propidium iodide (PI) for 15 min in dark, washed, and analyzed using flow cytometry. **(B)** Decomposed GYY4137 (spent GYY4137) which was aerated for at least 180 days at left at room temperature was used to pretreat U1 cells. Cells pretreated with 5 mM spent GYY4137 for 24 h or left untreated were stimulated with 5 ng/ml PMA for 24 h and HIV-1 reactivation was assessed by *gag* RT-qPCR. **(C)** J1.1 cells were pretreated with GYY4137 for 24 h and then stimulated with PMA for 12 h. Cells were then harvested, stained with PI, and subjected to flow cytometry to assess viability. **(D)** J-Lat cells were pretreated with 5 mM NAC or indicated concentrations of GYY4137 for 24 h and then stimulated with 2.5 μM prostratin for 24 h. HIV-1 reactivation was determined by estimating GFP expressing cells using flow cytometry. **(E)** Jurkat cells were treated with 5 mM, 2.5 mM, 1.2 mM, 0.6 mM and 0.3 mM of GYY4137 for 24 h, 48 h, 72 h, and 96 h. Samples were harvested and stained with PI to determine cells viability by flow cytometry. Error bar represent standard deviations from mean. Results are representative of data from two independent experiments. *, P<0.05; **,P<0.01; ***,P<0.001; ****, P<0.0001; ns, nonsignificant, by two-way ANOVA with Tukey’s multiple comparison test.

**Figure 6-figure Supplement 1.**
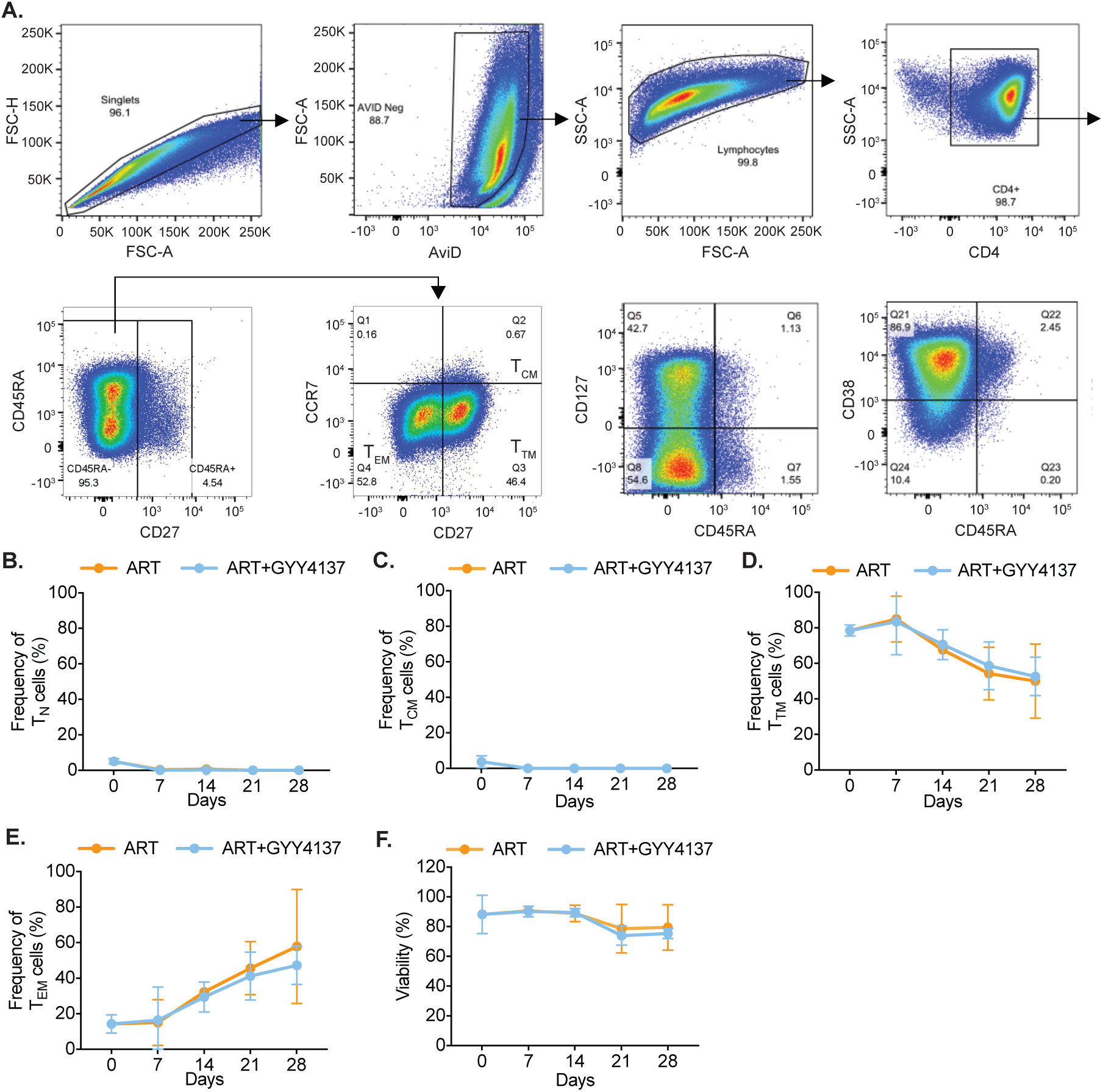
Phenotypic features of CD4^+^ T cells were preserved upon prolong treatment with GYY4137. **(A)** Flow cytometry-gating strategy used to analyze the expression of activation (CD38), quiescence (CD127), and frequency of different memory subsets of CD4^+^ T cells. The figure represents expression of different markers at Day 7 post activation of CD4^+^ T cells from a single patient sample. **(B-E)** Primary human CD4^+^ T cells from HIV infected patients cultured with ART or ART with GYY4137 were analyzed by flow cytometry overtime to determine frequency of different subsets of CD4^+^ T cells- T_N_ (naive [B]) and T_CM_ (central memory [C]), T_TM_ (transition memory [D]), T_EM_ (effector memory [E]). **(F)** Cell viability was assessed by Live-Dead staining overtime. Results obtained suggest no difference in viability between ART alone and ART with GYY4137 treated group overtime. Error bar represent standard deviations from mean. Results are representative of data from four patients samples.

**Table S1.**
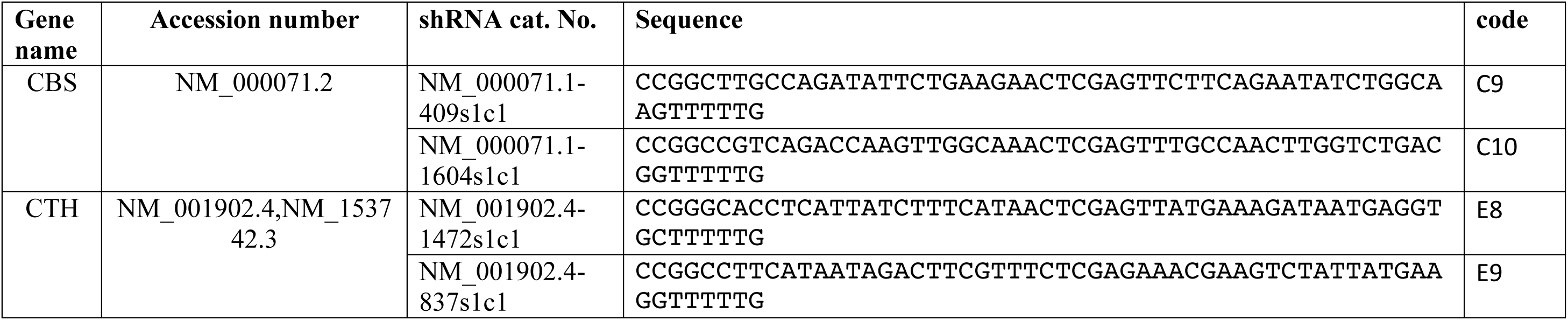
Sequences of shRNA clones.

**Table S2.**
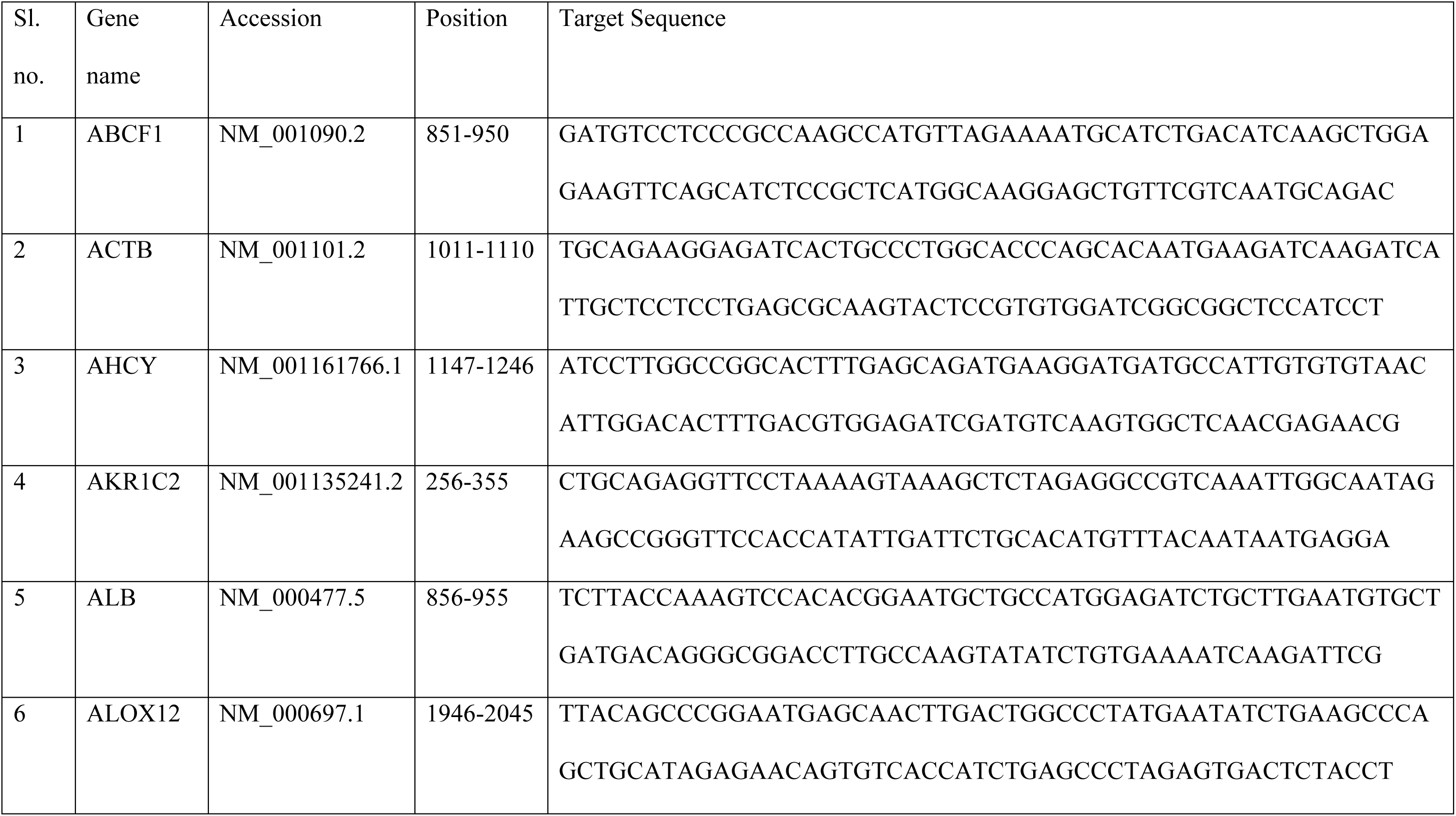
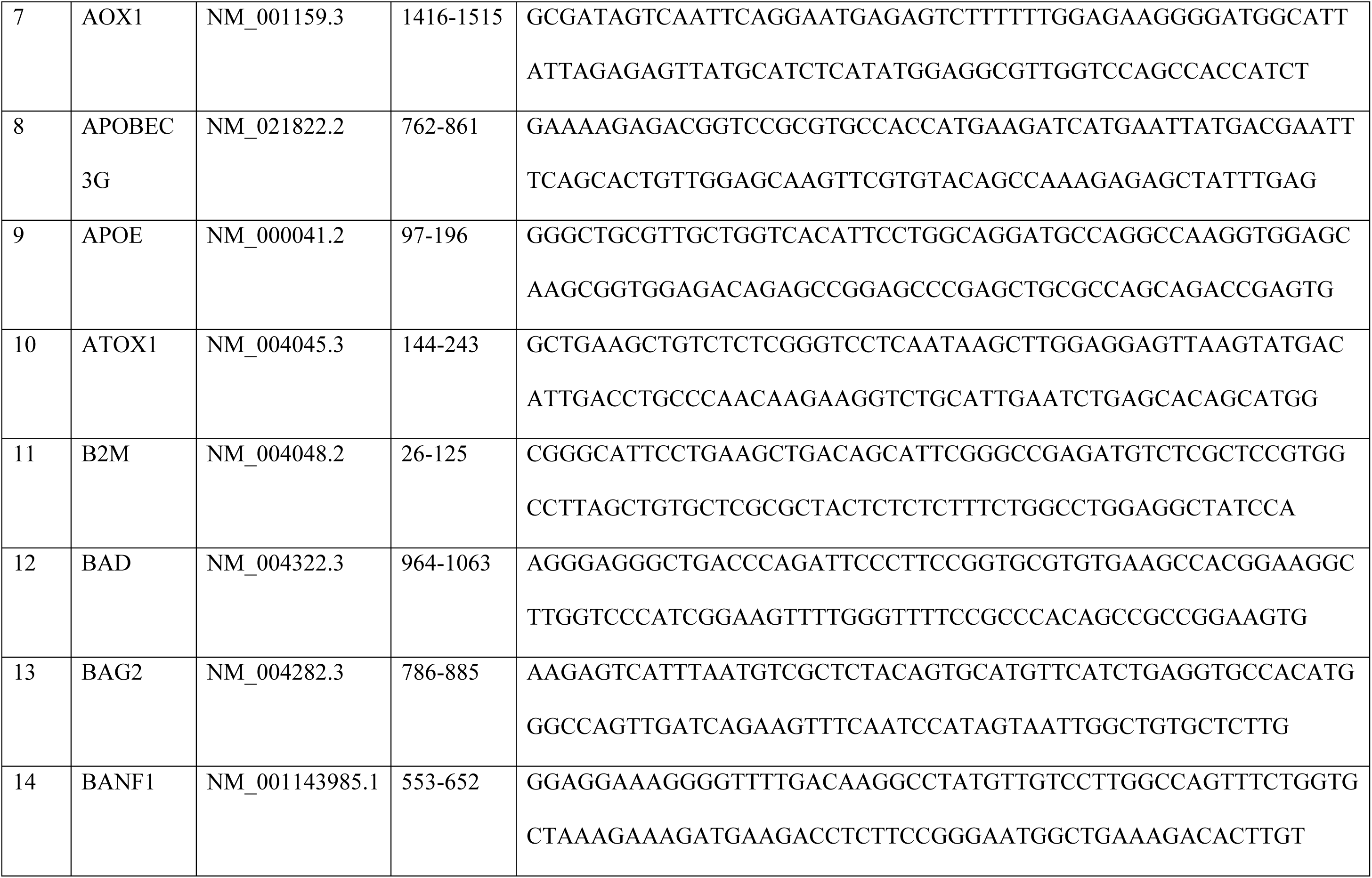
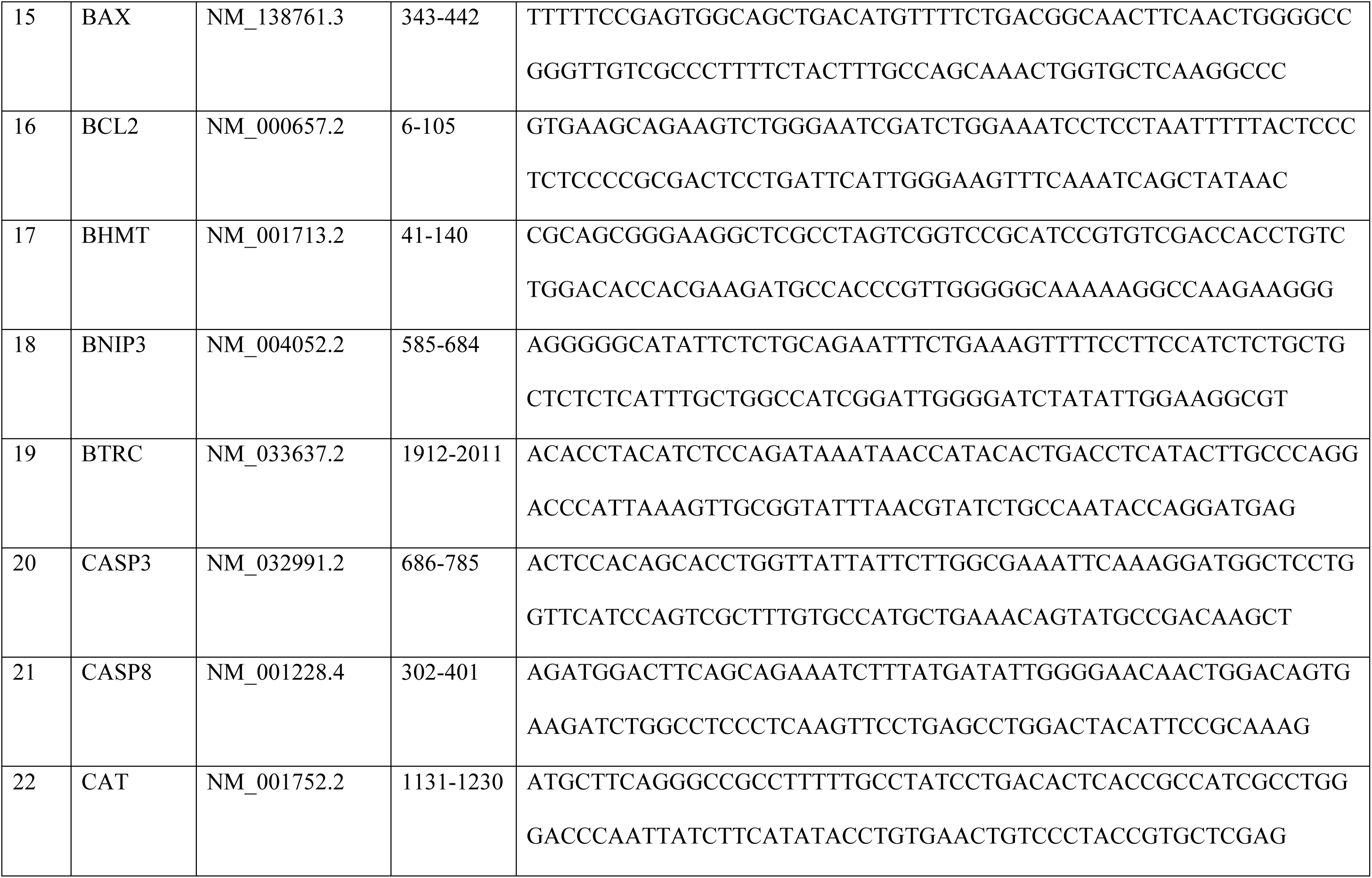
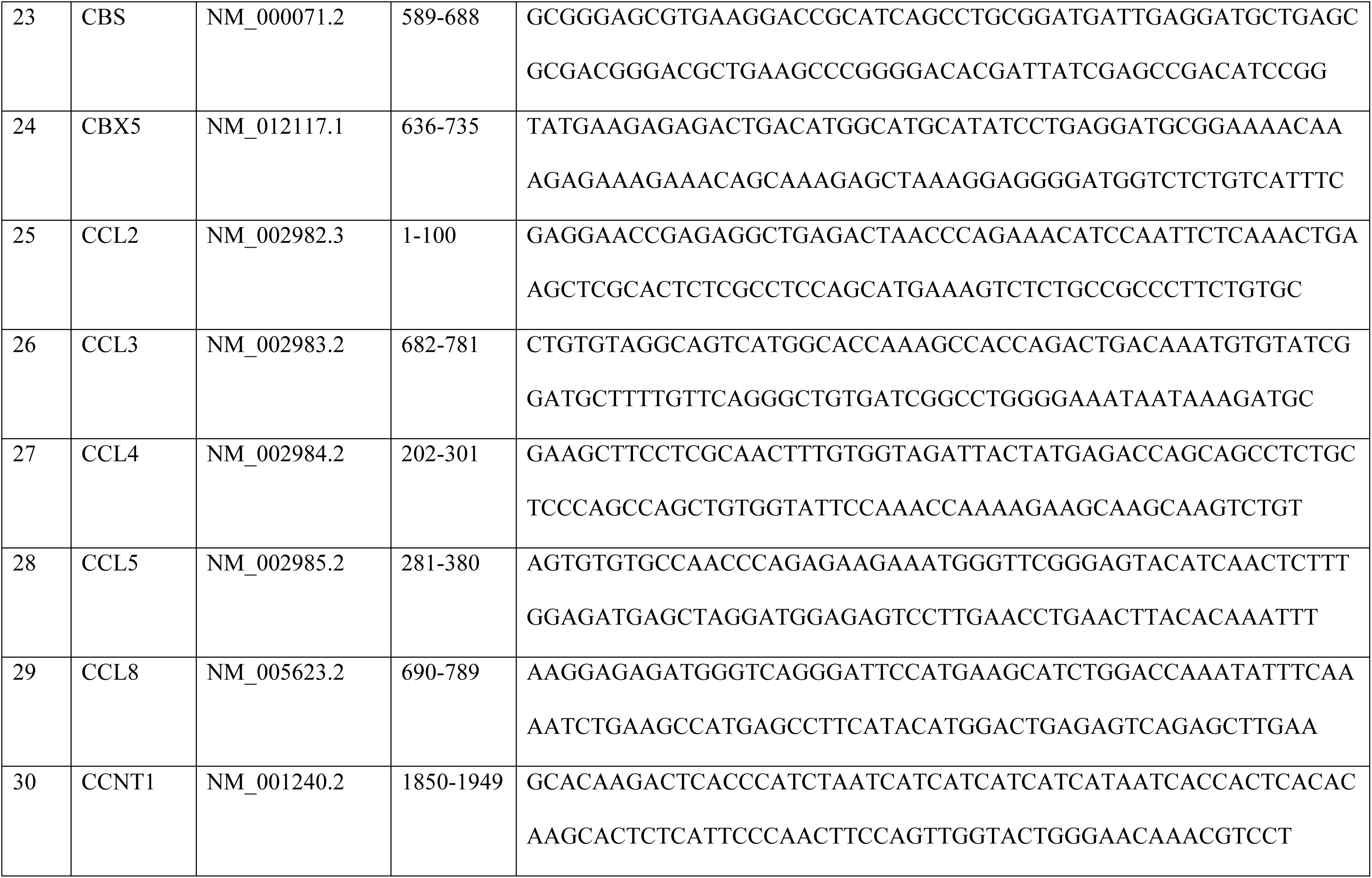
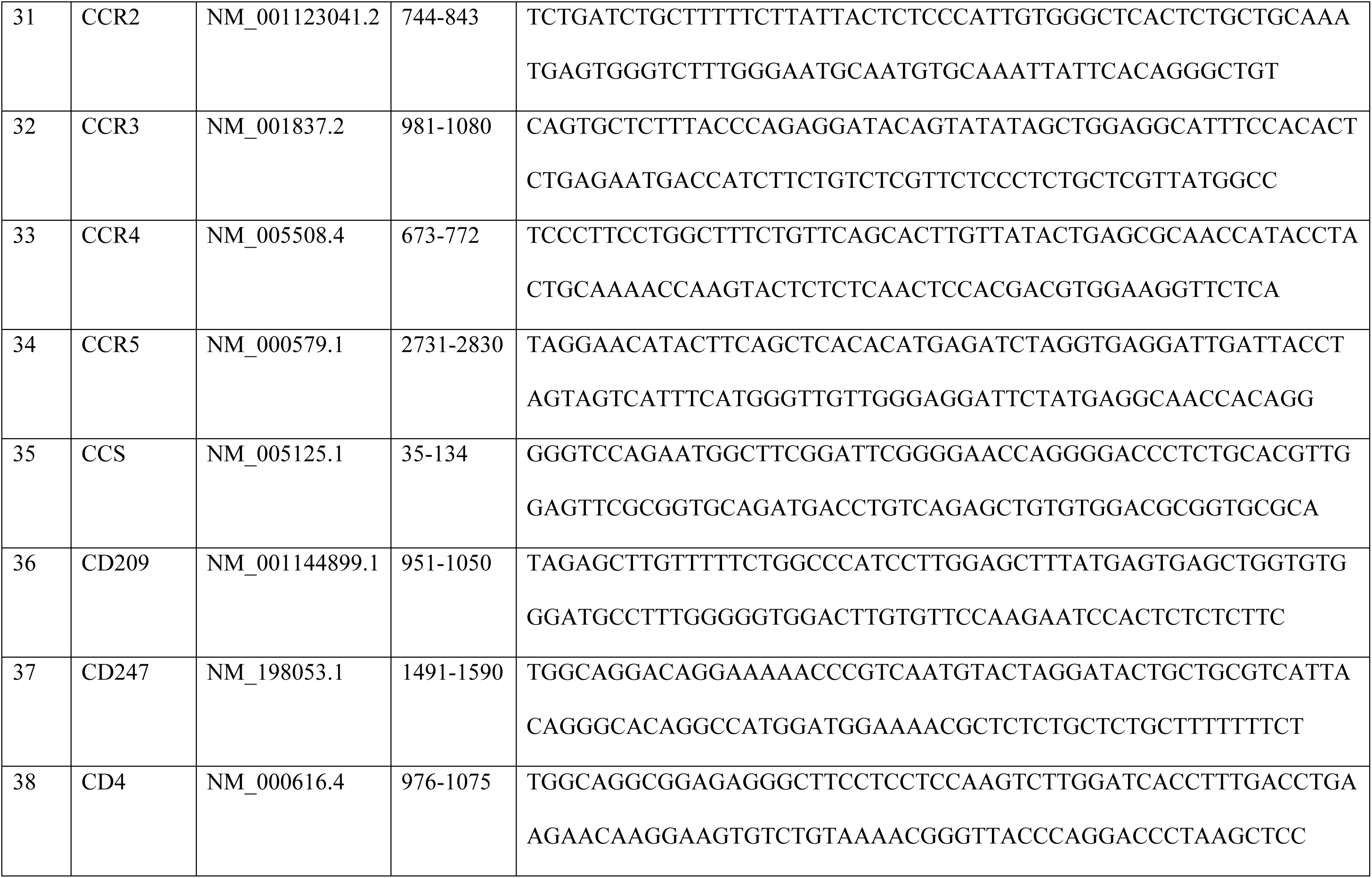
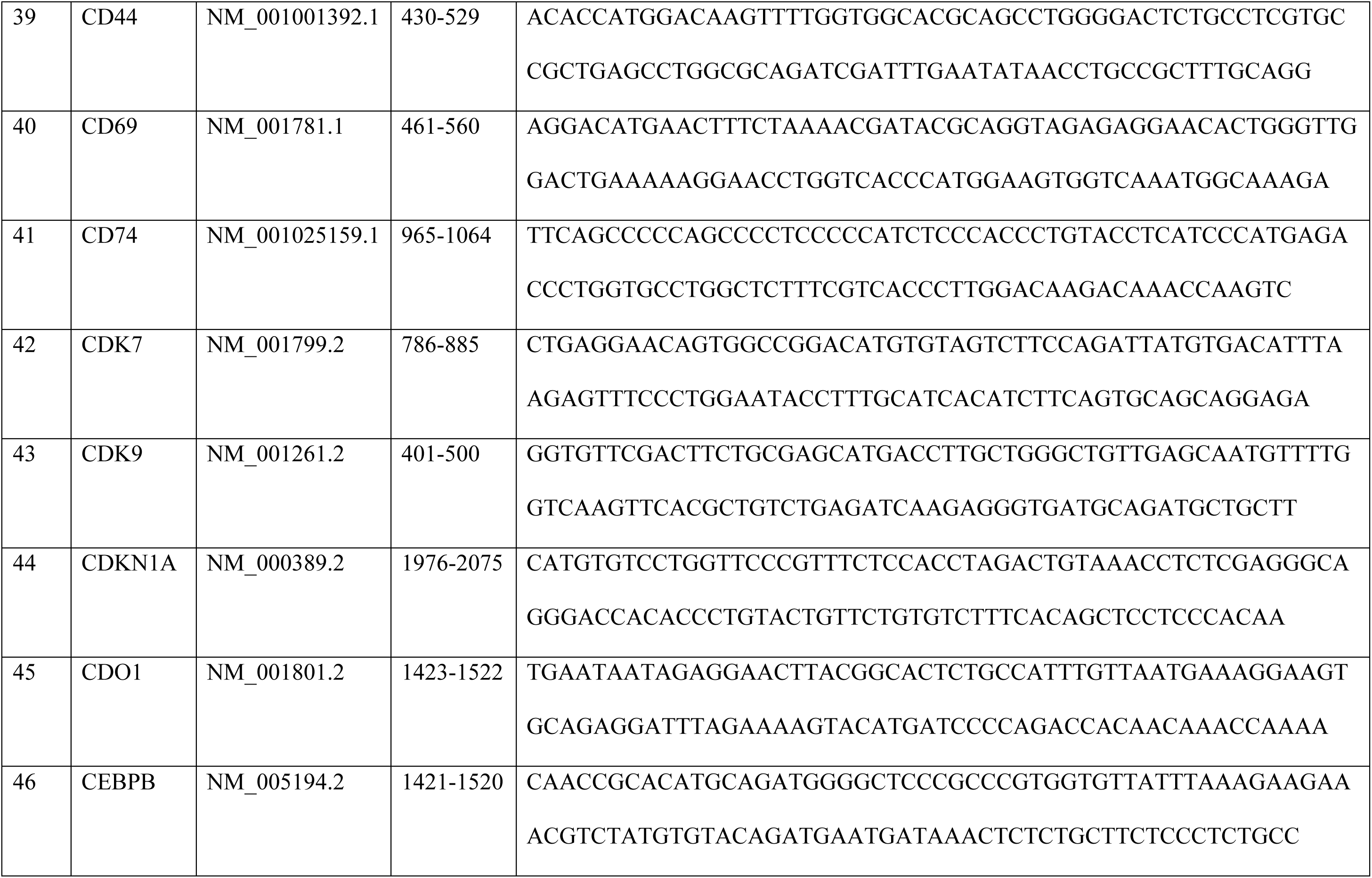
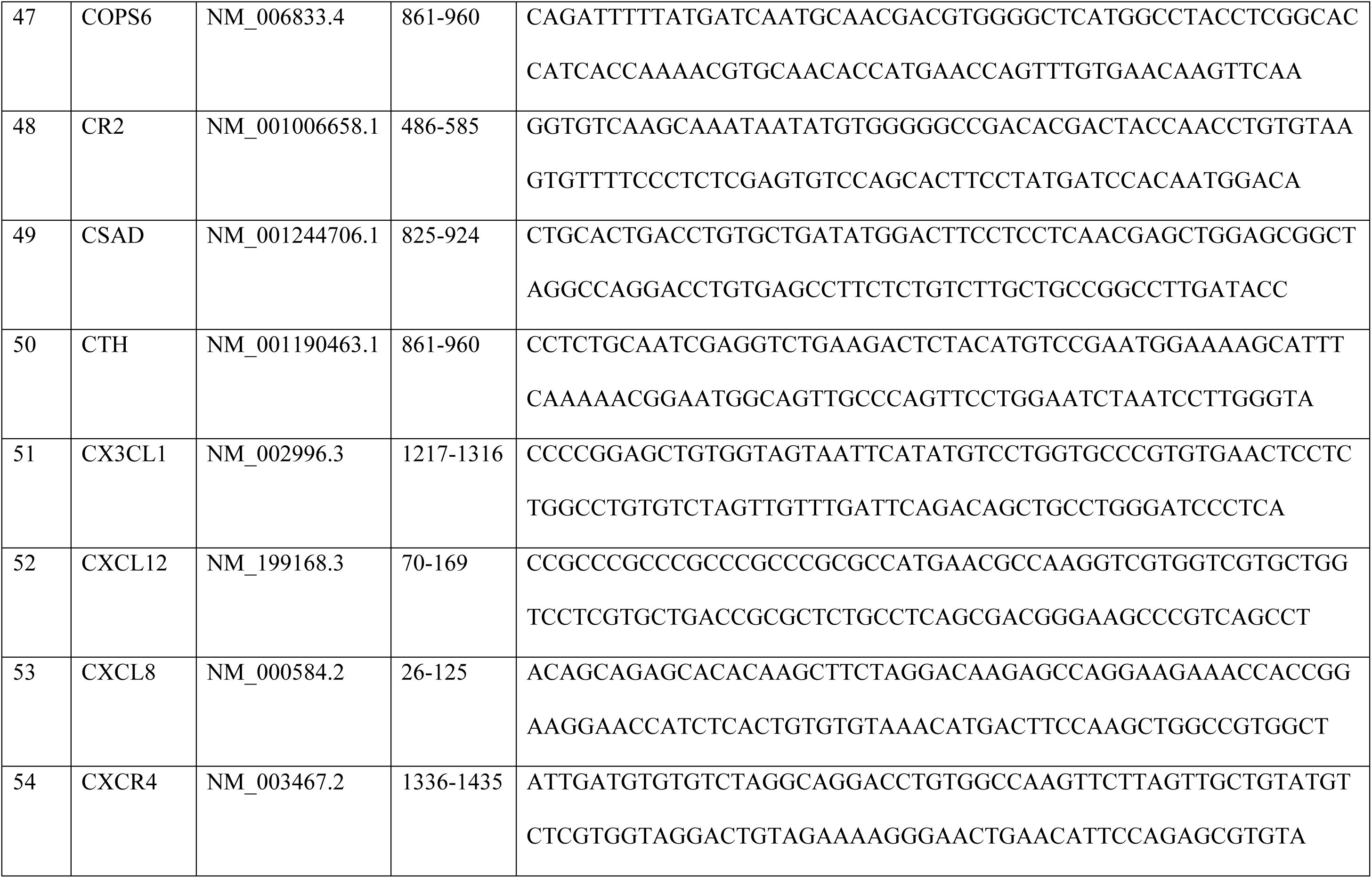
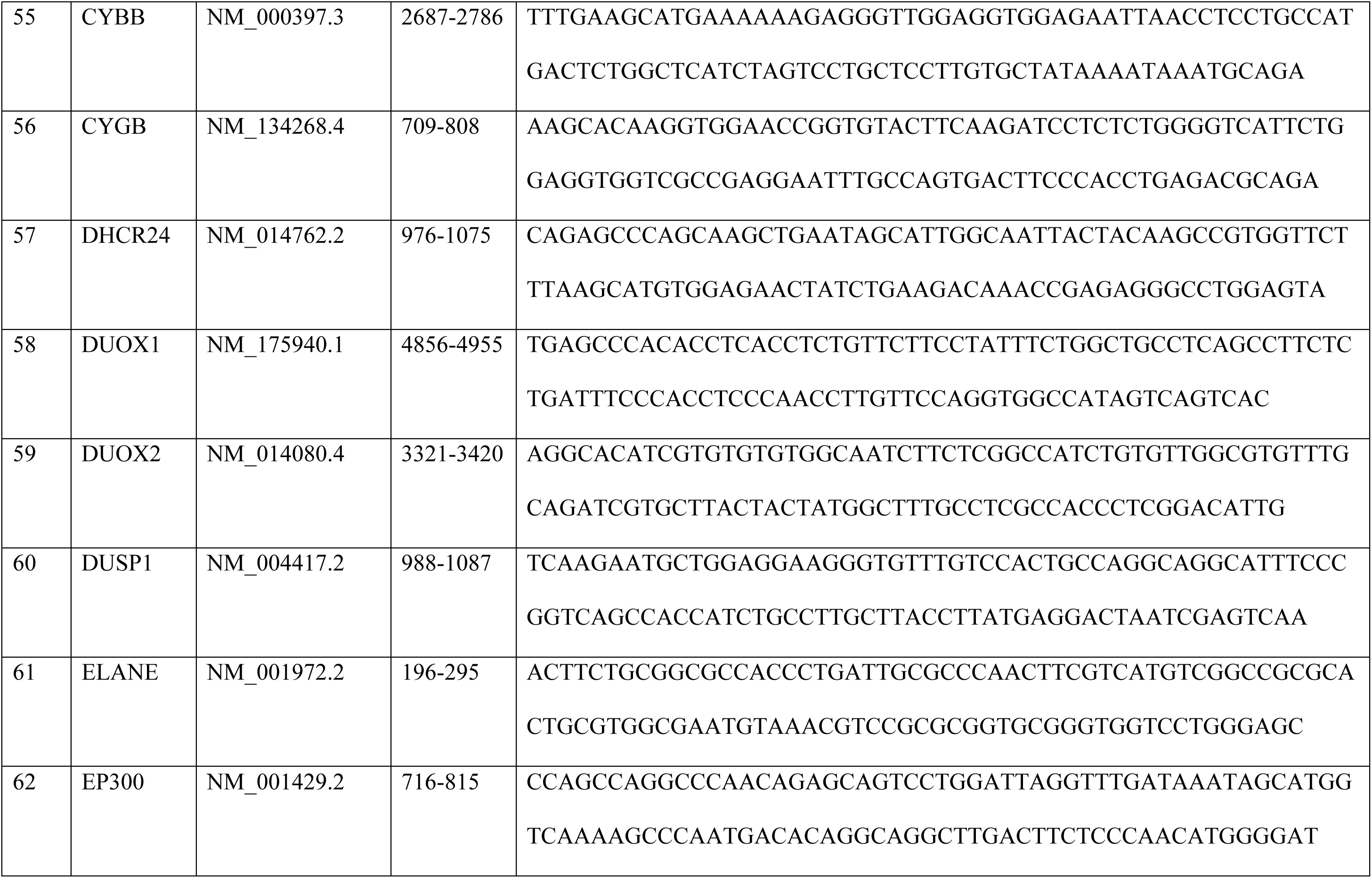
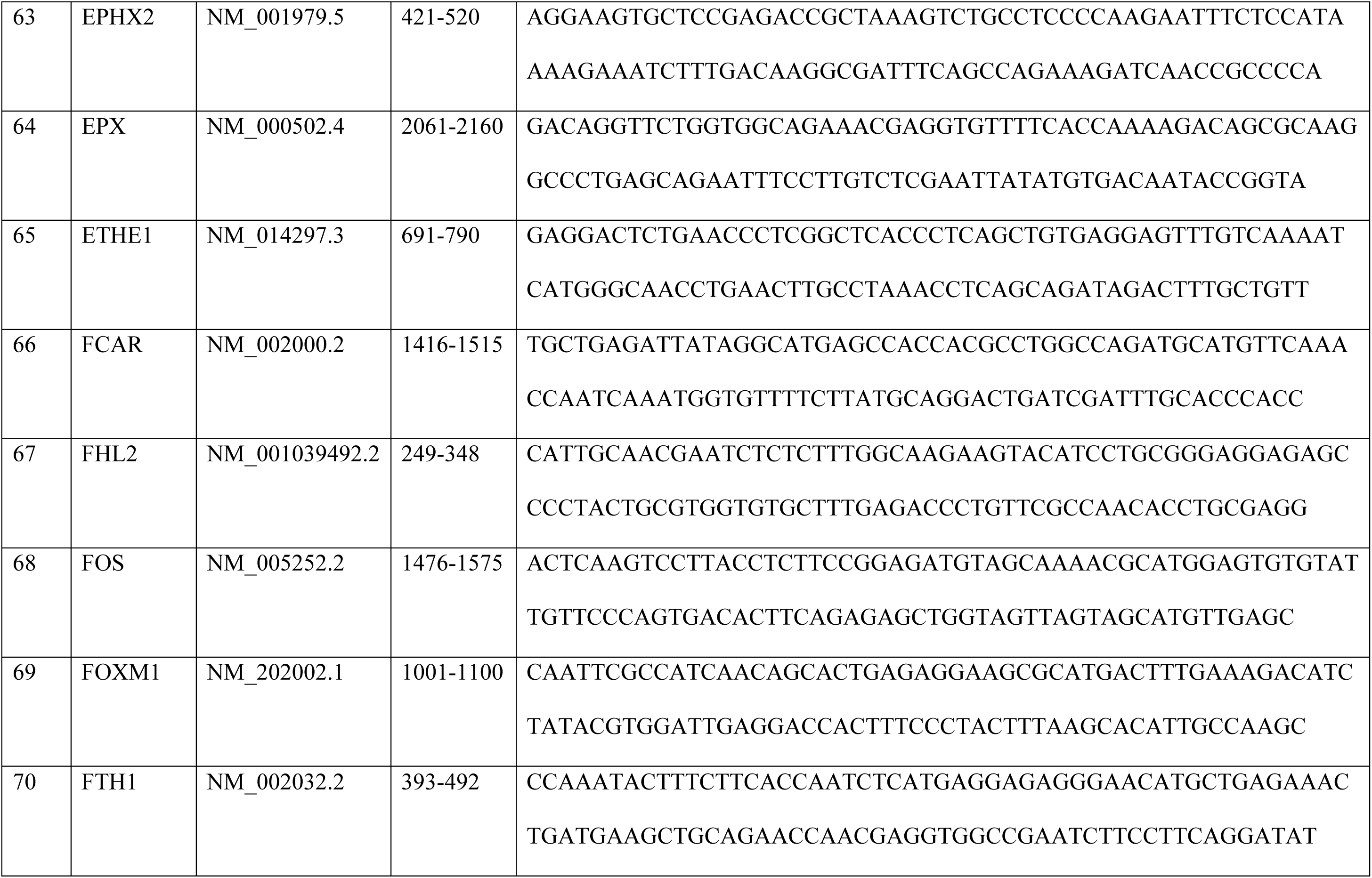
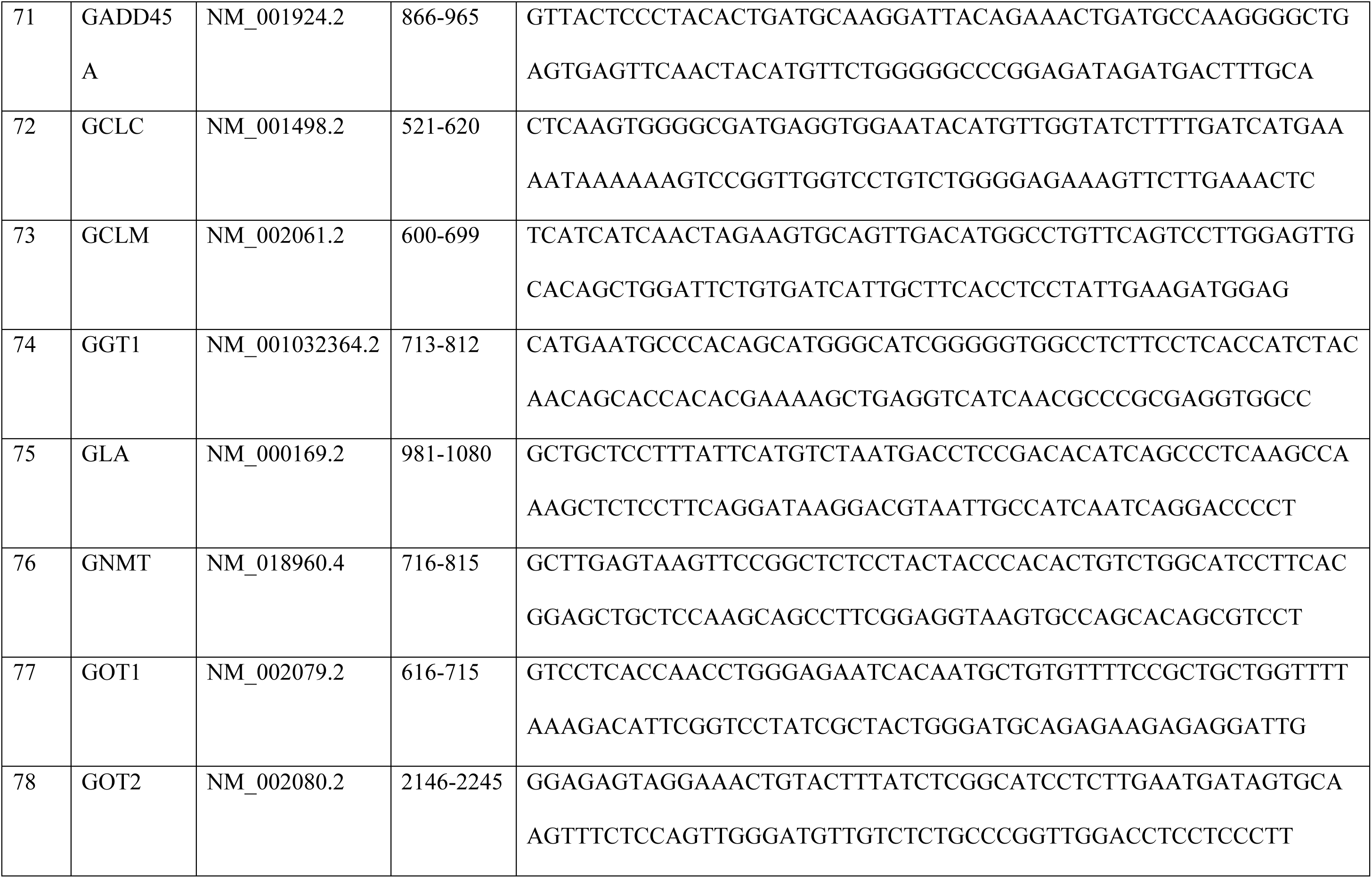
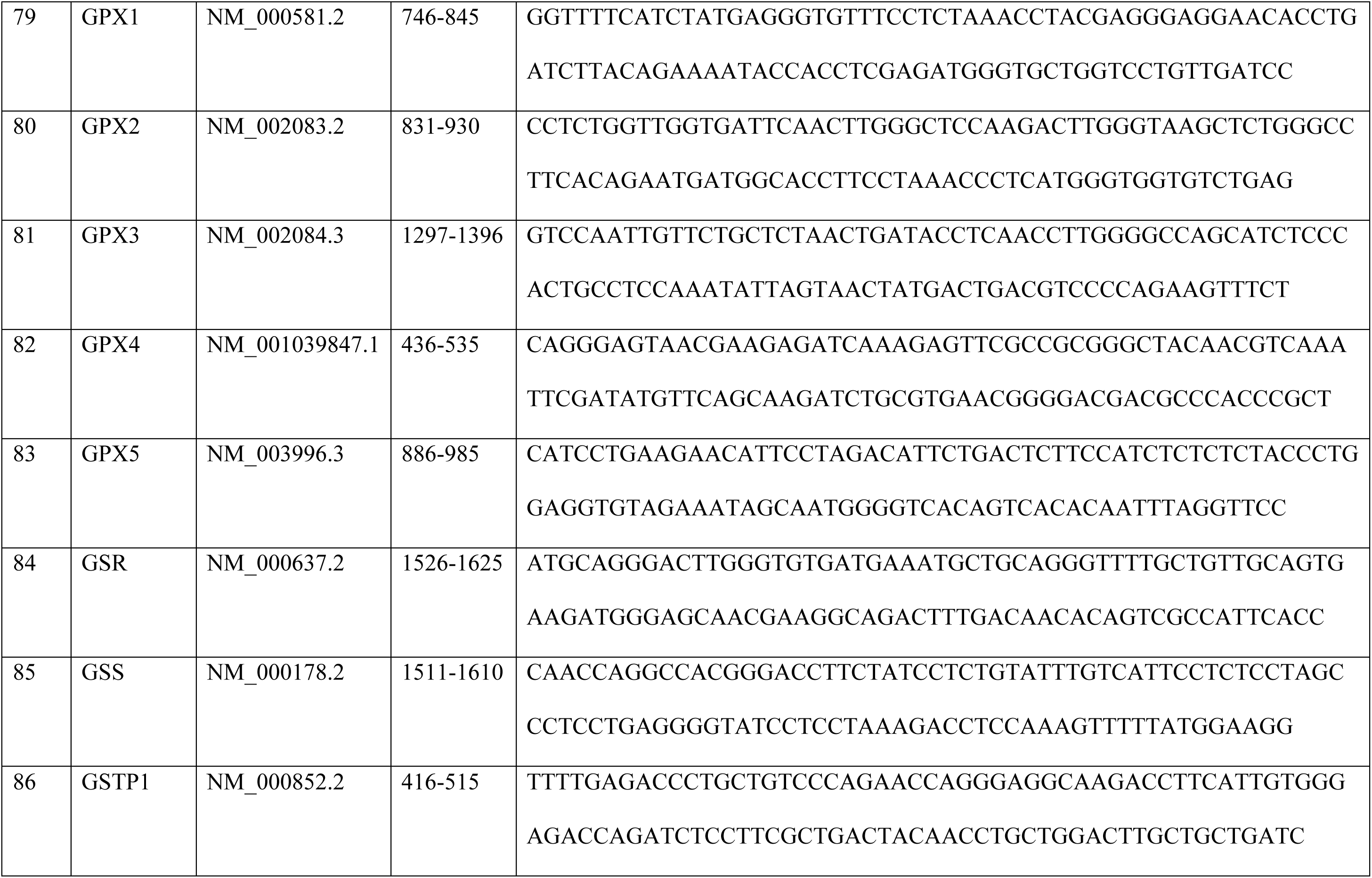
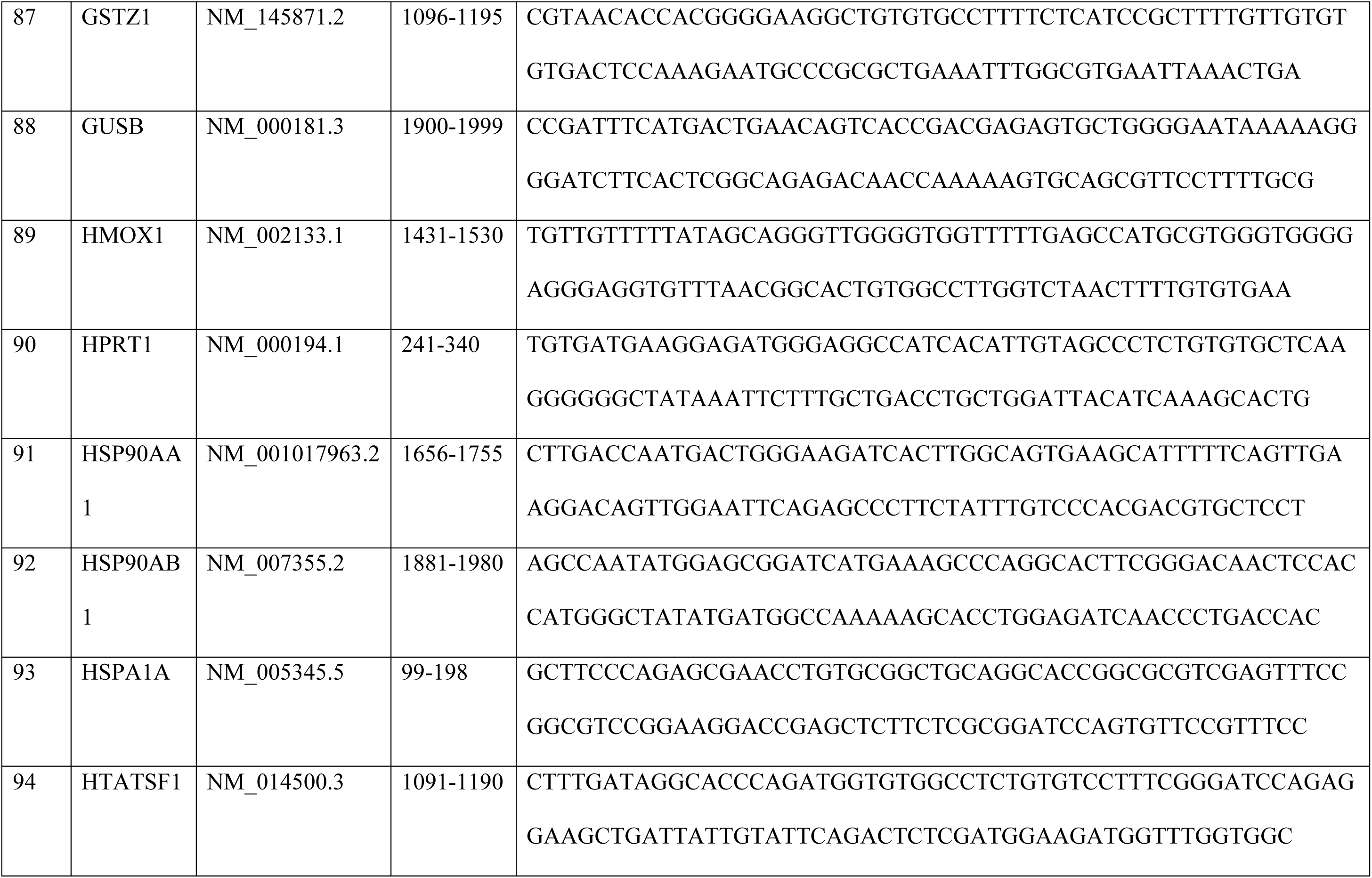
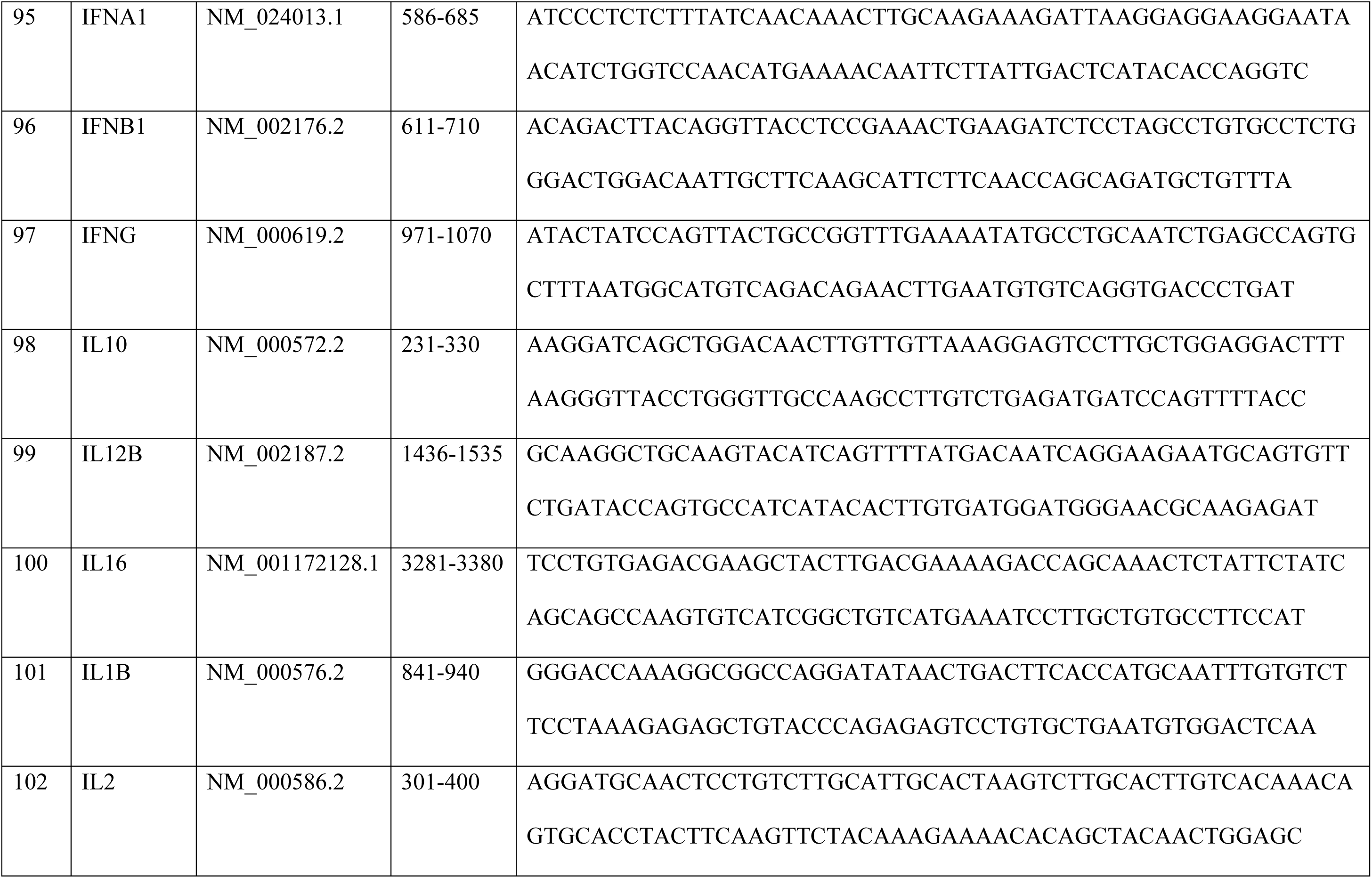
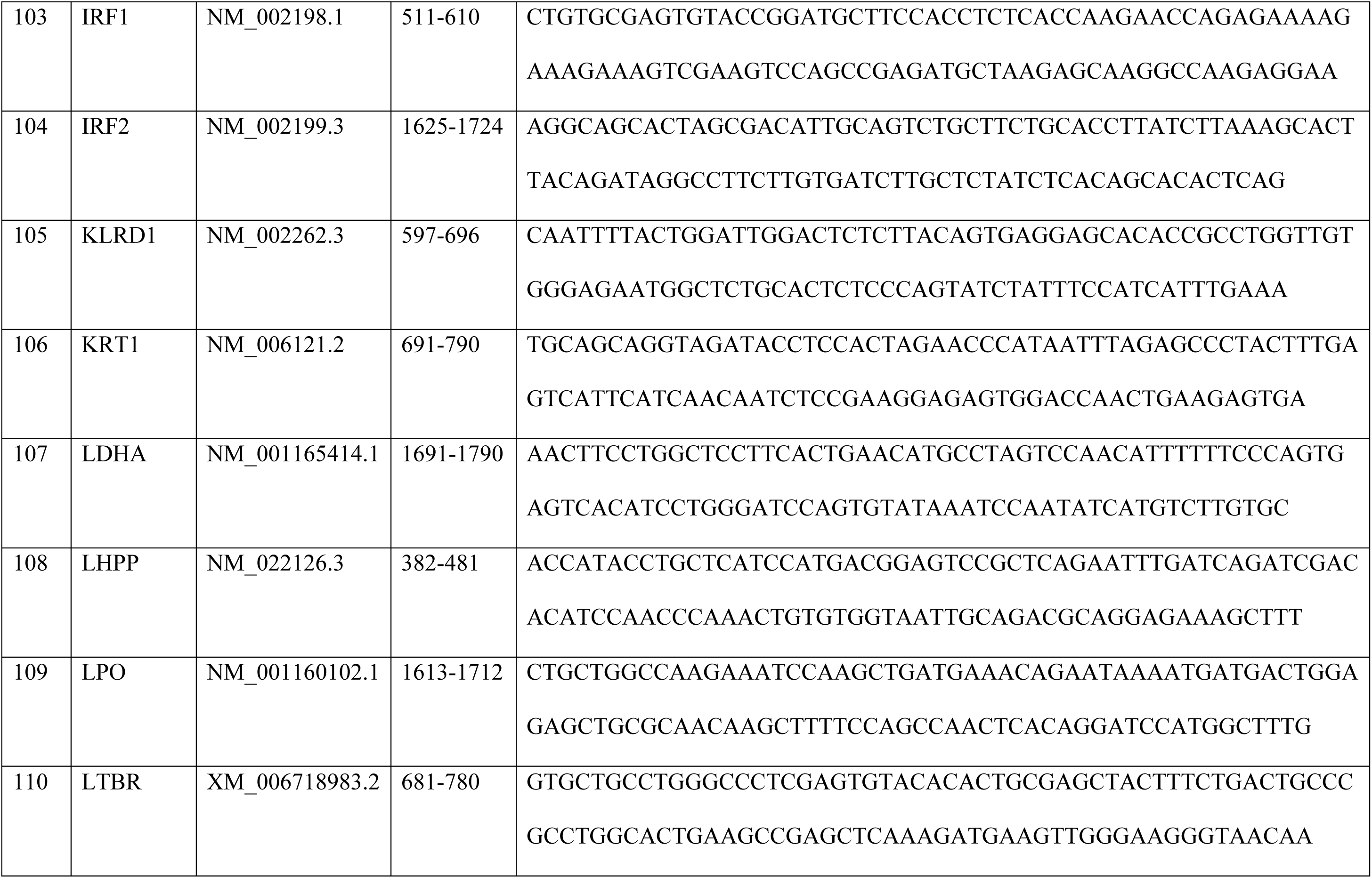
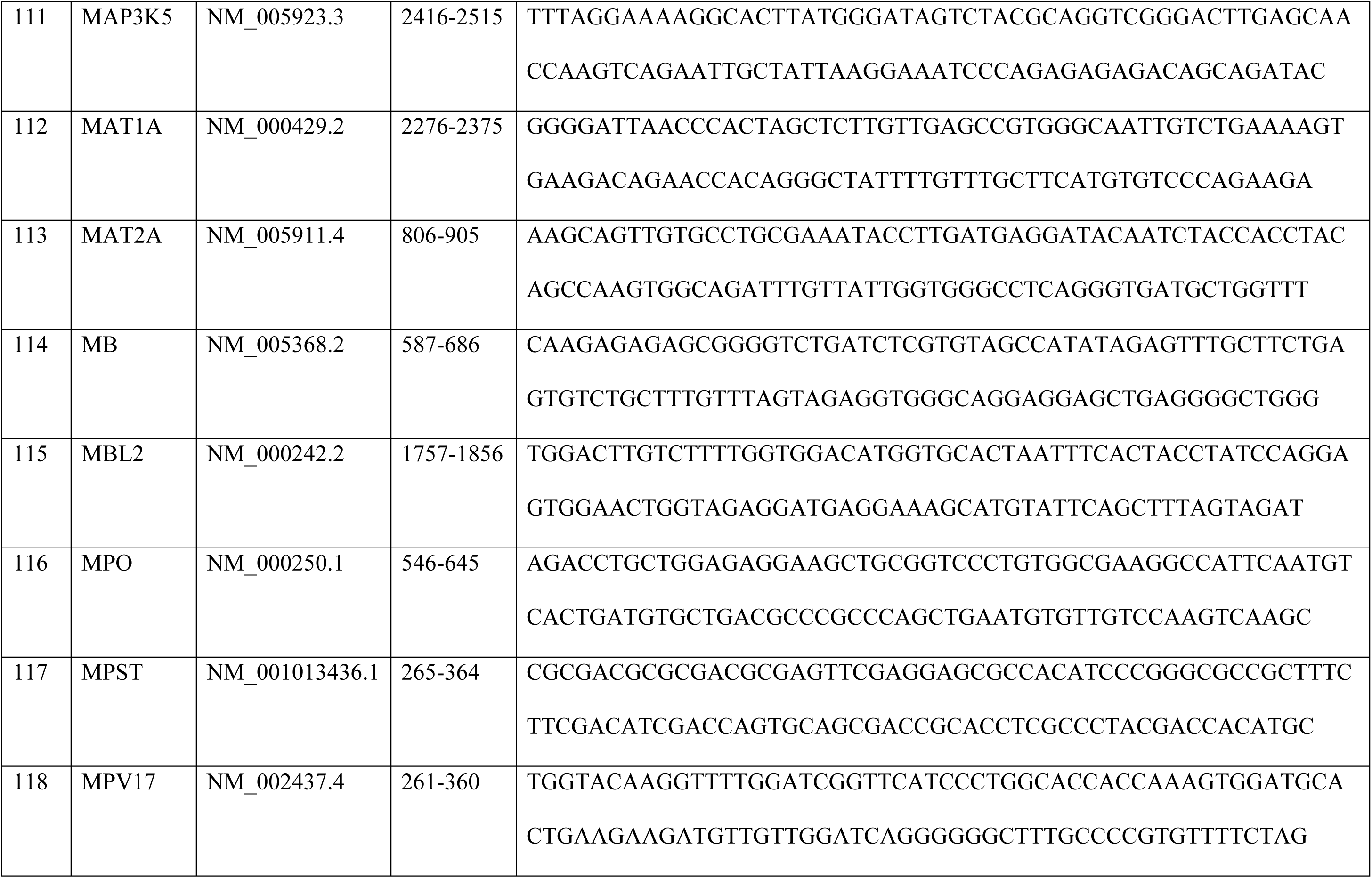
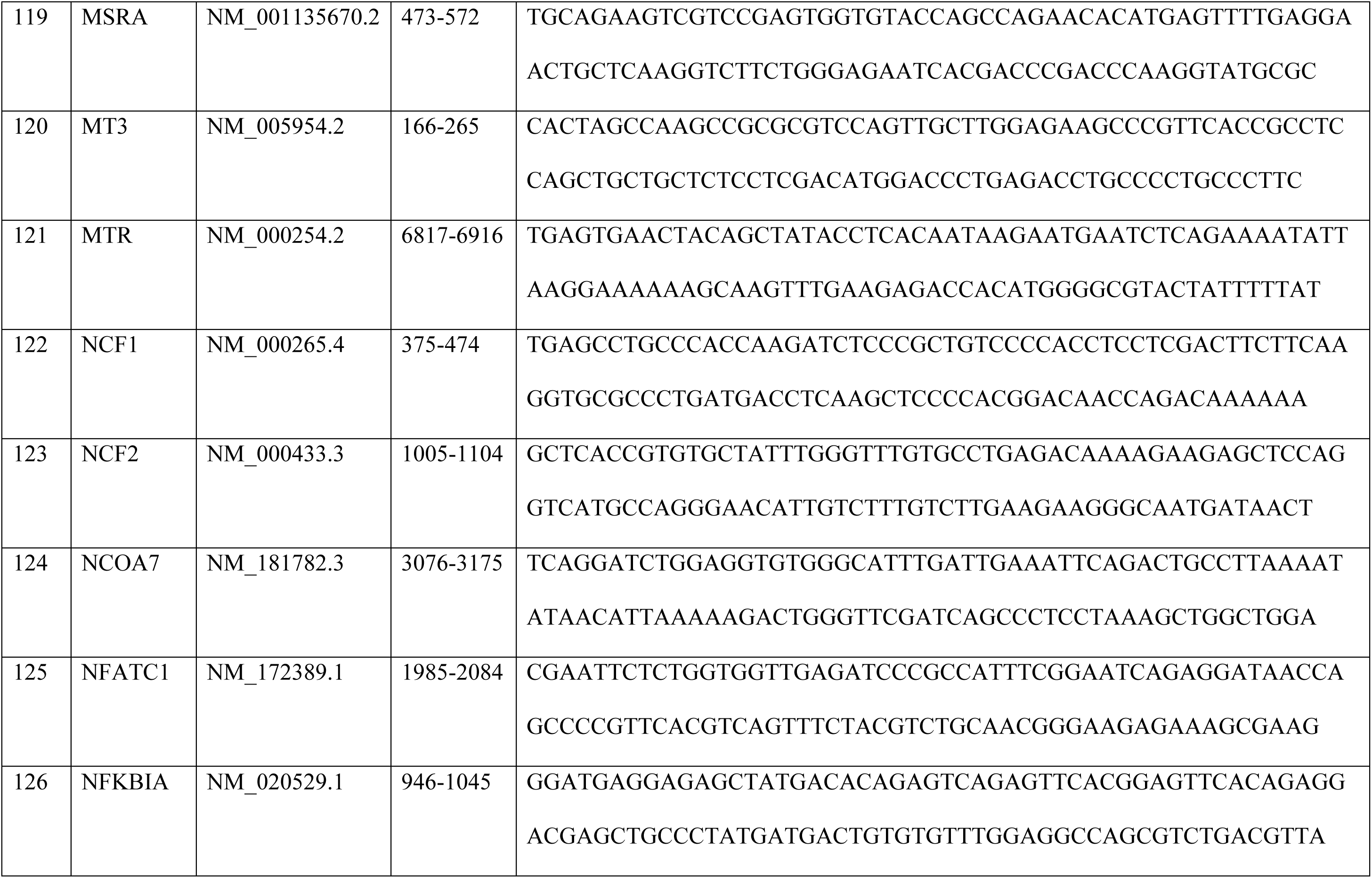
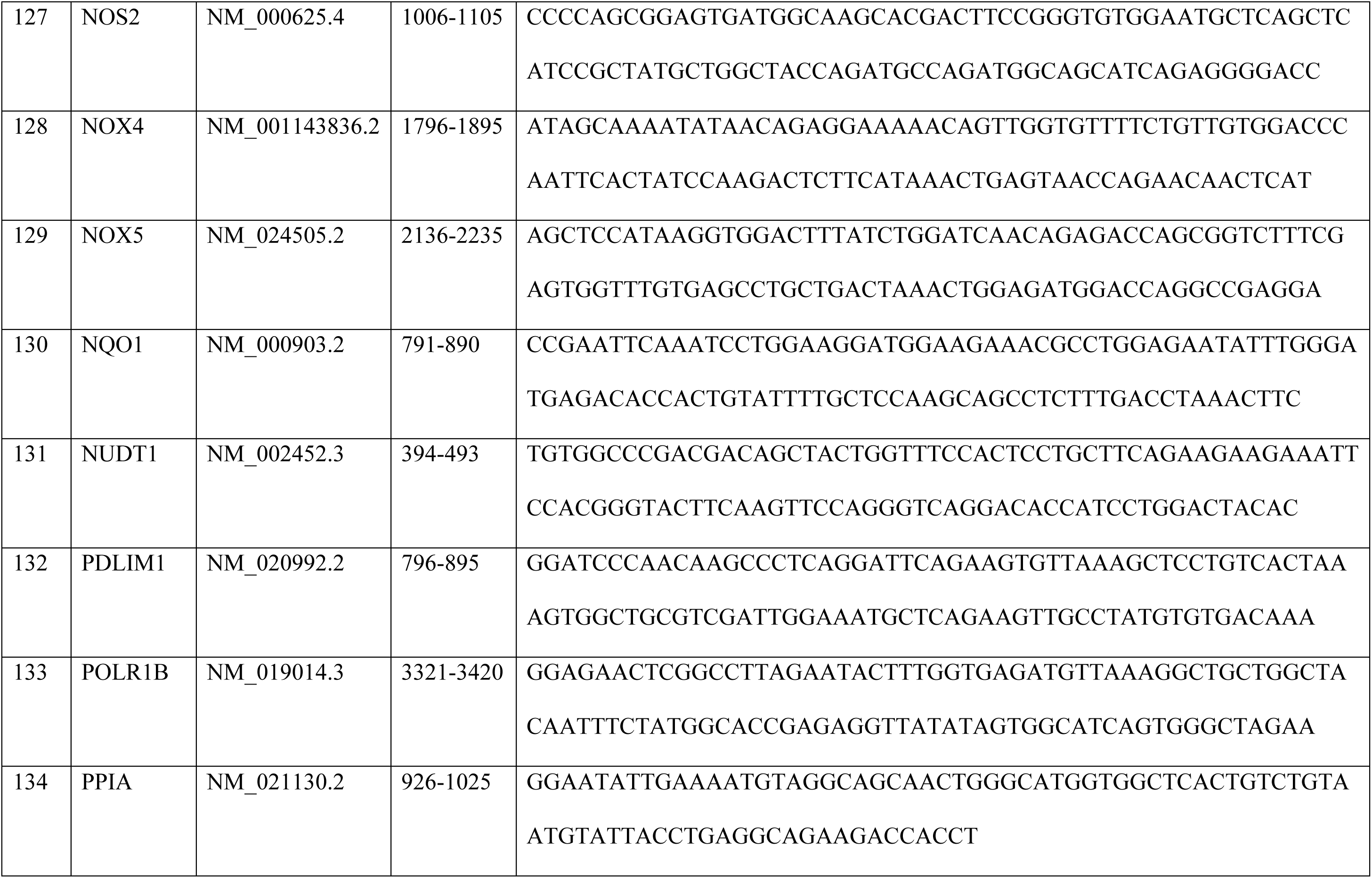
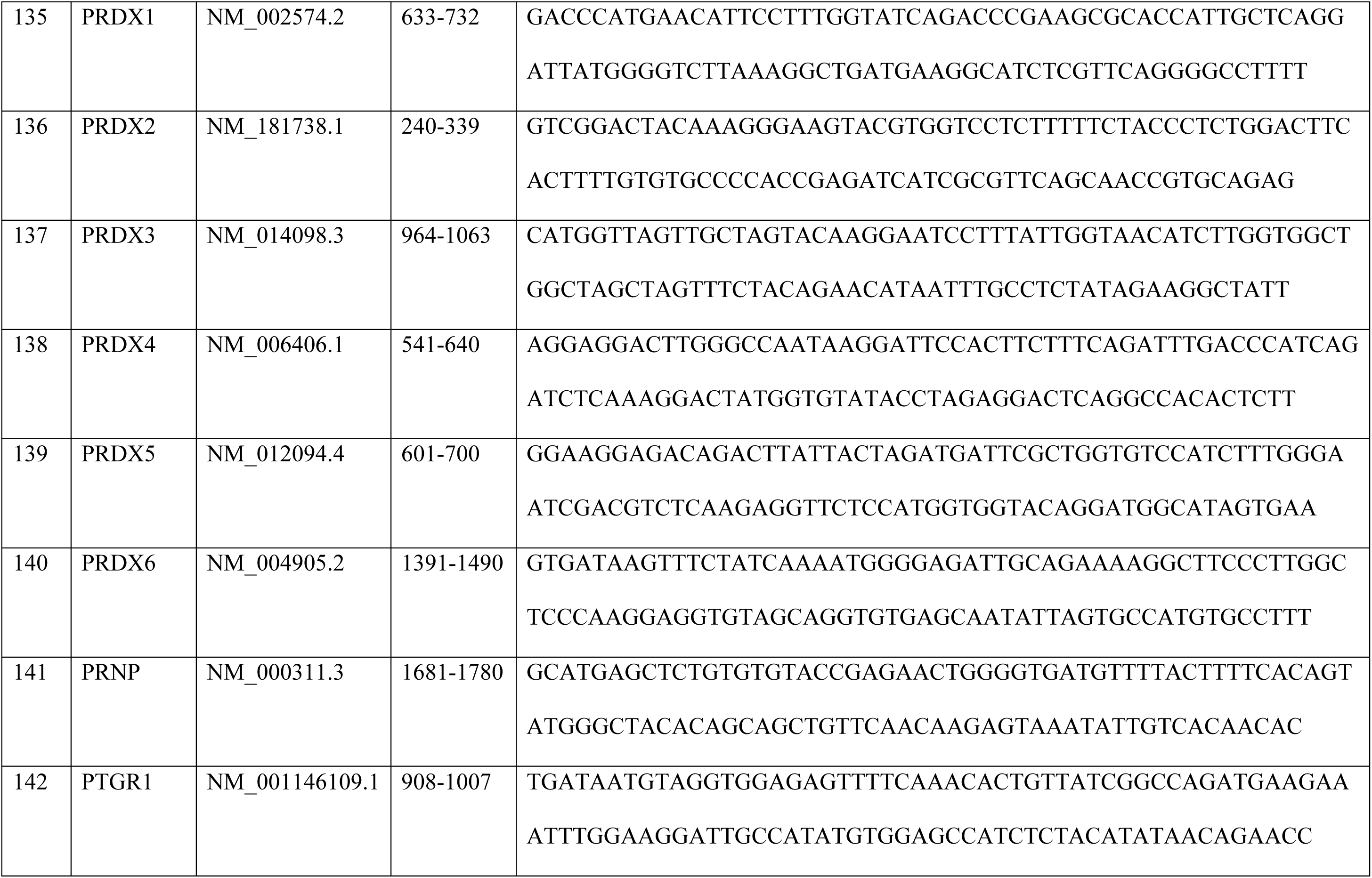
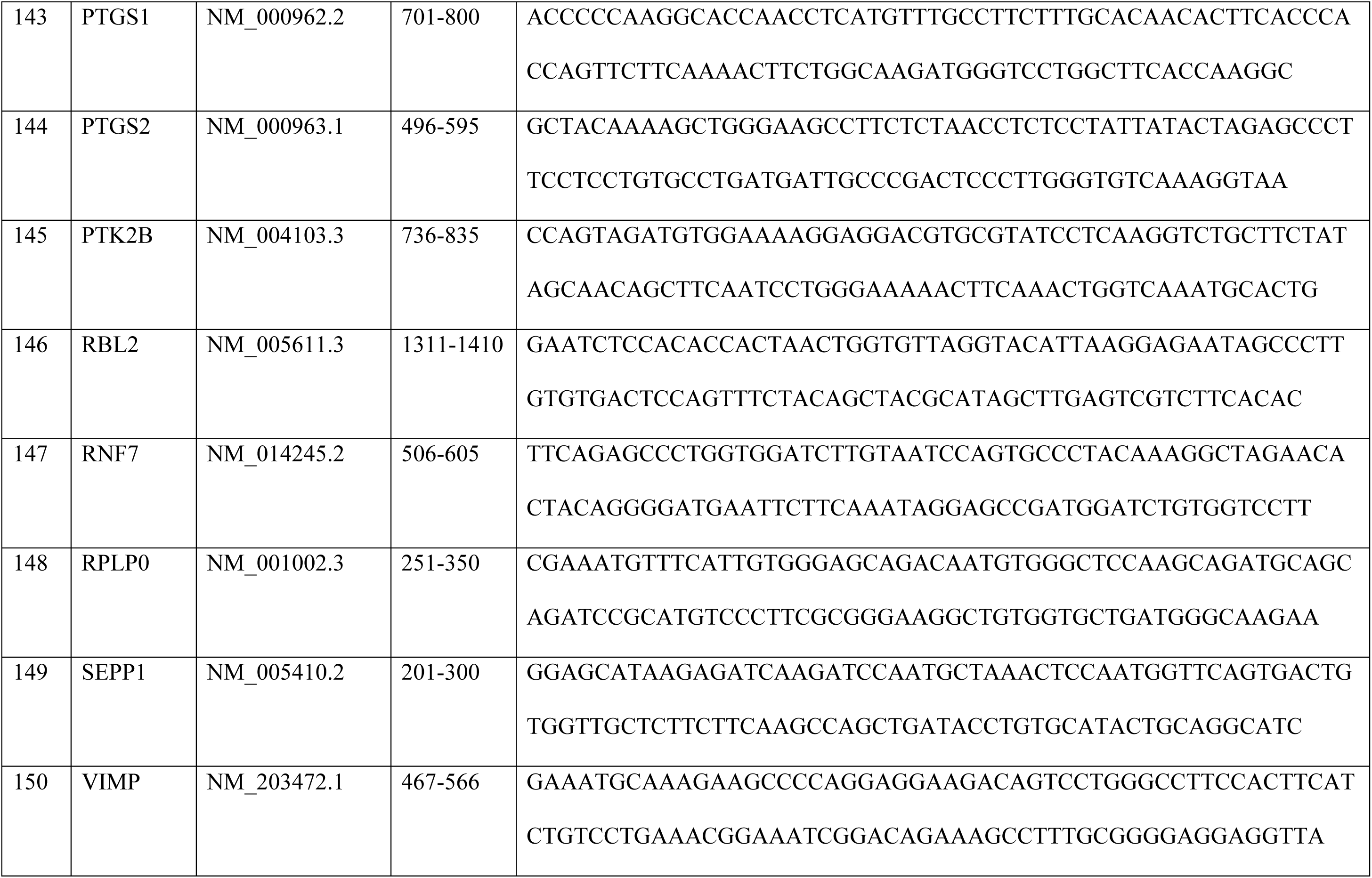
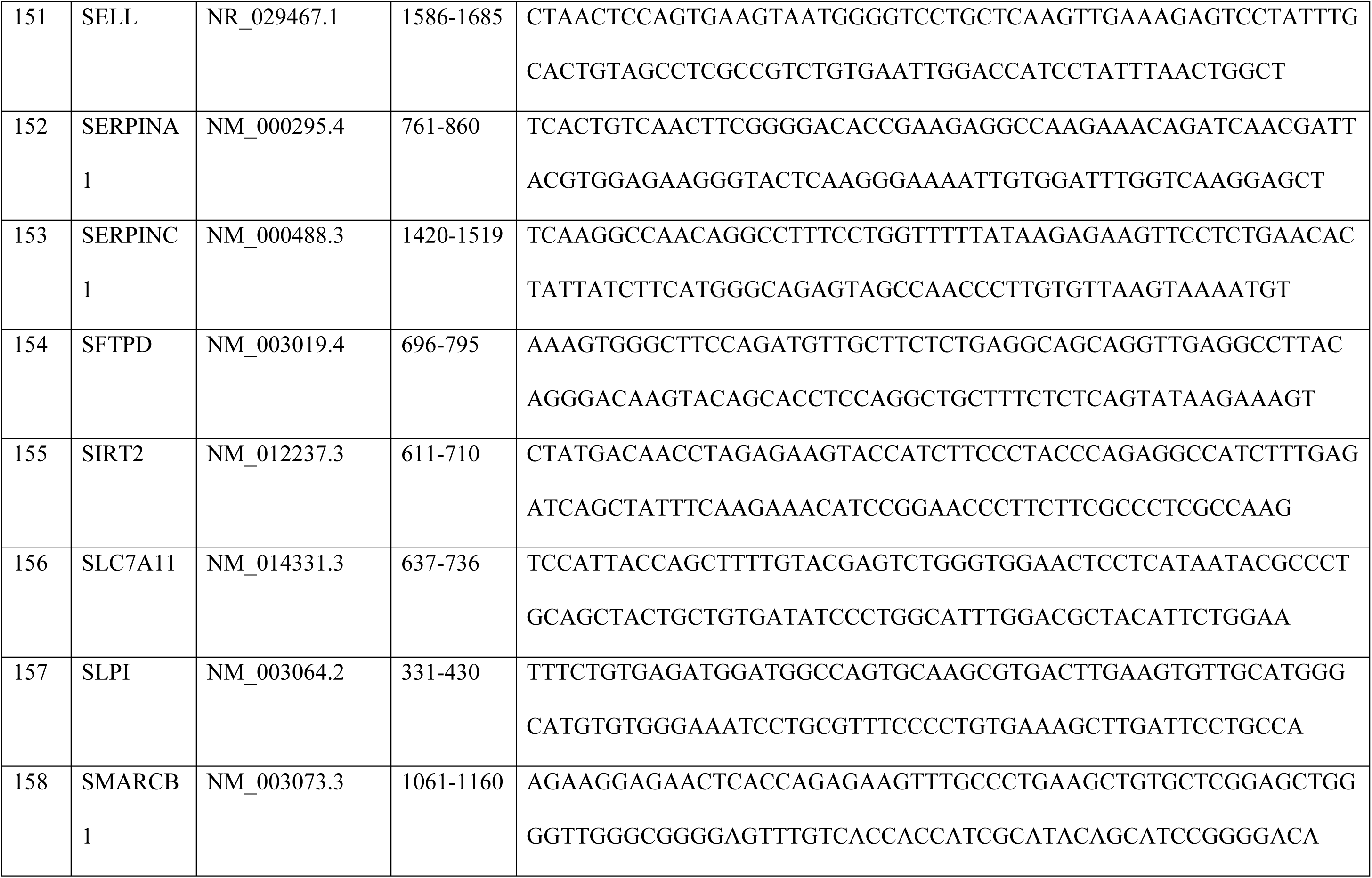
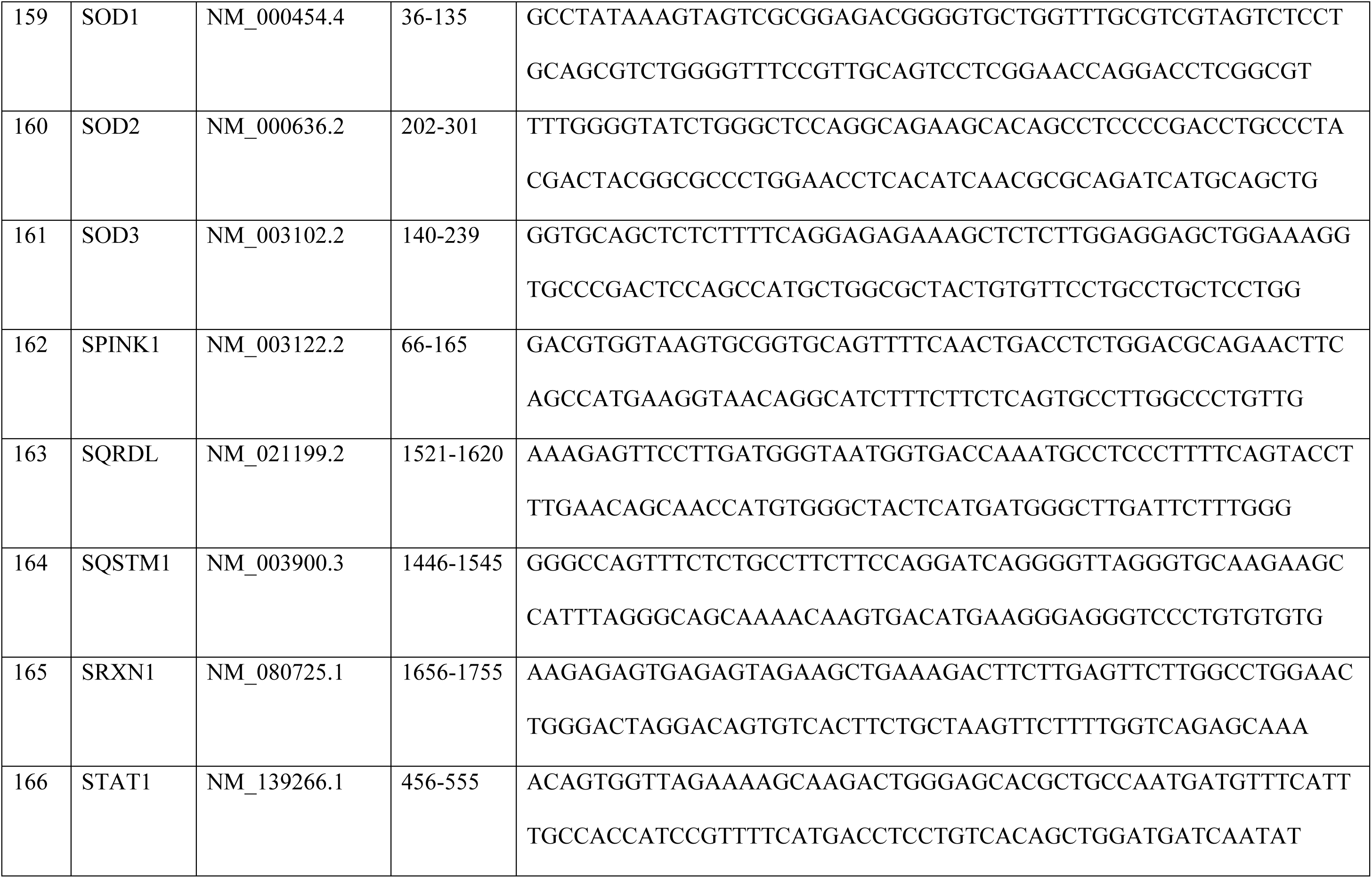
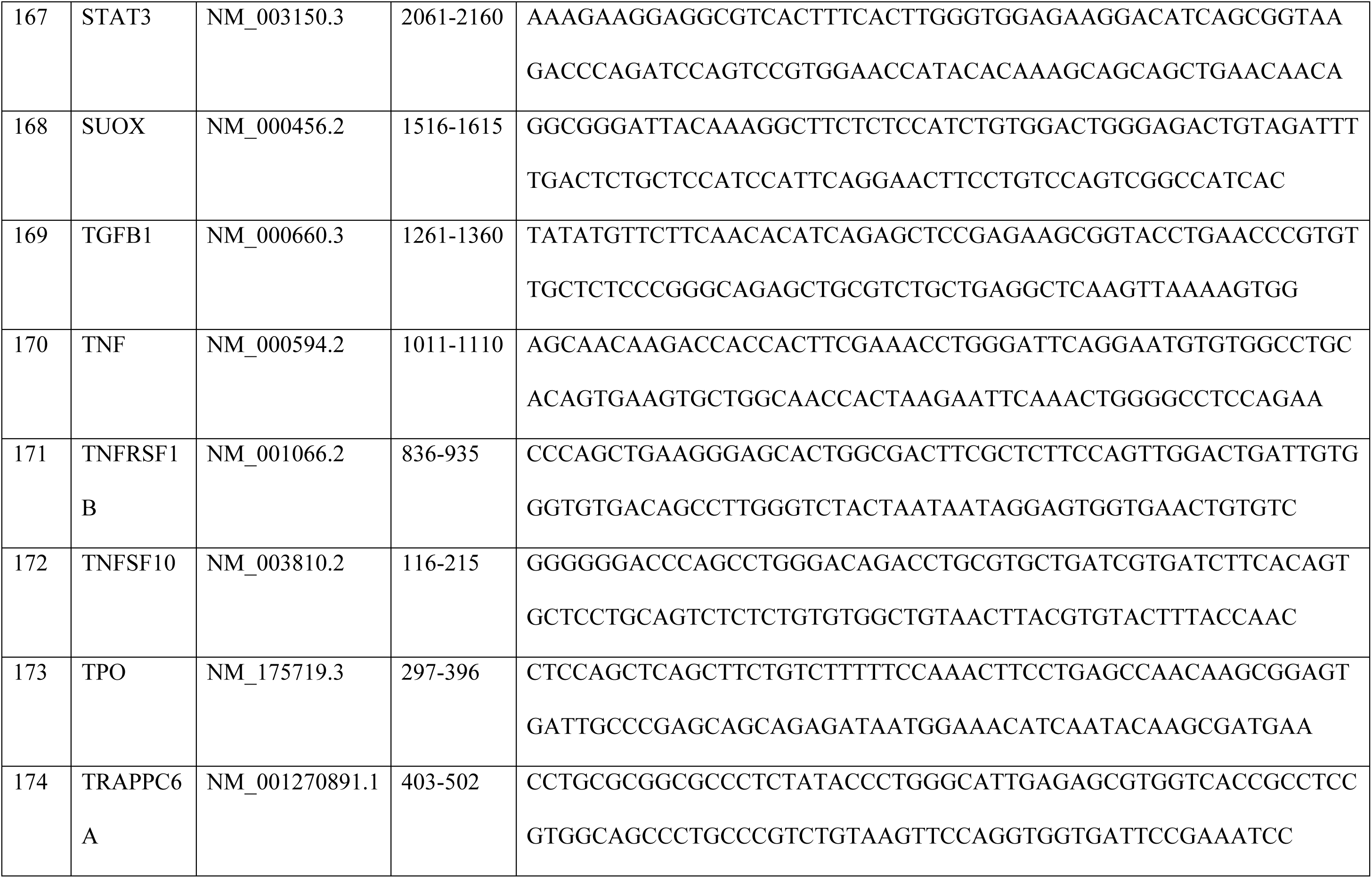
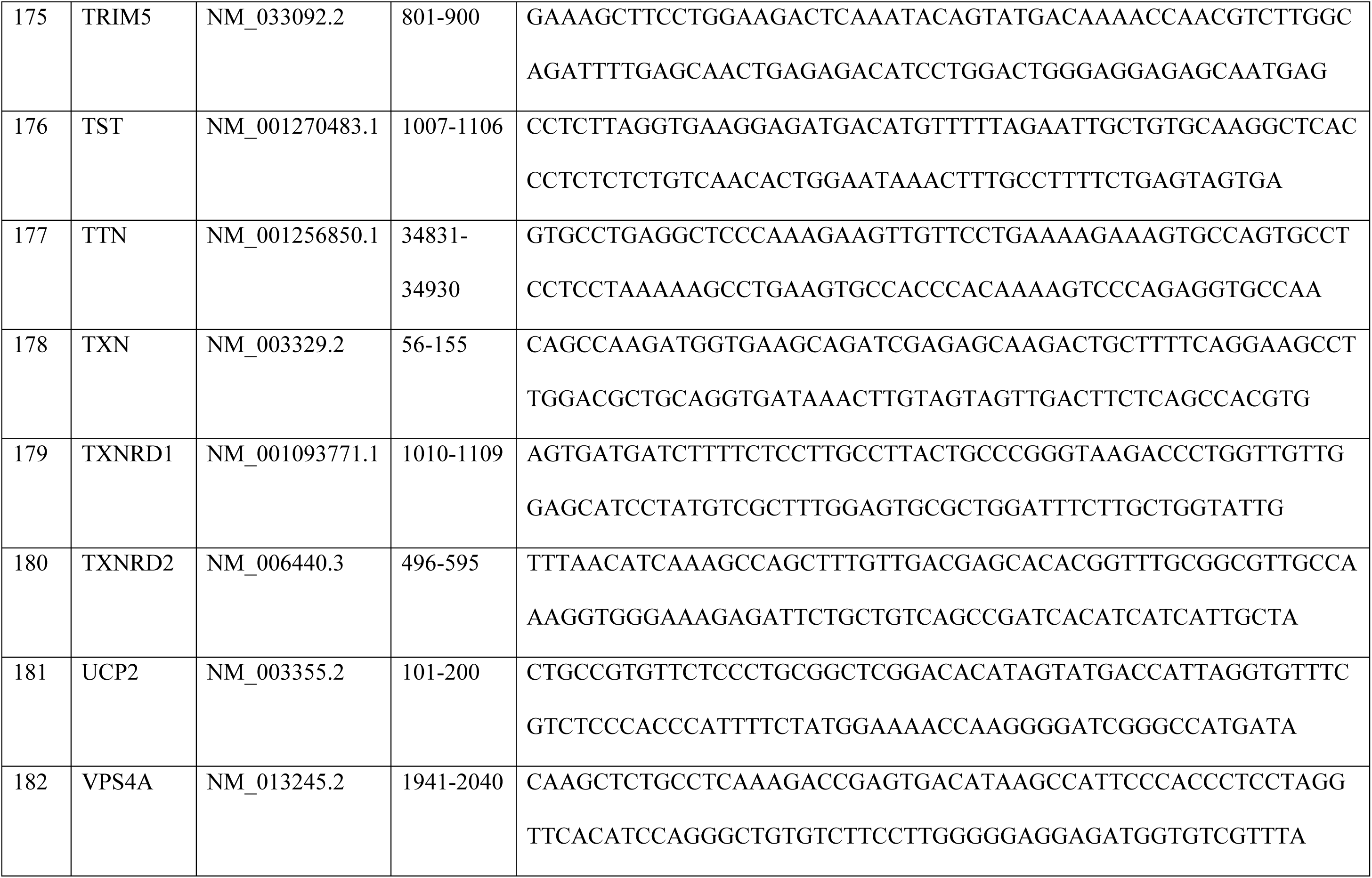
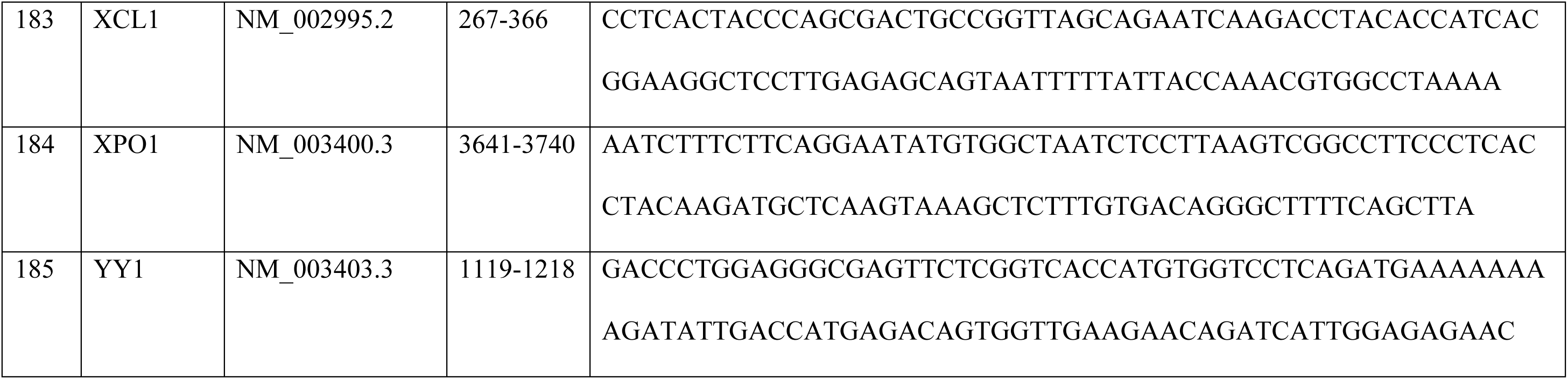
List of genes used for NanoString nCounter Gene Expression Analysis.

**Table S3.** List of differentially expressed genes from NanoString nCounter Gene Expression analysis for U1-shNT, shCTH, shNT+PMA and shCTH+PMA samples.

| Gene Name | shNT | shNT | shCTH | shCTH | shNT+PMA | shNT+PMA | shCTH+PMA | shCTH+PMA |
| --- | --- | --- | --- | --- | --- | --- | --- | --- |
| AHCY | 6800.61 | 6798.82 | 3007.09 | 3079.2 | 1541.84 | 1618.61 | 804.5 | 860.85 |
| APOBEC3G | 234.38 | 201 | 108.63 | 90.2 | 198.53 | 187.99 | 104.49 | 113 |
| ATOX1 | 2007 | 1951.48 | 1489.12 | 1382.74 | 895.87 | 941 | 675.16 | 735.96 |
| BAG2 | 699.43 | 740.18 | 499.9 | 483.13 | 178.78 | 223.11 | 152.72 | 167.26 |
| BANF1 | 3394.76 | 3101.1 | 2070.74 | 1902.73 | 1205.03 | 1173.42 | 780.38 | 832.61 |
| BAX | 3677.24 | 3480.76 | 1997.68 | 1939.6 | 2444.62 | 2646.38 | 1357.63 | 1471.93 |
| BNIP3 | 1022.62 | 969.89 | 7734.99 | 7048.59 | 522.51 | 467.92 | 5217.17 | 4695.3 |
| BTRC | 462.58 | 458.36 | 280.71 | 292.55 | 327.93 | 341.9 | 189.98 | 197 |
| CASP3 | 1773.86 | 1800.47 | 1415.1 | 1273.72 | 1021.31 | 1054.63 | 827.15 | 832.61 |
| CAT | 11705.25 | 11386.66 | 6447.75 | 6002.32 | 1934.96 | 1907.83 | 1090.93 | 1128.48 |
| CBX5 | 2396.81 | 2510.87 | 1287.24 | 1253.33 | 974.89 | 1005.05 | 558.98 | 523.35 |
| CCNT1 | 1375.42 | 1209.17 | 823.87 | 795.29 | 802.03 | 790.2 | 430.38 | 470.57 |
| CCS | 414.48 | 363.71 | 283.6 | 267.45 | 277.55 | 319.18 | 189.25 | 180.65 |
| CD44 | 2578.14 | 2314.13 | 1581.41 | 1450.97 | 2638.22 | 2492.47 | 1390.51 | 1492.74 |
| CD69 | 165.3 | 135.06 | 325.9 | 338.82 | 233.1 | 249.97 | 210.44 | 219.3 |
| CD74 | 1794.83 | 1626.06 | 1086.32 | 967.84 | 906.73 | 887.29 | 507.83 | 544.91 |
| CDK9 | 689.56 | 611.5 | 461.45 | 457.25 | 407.93 | 426.6 | 251.36 | 266.14 |
| COPS6 | 10065.84 | 9760.6 | 7308.15 | 6926.23 | 5343.6 | 5275.2 | 3573.83 | 3540.06 |
| CXCL8 | 592.11 | 505.15 | 8366.59 | 8854.85 | 12612.29 | 12636.94 | 9282.76 | 8908.13 |
| DHCR24 | 527.96 | 496.64 | 1047.87 | 1106.66 | 396.08 | 379.09 | 645.2 | 637.09 |
| FHL2 | 176.4 | 178.66 | 129.78 | 138.04 | 166.93 | 171.47 | 74.53 | 83.26 |
| GADD45A | 189.97 | 173.35 | 330.7 | 330.98 | 73.09 | 71.27 | 65.76 | 73.6 |
| GCLC | 1615.96 | 1501.63 | 825.8 | 839.21 | 1225.77 | 1263.28 | 938.21 | 918.1 |
| GCLM | 1486.44 | 1451.65 | 1045.94 | 1017.25 | 1027.24 | 1013.31 | 453.76 | 481.72 |
| GGT1 | 378.7 | 352.01 | 191.31 | 164.7 | 179.77 | 174.57 | 94.99 | 86.98 |
| GOT2 | 2367.2 | 2204.59 | 1317.04 | 1279.21 | 918.59 | 892.46 | 421.61 | 457.93 |
| GPX4 | 8558.43 | 7968.64 | 4669.26 | 4564.68 | 7274.61 | 7450.57 | 4099.21 | 4332.52 |
| GSR | 3815.4 | 3477.57 | 2787.9 | 2797.63 | 2851.57 | 2673.24 | 2004.3 | 2227.22 |
| GSS | 948.61 | 847.59 | 610.45 | 583.53 | 405.96 | 448.29 | 319.31 | 356.83 |
| GSTP1 | 33364.08 | 32080.89 | 18260.76 | 16725.39 | 8306.79 | 8218.04 | 4798.48 | 5133.9 |
| HTATSF1 | 1079.36 | 1024.13 | 738.31 | 683.13 | 435.59 | 497.88 | 273.28 | 315.2 |
| IRF2 | 843.75 | 776.34 | 472.98 | 465.1 | 904.76 | 854.24 | 554.6 | 556.06 |
| LHPP | 794.41 | 762.51 | 367.23 | 370.98 | 353.61 | 365.66 | 199.48 | 206.66 |
| LTBR | 767.27 | 713.59 | 462.41 | 417.25 | 559.05 | 569.15 | 283.51 | 321.89 |
| MAP3K5 | 181.33 | 142.51 | 116.32 | 135.69 | 106.67 | 92.96 | 79.65 | 70.62 |
| MAT2A | 4873.79 | 3889.14 | 1902.5 | 1924.69 | 2268.81 | 2273.49 | 851.26 | 882.41 |
| MPST | 1873.78 | 1978.07 | 1003.64 | 1087.84 | 805.99 | 819.12 | 576.52 | 527.07 |
| MPV17 | 1560.45 | 1525.03 | 1034.41 | 1008.62 | 915.62 | 913.12 | 740.2 | 734.48 |
| MSRA | 1390.22 | 1337.86 | 893.09 | 896.47 | 1120.08 | 1184.78 | 685.39 | 695.82 |
| NUDT1 | 1611.03 | 1535.66 | 817.14 | 742.74 | 530.41 | 505.11 | 279.86 | 300.33 |
| PRDX1 | 7352.01 | 7163.59 | 5674.82 | 5528.6 | 5215.2 | 5379.53 | 4369.56 | 4820.93 |
| PRDX3 | 11113.14 | 10357.22 | 6808.25 | 6555.26 | 5538.19 | 5225.62 | 3347.32 | 3353.47 |
| PRDX4 | 3350.35 | 3272.32 | 2406.25 | 2286.26 | 964.02 | 1014.34 | 831.53 | 872.01 |
| PRDX5 | 4445.75 | 4316.66 | 3067.65 | 2908.22 | 2390.3 | 2459.42 | 1750.75 | 1813.89 |
| PRDX6 | 3735.22 | 3587.11 | 2431.24 | 2299.59 | 1469.74 | 1525.65 | 1392.71 | 1487.54 |
| SIRT2 | 1845.4 | 1779.2 | 1095.93 | 1036.07 | 1374.92 | 1362.44 | 959.4 | 893.56 |
| SMARCB1 | 1532.08 | 1426.12 | 1213.22 | 1103.52 | 768.45 | 774.7 | 512.22 | 579.11 |
| SOD1 | 321.96 | 363.71 | 226.88 | 214.9 | 186.68 | 198.32 | 151.25 | 171.72 |
| SRXN1 | 967.11 | 944.37 | 543.16 | 519.21 | 926.49 | 859.4 | 439.15 | 459.42 |
| STAT1 | 5331.44 | 4973.89 | 3300.3 | 3220.37 | 4678.86 | 4914.71 | 2753.99 | 2807.81 |
| TST | 3956.02 | 3622.21 | 1776.57 | 1690.97 | 1624.81 | 1631.01 | 930.91 | 950.06 |
| TXN | 21556.45 | 20985.62 | 12849.35 | 12049.34 | 13859.79 | 14205.97 | 8202.79 | 8715.59 |
| TXNRD1 | 5689.18 | 5314.2 | 3246.46 | 3265.86 | 5325.82 | 5596.45 | 2686.04 | 2700.02 |
| TXNRD2 | 891.86 | 893.32 | 574.88 | 563.92 | 466.21 | 471.02 | 303.97 | 288.44 |
| UCP2 | 1350.75 | 1216.62 | 719.09 | 676.86 | 469.17 | 481.35 | 320.04 | 354.6 |
| VPS4A | 1090.47 | 1042.21 | 682.56 | 669.8 | 616.34 | 693.1 | 415.77 | 413.33 |
| YY1 | 3268.93 | 3107.48 | 2516.8 | 2403.12 | 1877.67 | 1889.24 | 1402.21 | 1405.02 |

**Table S4.** List of differentially expressed genes from NanoString nCounter Gene Expression analysis for U1 untreated, GYY4137, PMA and GYY4137+PMA treated samples.

| Gene name | Untreated<br>U1 | Untreated<br>U1 | GYY4137 | GYY4137 | PMA | PMA | GYY4137+PMA | GYY4137+PMA |
| --- | --- | --- | --- | --- | --- | --- | --- | --- |
| SLC7A11 | 155.21 | 164.96 | 1603.4 | 1227.34 | 791.24 | 593.89 | 5890.68 | 6221.89 |
| SRXN1 | 480.44 | 511.08 | 1121.46 | 1220.36 | 718.73 | 614.32 | 2618.63 | 2663.93 |
| VIMP | 35.98 | 32.38 | 46.05 | 43.04 | 36.26 | 34.52 | 105.07 | 104.38 |
| TXNRD1 | 3171.21 | 3258.42 | 4192.18 | 3911.2 | 3123.37 | 2865.17 | 7959.31 | 7667.31 |
| GCLM | 1189.47 | 1172.48 | 1850 | 1640.33 | 670.74 | 630.52 | 1773.11 | 1565.68 |
| GCLC | 737.95 | 696.86 | 974.12 | 823.65 | 933.07 | 806.64 | 1310.13 | 1316.08 |
| FTH1 | 21411.12 | 20353.01 | 127359.45 | 102750.92 | 238445.8 | 221289 | 643242.69 | 631401.5 |
| HMOX1 | 395.78 | 336.09 | 4481.75 | 3622.68 | 2674.43 | 2214.22 | 10581.22 | 11129.97 |
| GSR | 2162.35 | 2208.51 | 3238.52 | 3362.09 | 3398.49 | 3150.49 | 5964.56 | 6169.7 |
| GPX4 | 5594.59 | 5173.24 | 5606.28 | 4895.39 | 4513.91 | 4165.66 | 7481.55 | 7810.26 |
| DUSP1 | 47.97 | 40.86 | 77.77 | 51.19 | 246.33 | 176.83 | 577.9 | 599.04 |
| PRDX1 | 6725.5 | 6387.34 | 3107.55 | 3285.31 | 7922 | 7614.85 | 4109.35 | 4440.64 |
| PRDX3 | 6118.77 | 5924.83 | 3687.72 | 3491.23 | 3234.27 | 3194.87 | 1937.29 | 1765.36 |
| PRDX5 | 2971.55 | 2990.94 | 2560.12 | 2715.27 | 1629.4 | 1596.38 | 1553.12 | 1477.19 |
| PRDX6 | 2177.87 | 2149.16 | 1154.2 | 1151.72 | 1869.33 | 1651.33 | 981.78 | 1066.48 |
| GPX1 | 1494.24 | 1239.54 | 1951.3 | 1676.39 | 1647.53 | 1423.07 | 620.59 | 710.23 |
| PRNP | 1113.27 | 1190.98 | 1645.35 | 1504.22 | 1259.37 | 1144.8 | 3058.62 | 2820.5 |
| CCS | 219.41 | 227.4 | 304.92 | 261.75 | 287.92 | 226.85 | 469.55 | 494.67 |
| MSRA | 816.26 | 825.59 | 862.58 | 795.73 | 893.61 | 778.46 | 1318.34 | 1347.85 |
| UCP2 | 557.34 | 656.77 | 514.68 | 492.1 | 556.64 | 447.35 | 392.38 | 347.17 |
| ATOX1 | 1116.8 | 1091.54 | 690.68 | 665.44 | 635.55 | 554.43 | 508.95 | 467.44 |
| DHCR24 | 620.84 | 665.25 | 336.64 | 401.36 | 693.13 | 660.81 | 305.37 | 258.68 |
| NOS2 | 12.16 | 12.48 | 25.58 | 17.45 | 36.26 | 33.11 | 59.1 | 52.19 |
| CYBB | 294.9 | 336.09 | 400.08 | 375.76 | 1066.36 | 929.22 | 630.44 | 739.73 |
| NQO1 | 721.02 | 559.64 | 960.81 | 894.62 | 1336.15 | 1306.83 | 655.07 | 614.93 |
| NCF1 | 222.94 | 285.99 | 168.83 | 235 | 9386.11 | 7471.84 | 1494.01 | 1275.24 |
| CAT | 6398.15 | 7127.37 | 8670.86 | 9332.43 | 1282.83 | 1146.91 | 1666.4 | 1747.21 |
| GSTP1 | 16617.27 | 17221.78 | 10642.62 | 11077.46 | 6391.77 | 5419.66 | 3776.07 | 3982.28 |
| BCL2 | 249.75 | 277.51 | 311.06 | 241.98 | 132.23 | 133.85 | 88.66 | 95.3 |
| FHL2 | 132.63 | 137.98 | 35.81 | 36.06 | 121.57 | 90.17 | 57.46 | 56.73 |
| CDKN1A | 227.88 | 194.26 | 539.24 | 415.32 | 1884.26 | 1800.68 | 3524.88 | 3526.19 |
| LTBR | 506.55 | 504.14 | 718.31 | 631.7 | 545.98 | 414.24 | 751.93 | 782.84 |
| SQSTM1 | 2445.96 | 2482.17 | 8485.65 | 7189.53 | 7390.95 | 6768.05 | 19671.71 | 20338 |
| CTH | 326.64 | 295.24 | 814.49 | 685.22 | 146.09 | 130.33 | 2426.54 | 2568.63 |
| ETHE1 | 600.38 | 526.5 | 435.9 | 408.34 | 988.52 | 925 | 812.68 | 844.11 |
| SQRDL | 3696.1 | 4049.33 | 4393.75 | 4257.88 | 6131.58 | 5533.78 | 6852.75 | 7215.76 |
| GOT1 | 152.39 | 152.63 | 181.11 | 141.93 | 122.63 | 115.54 | 366.11 | 310.87 |
| GGT1 | 194.72 | 201.97 | 201.58 | 184.97 | 91.71 | 78.2 | 55.82 | 56.73 |
| TST | 1945.05 | 1955.67 | 855.42 | 844.6 | 772.05 | 712.95 | 467.9 | 510.55 |
| CD74 | 745.71 | 813.26 | 473.76 | 437.42 | 426.54 | 419.17 | 147.76 | 201.95 |
| SERPINA1 | 83.95 | 80.94 | 64.46 | 58.17 | 148.22 | 159.22 | 50.89 | 38.57 |
| TGFB1 | 2386.7 | 2275.58 | 1671.96 | 1548.42 | 6333.12 | 5824.74 | 4152.04 | 4073.05 |
| XPO1 | 7199.59 | 7061.85 | 6369.61 | 5794.67 | 4144.95 | 3745.78 | 6219.03 | 6423.84 |
| APOBEC3G | 84.66 | 98.67 | 79.81 | 60.49 | 85.31 | 100.04 | 50.89 | 59 |
| CBX5 | 1276.24 | 1385.23 | 1079.51 | 952.79 | 704.86 | 643.2 | 778.2 | 839.57 |
| PPIA | 788.75 | 764.69 | 268.09 | 295.49 | 294.32 | 322.66 | 49.25 | 72.61 |
| CD4 | 1326.33 | 1436.11 | 1016.07 | 1086.57 | 523.58 | 420.58 | 246.27 | 235.99 |
| CXCR4 | 13544.12 | 13203.29 | 15015.91 | 13298.3 | 689.94 | 672.79 | 466.26 | 621.74 |
| BTRC | 255.39 | 255.15 | 316.18 | 273.39 | 255.93 | 225.44 | 495.81 | 458.36 |
| PTK2B | 1329.86 | 1517.82 | 1342.48 | 1134.27 | 1674.19 | 1270.9 | 3061.9 | 3331.05 |
| FOS | 349.93 | 386.2 | 463.52 | 369.95 | 1903.45 | 1568.2 | 1164.02 | 1064.21 |
| NFKBIA | 1055.42 | 901.91 | 1296.43 | 1012.12 | 2064.48 | 1982.44 | 2700.71 | 2843.19 |
| STAT1 | 3422.36 | 3465.79 | 4003.9 | 3601.74 | 3464.61 | 3107.51 | 5342.32 | 5284.75 |
| STAT3 | 565.1 | 560.41 | 985.37 | 882.99 | 621.69 | 555.14 | 1313.42 | 1377.35 |
| IRF1 | 50.8 | 53.96 | 62.42 | 41.88 | 33.06 | 44.38 | 113.28 | 108.92 |
| YY1 | 2330.96 | 2374.25 | 2490.54 | 2216.19 | 1533.43 | 1330.78 | 2190.12 | 2536.86 |

**Table S5.** Characteristic of ART treated HIV-1 infected study participants.

| Sl. no. | Patient ID | Age | Sex | CD4 cell count<br>(cells/mm<sup>3</sup>) | Viral load (copies/ml) | Date of<br>Diagnosis | Date of start<br>of ART | Data of sample collection |
| --- | --- | --- | --- | --- | --- | --- | --- | --- |
| 1 | Subject A | 32 | F | 555 | 5540.00 | 02/07/15 | 23-03-2015 | 16-08-2016 |
| 2 | Subject B | 31 | M | 484 | <34 | 04-05-2012 | 23/06/15 | 05-10-2016 |
| 3 | Subject C | 34 | F |  | Target not detected | 23-06-2011 | 08/01/15 | 10-08-2016 |
| 4 | Subject D | 25 | F | 642 | Target not detected | 26/09/06 | 16/10/06 | 01/10/19 |
| 5 | Subject E | 40 | M | 351 | Target not detected | 02/01/08 | 01/12/08 | 18/11/19 |

**Table S6.**
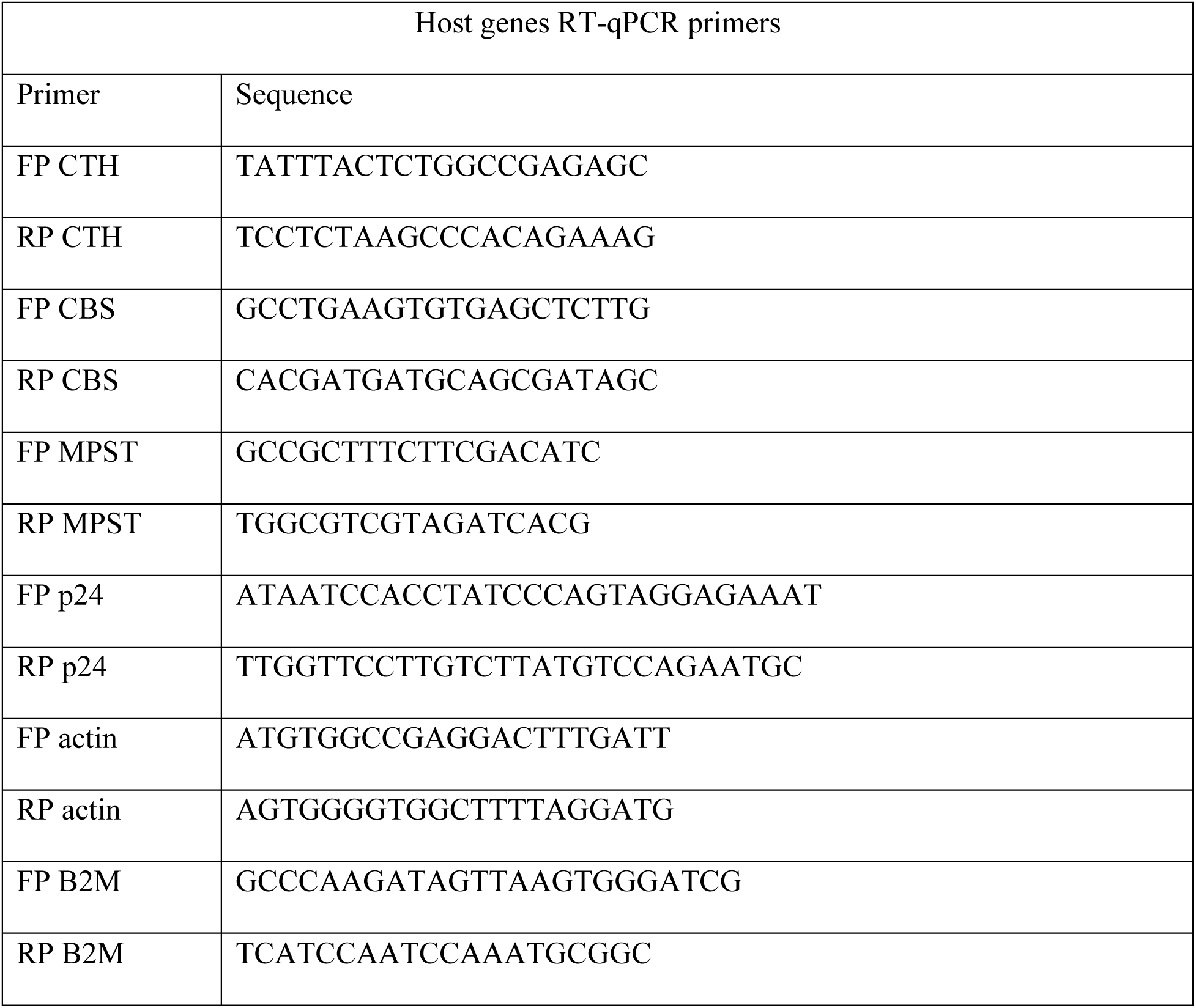
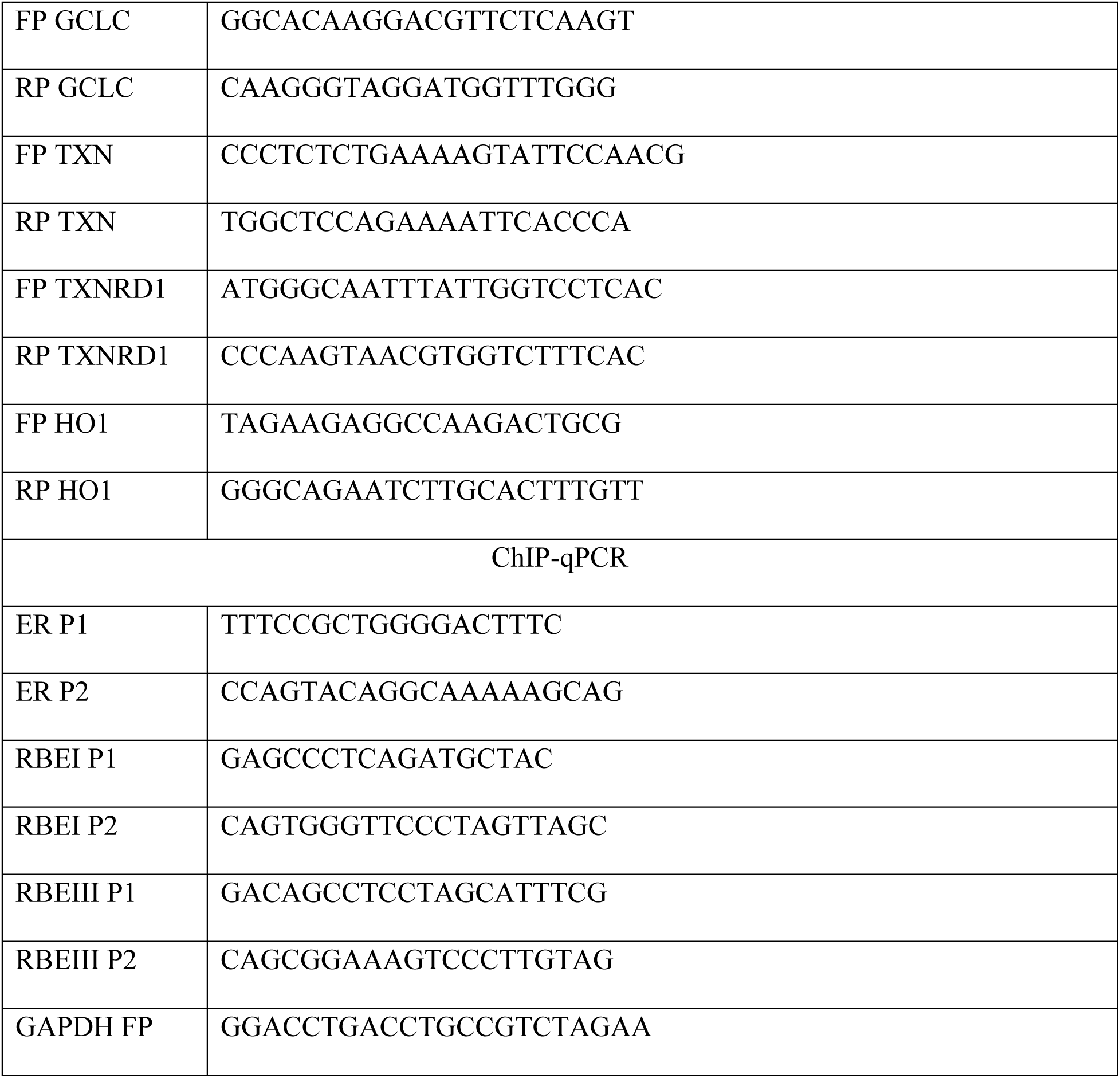
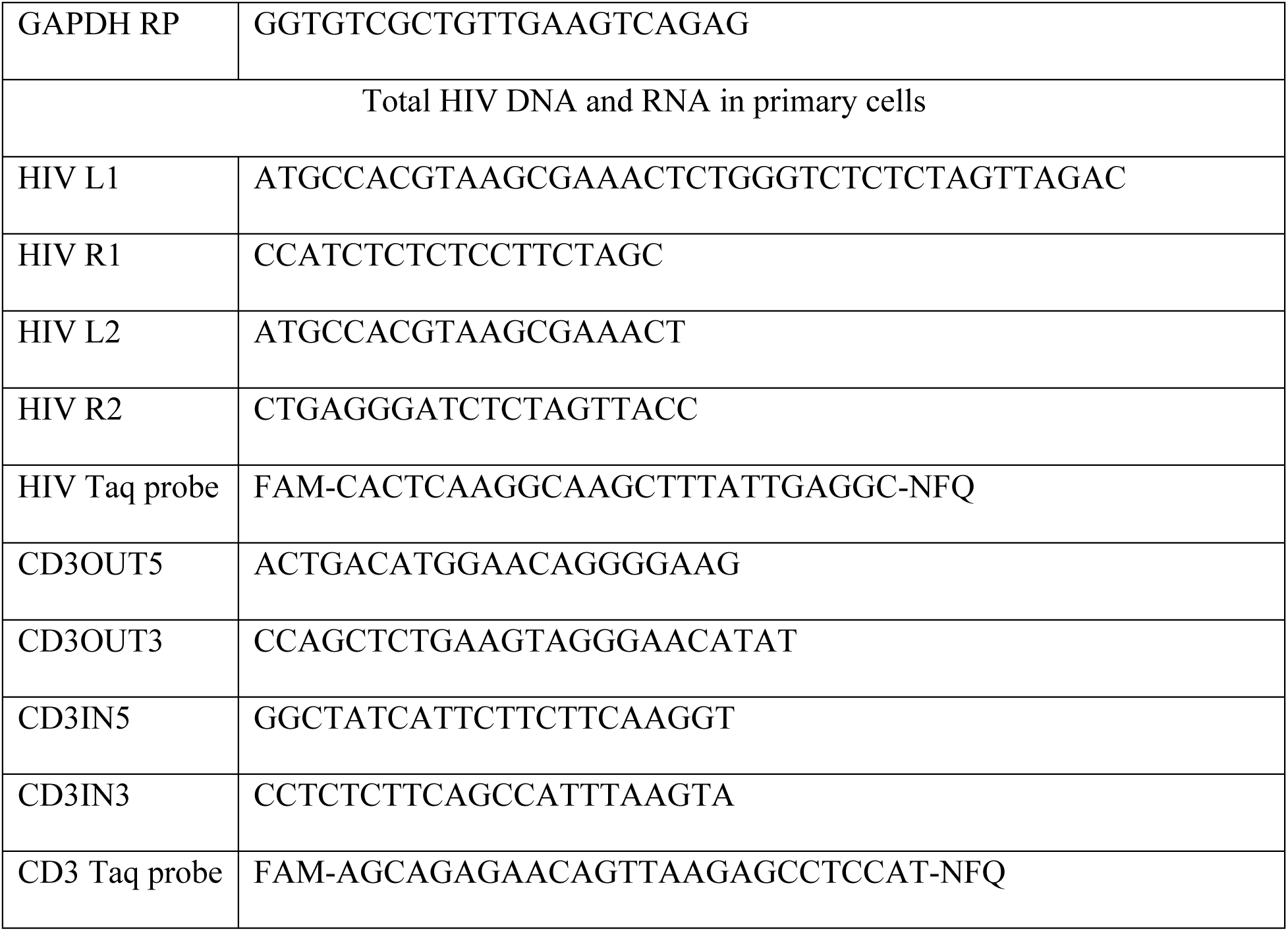
List of primers and probes used in this study.

